# High levels of mitotic gene conversion are needed to effectively purge deleterious mutations in asexual organisms

**DOI:** 10.1101/2024.08.30.610445

**Authors:** Dominik Kopčak, Matthew Hartfield

## Abstract

Self-fertilisation and asexual reproduction are both hypothesised to cause long-term extinction due to inefficient selection against deleterious mutations. Self-fertilisation can counter these effects through creating homozygous genotypes and purging deleterious mutations. Although complete asexuality lacks meiotic gene exchange, mitotic gene conversion creates homozygous regions that could limit deleterious mutation accumulation in an analogous manner. We compare mutation accumulation in self-fertilising and facultative sexual populations subject to mitotic gene conversion, and quantify the efficacy of purging in the latter. We first show analytically that purging is most effective with high levels of asexuality and gene conversion, and when deleterious mutations are recessive. We further show using simulations that, when mitotic gene conversion becomes sufficiently high in obligate asexuals, there is a reduction in the mutation count and a jump in homozygosity, reflecting purging. However, this mechanism is not necessarily as efficient at purging under high self-fertilisation, and elevated rates of mitotic gene conversion seem to be needed for widespread purging compared to empirical estimates. If gene conversion rates are allowed to evolve, then elevated rates that increase mean fitness can arise, but only if there is sufficient variance in the gene conversion rate. Conversely, if gene conversion rates are already high and rates are not constrained then they will slightly decrease, reducing mean fitness.

**Significance Statement:** Asexuality has been argued to be an evolutionary ‘dead end’, due to a lack of gene exchange causing inefficient selection acting against deleterious mutations. It has been proposed that asexuals can counter these negative effects through mitotic gene conversion, which exposes mutations to selection within individual lineages. Here, we theoretically investigate how effective this mechanism is. We compare results to those obtained when individuals reproduce by self-fertilisation, which has similar effects on exposing deleterious variants. While mitotic gene conversion can be effective in removing recessive deleterious mutations, high rates are required (that are not necessarily maintained by selection) and it is not always as effective as selfing.

## Introduction

Mutations are the ultimate source of genetic novelty in all organisms and are essential for biological evolution (Loewe and Hill 2010). However, the generation of variation is a double-edged sword; although favourable mutations may emerge that are crucial for long-term survival, it may also result in a burden of deleterious mutations, which are much more common than favourable ones (Eyre-Walker and Keightley 2007).

The reproductive system, which determines how offspring are generated in a population, has a profound impact on the prevalence of deleterious variation. Reproduction can either be with or without sex. Sex usually involves two mechanisms (Hartfield and Keightley 2012): recombination (the exchange of genetic information between homologous DNA sequences, most commonly considered as meiotic crossing-over), and segregation followed by fusion of gametes (specialised cells with halved ploidy and DNA content; Veller et al. 2019). In the simplest case when asexuality is equivalent to clonal reproduction, the genotype of an offspring is identical to that of a parent, except for *de novo* mutations. Clonal reproduction encompasses diverse mechanisms, including fission or budding in unicellular eukaryotes; budding (e.g., hydra, jellyfishes), fragmentation (e.g., corals) and whole-body fission (e.g., flatworms, starfish) in animals; or many forms of vegetative propagation in plants (De Meeûs et al. 2007). Nevertheless, most eukaryotic organisms undergo some variant of sexual reproduction, at least occasionally (Speijer et al. 2015). Facultatively sexual organisms reproduce via a mixture of sexual and asexual reproduction, so that genetic evolution is affected by the frequency of both recombination and clonal reproduction (Hartfield 2016).

Gametes from which offspring originate may also be derived from the same individual, leading to self-fertilisation or selfing. The opposite of selfing is panmictic outcrossing, with anyone in the population having an equal chance of being a mate of anyone else, and self-mating being rare or even prohibited (if, e.g., self-incompatibility mechanisms are present; Nettancourt 2001). Cloning and selfing are two types of uniparental reproduction, which forms a basis of their similarity (e.g., the lack of mate finding) and potential ecological advantages (e.g., reproductive assurance; Baker 1955; Baker 1967; Theologidis et al. 2014; Mráz and Mrázová 2021).

Despite these apparent similarities, cloning and selfing differ in the manner in which they affect selection acting against deleterious mutations. The operation of ‘Muller’s Ratchet’ is a proposed feature of asexual populations that leads to stochastic mutation accumulation, due to their inability to generate offspring that have fewer deleterious alleles than those of their ancestors (Muller 1964; Haigh 1978). This mechanism is a form of selection interference, which refers to the effect of strong genetic linkage on the dynamics of variants under selection (Felsenstein 1974). In completely clonal populations all sites are in absolute genetic linkage, therefore the presence of multiple deleterious alleles selectively interferes with one another having a strong impact on mutation accumulation (Hill and Robertson 1966; Roze and Barton 2006; Hartfield and Keightley 2012). Recombination breaks this linkage and allows the dynamics of variants to respond to selection independently, i.e., more efficient removal of deleterious mutations and restoration of the most-fit genotype. Another important consequence due to the lack of gamete fusion in diploid asexuals is the divergence of homologous chromosomes found within one individual. The two genome copies are always transmitted together down the lineage, gradually and independently accumulating mutations leading to a high level of heterozygosity; a process known as ‘allelic sequence divergence’ or the Meselson effect (Mark Welch and Meselson 2000; Butlin 2002; Hartfield 2016).

In contrast to cloning, the main genetic consequence of selfing is genome homogenization. Under complete selfing, parents with heterozygous genotypes *Aa* have a 50% probability of losing heterozygosity, independent of the allele frequencies in the rest of the population. The probability that heterozygosity is retained decreases exponentially every generation, rapidly leading to overall genome homogenisation. This homogenisation also occurs, albeit at a lower rate, under partial selfing (Wright 1951). As a consequence, the effective recombination rate is reduced since recombination creates variability only when heterozygosity is present (Nordborg 2000). An additional consequence of increased homozygosity is the exposure of recessive deleterious alleles to selection, giving rise to inbreeding depression (Charlesworth and Willis 2009). An outcome of selfing broadly shared with cloning is the removal of neutral variation through extensive linked selection acting against deleterious mutations, reducing the effective population size (Agrawal and Hartfield 2016).

Both reproductive systems have been called ‘dead-end’ strategies, meaning that species using them have an increased extinction probability. In asexual and selfing populations, this phenomenon has often been ascribed to mutational meltdown, i.e., the accumulation of deleterious mutations leading to extinction (Lynch et al. 1993; Lynch et al. 1995; Olofsson et al. 2023), but note that other mechanisms have been proposed for the phylogenetically short-lived appearance of asexual species (Engelstädter 2008; Janko et al. 2008; Schwander and Crespi 2009). While the ‘dead-end’ hypothesis has also been invoked for selfing species (Wright et al. 2013), they can increase their fitness over time through purging deleterious mutations (Glémin 2003; Abu Awad and Billiard 2017); this purging can also stop Muller’s ratchet from operating (Charlesworth et al. 1993). Furthermore, there exist ancient asexuals that have seemingly existed for long periods of time in the absence of sexual reproduction, including species of bdelloid rotifers (Flot et al. 2013), oribatid mites (Brandt et al. 2021), ostracods (Tran Van et al. 2021), and trypanosomes (Weir et al. 2016). There is some debate as to whether these species are truly ancient asexuals; e.g., Vakhrusheva et al. (2020) argued that the bdelloid rotifer *Adineta vaga* show evidence of inter-individual gene exchange, and rare males of the Oribatid mite *Oppiella nova* are observed, although they appear to be sterile (Wehner et al. 2018). Still, it is apparent that sex is sufficiently rare in these species to beg the question of whether there are underexplored mechanisms at play that prevent them from undergoing excessive accumulation of deleterious variants, similar to what occurs in self-fertilising species.

Although genome exchange is an intrinsic part of sexual reproduction, gene conversion is a mechanism of genome exchange that acts in both sexuals and asexuals in the form of mitotic recombination (Lee et al. 2009). These gene conversion events can result in the creation of homozygous tracts, either over localised regions or over large parts of chromosomes depending on the underlying mechanism (Smukowski Heil 2023). In addition, it is becoming increasingly known that many asexual species modify their meiosis cycle when reproducing, which also results in increased runs of homozygosity (e.g., in the bdelloid rotifer *Adineta vaga*, cytological evidence is consistent with meiosis I being aborted, leading to runs of homozygosity in offspring; Blanc et al. 2025). This creation of homozygosity is similar to what happens under selfing, and in asexual species it has been shown to have an important impact on neutral diversity when sex is rare (Hartfield et al. 2016; Hartfield 2021). Mitotic gene conversion has been argued to affect deleterious mutation accumulation in asexual species, including *Adineta vaga* (Flot et al. 2013), the crustacean *Daphnia pulex* (Tucker et al. 2013), and the planarian worm *Schmidtea mediterranea* (Kershenbaum et al. 2023). Khakhlova and Bock (2006) also show experimental evidence of gene conversion eliminating deleterious mutations in plastids of tobacco plants, which are uniparentally inherited and are effectively asexual. Most of these studies argue that this gene conversion is beneficial for asexual species by removing deleterious mutations (an exception being Tucker et al. (2013) that argued that mitotic gene conversion reduces fitness by creating homozygous mutations). However, most of these arguments are verbal, and there exist little theoretical predictions to determine whether realistic levels of gene conversion can purge deleterious mutations, and hence maintain high fitnesses in asexual populations in an analogous manner to self-fertilisation. One exception is a model in Flot et al. (2013), showing how gene conversion slows down Muller’s ratchet in asexual populations.

The overall goal of this study is to investigate the efficacy of mitotic gene conversion in facultative sexual and asexual species in purging deleterious mutations through creating homozygous regions, and compare its effects to self-fertilisation to determine to what extent these mechanisms of genetic homogenisation are similar. We use two complementary approaches: first, we develop single-locus deterministic analytical models to quantify the frequency of deleterious mutations under facultative sex and gene conversion, and compare results to equivalent models under self-fertilisation. We then use stochastic multi-locus simulations to determine how both reproductive mechanisms influence the genome-wide prevalence of deleterious mutations. Furthermore, we present simulation outputs in a way so that they can be compared with genome data, to enable empirical comparisons with the models presented here.

Our principal questions are: (i) Under facultative sex, how does the frequency of deleterious variants (and hence also the population’s mean fitness) scale with the prevalence of sex and gene conversion, both at individual loci and genome-wide? (ii) What values of sexual reproduction and gene conversion produce a level of mutation purging similar to a given level of self-fertilisation? (iii) Can realistic levels of mitotic gene conversion be as effective as self-fertilisation at purging deleterious mutations? (iv) Are high levels of mitotic gene conversion attained when the rate is allowed to evolve? The findings of this study will provide important quantification of a mechanism for maintaining the fitness of facultative sexuals and asexuals, which has been often discussed but seldom investigated theoretically. It also provides insights as to which mode of uniparental reproduction might be favoured by evolution due to its effects on removing deleterious variation.

### Single-Locus Mathematical Model Results

To gain some insight into how mitotic gene conversion affects the purging of deleterious mutations and how it compares to self-fertilisation, we first investigate a single-locus model of deleterious allele frequencies in a deterministic Wright-Fisher population. We assume populations are diploid, which is the ploidy state for several potential long-term asexuals of interest including oribatid mites (Brandt et al. 2021), planarian worms (Kershenbaum et al. 2023) and some species of nemertean worms (Ament-Velásquez et al. 2016). Individuals are subject to either facultative sex with sexual reproduction occurring at frequency *σ* (i.e., every generation, a fraction *σ* of offspring are generated sexually, and the remaining 1–*σ* are produced asexually), or self-fertilisation at rate *S* (i.e. a fraction *S* of reproductions are produced by selfing). We consider the frequency of a deleterious allele (denoted *A*) that arose from mutation from a wild-type allele *a* with probability *μ* irrespective of the reproductive mode; heterozygous mutant genotypes have fitness 1–*hs* and homozygotes 1–*s*, for *h* the dominance coefficient and *s* the selection coefficient. Mitotic gene conversion occurs with frequency *γ*, which converts heterozygotes to either wild-type or derived homozygote with equal frequency, i.e., the conversion is unbiased. Note that, being mitotic, gene conversion acts under both sexual and asexual reproduction. Mitotic gene conversion events affect all offspring, which is biologically equivalent to gametic asexual reproduction where all offspring are derived from a single ovum (Orive and Krueger-Hadfield 2021). For completeness we will derive equations considering the effects of gene conversion acting in both facultative sexuals and self-fertilising species, although the greater interest will be the effects under infrequent sex.

We can track the prevalence of deleterious mutations by creating numerical recursions of how their frequency is affected assuming the following lifecycle; selection, reproduction, mutation and gene conversion, with non-overlapping generations (following the methods of Agrawal and Hartfield 2016; Hartfield 2021). Although these recursions calculate the change in genotype frequencies over time, we can convert these genotype frequencies to allele frequencies; specifically, if the frequency of the mutant allele is *p_A_*, then the frequency of the mutant homozygote is 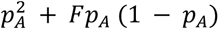 (with similar equations for the wild-type homozygote), and the heterozygote has frequency 2 *p_A_* (1 − *p_A_*) (1 − *F*). Here, *F* denotes the reduction in homozygosity compared to Hardy-Weinberg equilibrium (Wright 1951), and shows to what extent either gene conversion under facultative sex or self-fertilisation affects genotype composition by reducing heterozygote frequencies. After making these transformations, we linearise the equations assuming *μ*, *p_A_* ≪ 1 to obtain steady-state values of *p_A_* and *F* (see Otto 2003 for a similar approach). Appendix 1 in the Supplementary Text and the Supplementary *Mathematica* file outlines the derivations and full recursion equations used to obtain analytical solutions.

After these algebraic calculations, we obtain the general equation for the deleterious allele frequency:

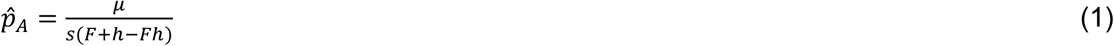

This is a known form for mutation-selection balance under non-random mating (Ohta and Cockerham 1974; Glémin 2003). The inbreeding coefficient *F* in Equation 1 differs depending on the form of uniparental reproduction. Under facultative sex:

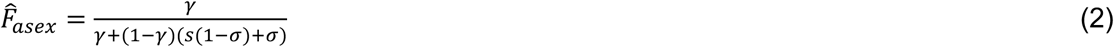

While under self-fertilisation:

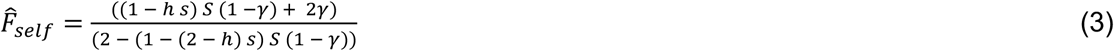

We can use heuristic arguments to compare *F* in each case to understand which reproductive mode has the greatest effect on homozygosity. Focussing on the effects of selfing (i.e., setting gene conversion and selection to zero), then *F̂_self_* ≈ *S*/(2–*S*) in line with classic results on how selfing affects homozygosity (Nordborg and Donnelly 1997). Under facultative sex (Equation 2), positive *F̂_asex_* is only formed when gene conversion is present; interestingly, dominance does not appear in Equation 2 and instead affects allele frequencies through the mutation-selection balance term (Equation 1). Assuming selection is absent and *γ* << *σ* so the denominator approximates to *σ*, then 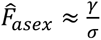 (as also found by Hartfield 2021). By comparing these two approximations, we can show that the inbreeding values are equivalent if S = 2/(1+*φ*) for *φ* = *σ*/*γ* (Hartfield et al. 2016; Hartfield 2021). As sex is much more common than gene conversion under these assumptions (i.e., *φ* >> 1) then we see that gene conversion acting under frequent sex is only equivalent to a small selfing rate. Note that this approximation for *S* will break down once gene conversion and sex act at the same rate. Here, gene conversion becomes more effective at making homozygous genotypes once sex becomes rarer due to the enforced pairing of diploid gene copies, which are subsequently maintained by asexual reproduction (Hartfield et al. 2016). The genetic outcome then becomes more similar to that expected under higher levels of self-fertilisation.

Figure 1 illustrates these derived equations for a single locus and different values of the rate of sex and mitotic gene conversion. If mutations are extremely recessive (*h* = 0.01, Figure 1a) then even small amounts of gene conversion can reduce the prevalence of deleterious mutations compared to outcrossing expectations, even for intermediate frequencies of sex. This purging effect is weakened as the dominance coefficient increases, so that the frequency of additive alleles becomes closer to outcrossing expectations (Figure 1b, c). These results illustrate that mitotic gene conversion has the strongest effect on purging deleterious mutations if they are highly recessive, with the purging effect magnified as sex becomes rarer. Under self-fertilisation (Figures 1d–f), the same general results arise, but we note that only a small amount of selfing is needed to remove highly recessive deleterious mutations, reflecting its stronger effect on creating homozygosity. Mitotic gene conversion also only has a minor effect on allele frequencies (note that the contour lines are nearly vertical).

**Figure 1:**
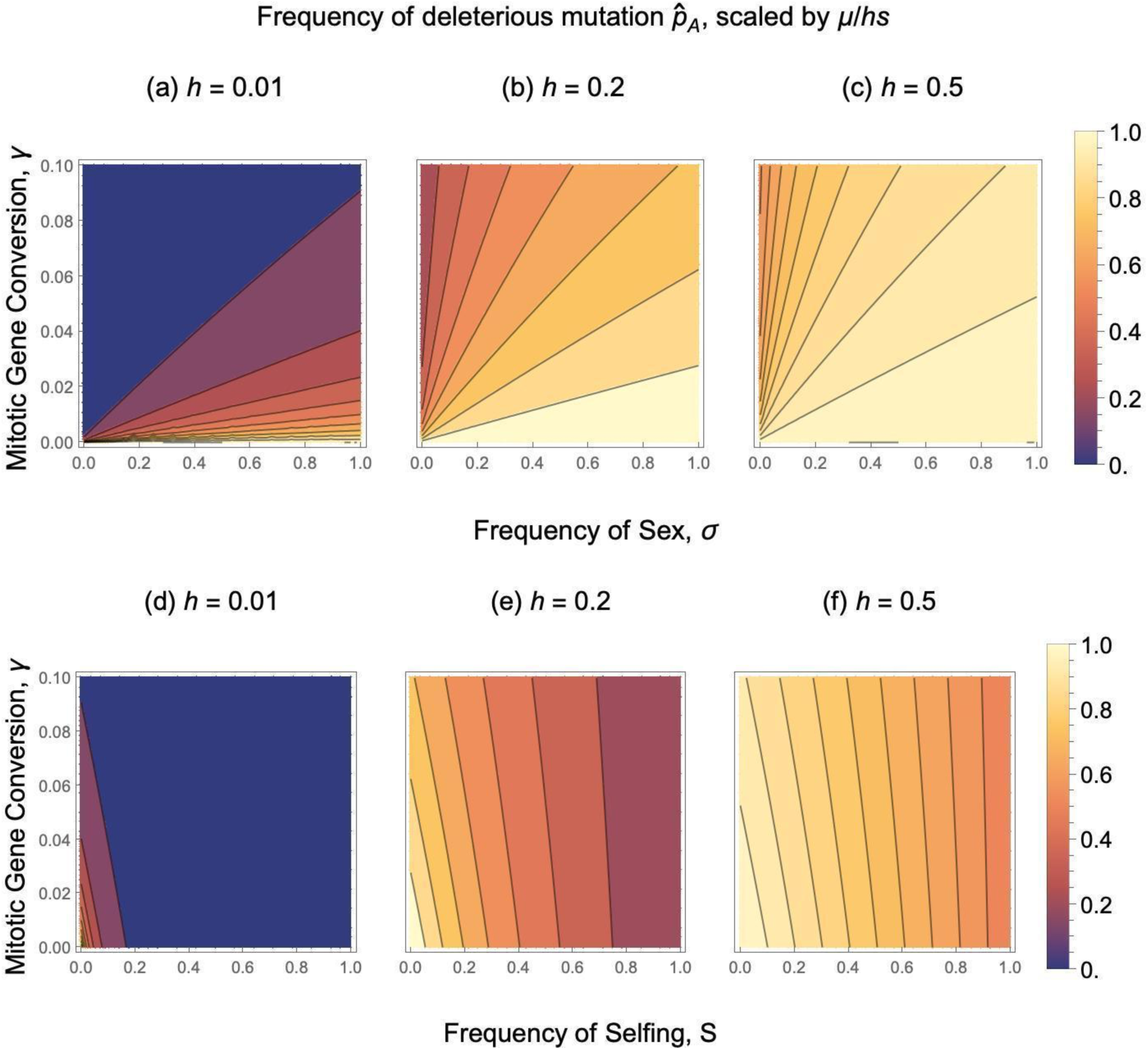
Plots of single-locus analytical results for the steady-state frequency of a deleterious mutation. Results show Equation 1 for different dominance values, scaled to the expected result under outcrossing (μ/hs). Note that μ is not present in this scaled equation. Panels (a) – (c) are for different frequencies of sex (with F̂ given by Equation 2), while (d) – (f) are for different frequencies of self-fertilisation (with F̂ given by Equation 3). **Alt text:** Contour plots showing deleterious mutation frequency with subfigures labelled a to f, illustrating effects of reproductive modes and mitotic gene conversion.

Plots of *F̂* (Figure 2a, b) show that while mitotic gene conversion under facultative sex leads to elevated homozygosity, the increase is mostly modest unless sex is rare and deleterious mutations have a weak selection effect. In contrast, under self-fertilisation then *F̂* values tend to be much higher (Figure 2c, d).

**Figure 2:**
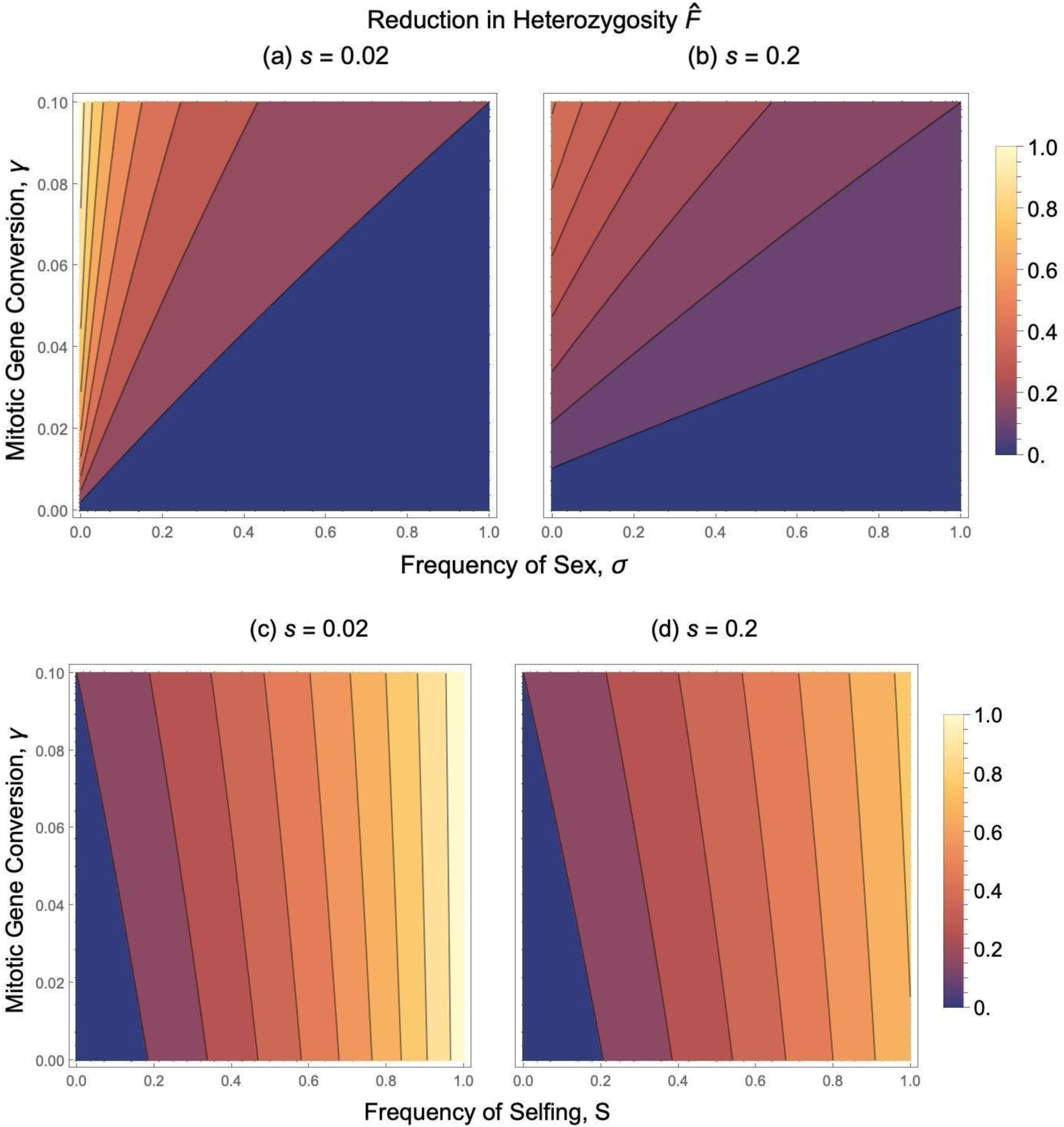
Plots of single-locus analytical results for the reduction in heterozygosity, F̂. Results show either Equation 2 under facultative sex (panels a, b), or Equation 3 under self-fertilisation (panels c, d). **Alt text:** Contour plots showing reduction in heterozygosity with subfigures labelled a to d, illustrating effects of reproductive modes and mitotic gene conversion.

To test the accuracy of these approximate analytical solutions (Equations 1-3), we compared them to exact values obtained from numerical recursions over 50,000 generations, where the deleterious heterozygous had an initial frequency of 0.001 (Supplementary Figures S1 and S2, with code in Supplementary *Mathematica* file). We see that the two match over a wide range of parameters. One caveat of these models is that they do not consider drift, and hence processes like Muller’s ratchet (Muller 1964; Haigh 1978; Flot et al. 2013) that can arise when sex is rare. That said, both our model and that of Flot et al. (2013) come to the same general conclusion; when mitotic gene conversion is present in asexuals, it creates homozygote genotypes leading to a lower prevalence of deleterious mutations.

### Simulation Results

While the single-locus model provides insight into how rare sex and gene conversion shape deleterious mutation frequencies, it does not show how gene-conversion events that affect mutations at multiple loci purge mutations across larger, linked stretches of the genome. Hence, we next use simulations written in SLiM (Haller and Messer 2023) to compare (i) mutation accumulation in populations undergoing different levels of either self-fertilisation, or asexual reproduction; and (ii) how gene conversion leads to mutation removal in obligate asexuals, compared to selfing species. We also investigate how the mitotic gene conversion rate can evolve. Note that we do not consider gene conversion in selfing populations, as the single-locus results demonstrate that it only has a minor effect under this reproductive mode. To achieve this goal, we had to write a new SLiM routine to consider gene conversion under asexual reproduction. More information is available in the “Multi-Locus Simulation Methods” section.

### Comparing self-fertilisation and facultative sex

We first compared the mean fitness and mutation counts under different levels of self-fertilisation or asexual reproduction, to determine how mutation accumulation differs between the two reproductive modes. All results in the main text assume a fixed *s* and *h*; we will present results with varying *s*, *h* values at the end of the results.

Figure 3 shows the final mean fitness over different rates of uniparental reproduction (see Supplementary Figure S3 for fitness change over time). Under self-fertilisation, while the mean fitness is fairly high overall, it is lowest when there is complete outcrossing. It generally increases with the selfing rate, reaching a maximum for around 95% self-fertilisation, in line with deleterious mutations being purged. However, fitness then drops for extremely high rates, likely because of a large drop in the effective population size so genetic drift more strongly affects deleterious mutations, allowing them to persist (Roze 2016). In contrast, the mean fitness drops dramatically for highly asexual populations.

**Figure 3:**
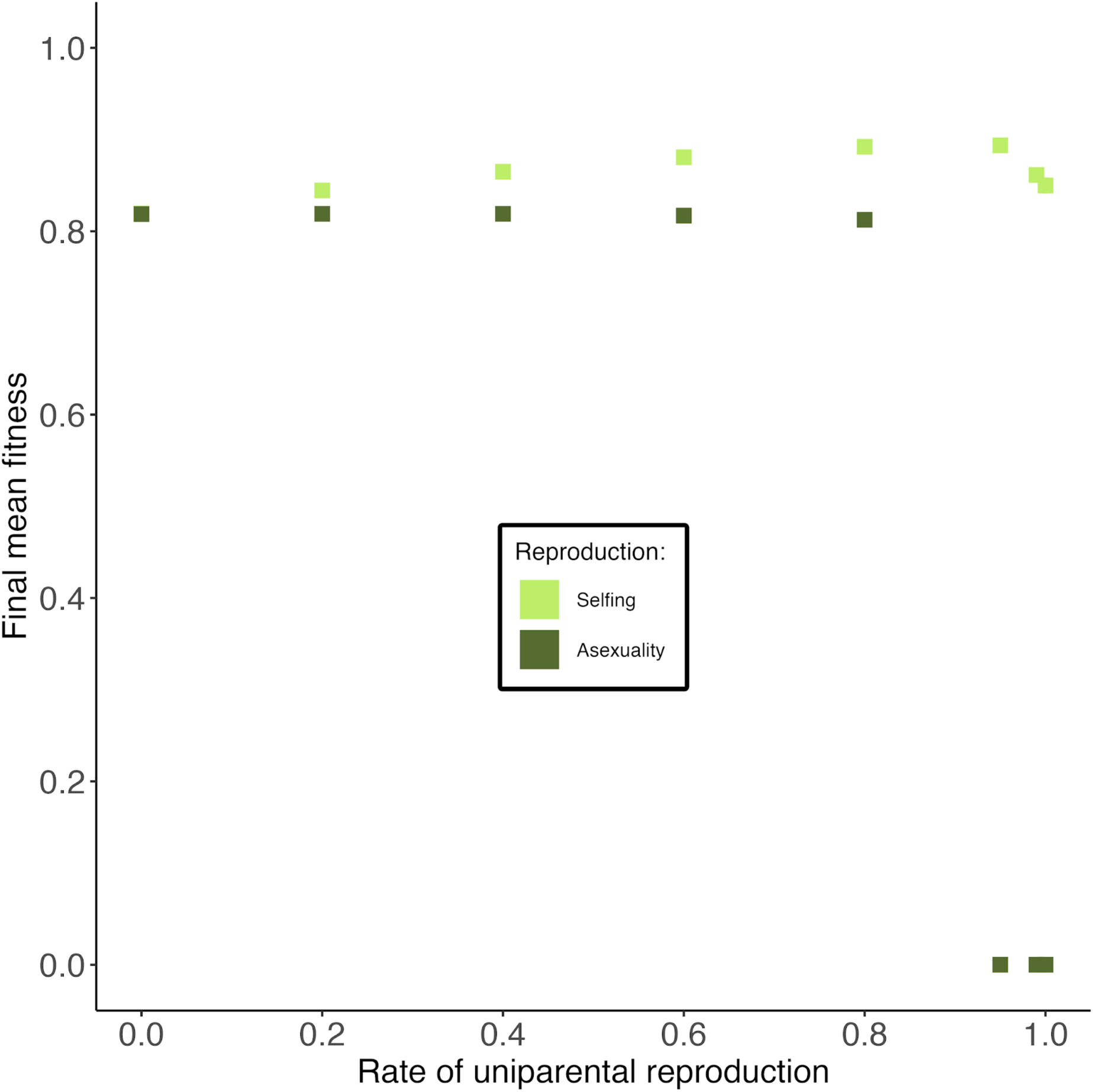
Mean fitness at the end of simulations for different modes of uniparental reproduction. Error bars represent 95% confidence intervals; if they are not shown then they lie within a point. **Alt text:** Graphs showing mean fitness, with different points indicating different rates of either self-fertilisation or partial asexual reproduction.

Figure 4 (and Supplementary Figure S4) shows mutation counts under these reproductive regimes. Selfing populations undergo mutation purging, as evidenced by a decreasing number of mutations in the population (approximately linearly with the selfing rate) and an increase in homozygous mutations reflecting higher inbreeding. In contrast, asexual populations have approximately the same mutation counts (both total and homozygote) as outcrossing populations, until the population becomes around 95% asexual. At this point, there is a spike in homozygosity that is then reduced with even higher levels of asexuality, while the total mutation count rapidly increases. The high mutation count reflects inefficient selection acting once asexuality becomes prevalent, as reflected by the drop in mean fitness (Figure 3). The spike in homozygosity likely reflects the fact that for high-but-incomplete asexuality, mutation accumulation starts acting but there is still sufficient segregation to create homozygous genotypes. In populations that become completely asexual (i.e., the level of clonal reproduction approaches 1), there is a lack of genetic segregation so genotypes only form as heterozygotes (the signature of ‘allelic sequence divergence’). The relative homozygosity (i.e., the proportion of mutations in the homozygous state; see Supplementary Figure S5 and the Methods for a formal definition) also reflects these outcomes; it increases towards 1 with higher selfing, reflecting greater homozygosity due to inbreeding. In contrast, it stays constant with increased asexuality up to 95% asexual reproduction, but then decreases with higher rates. No apparent spike at 95% asexuality is observed using this measure.

**Figure 4:**
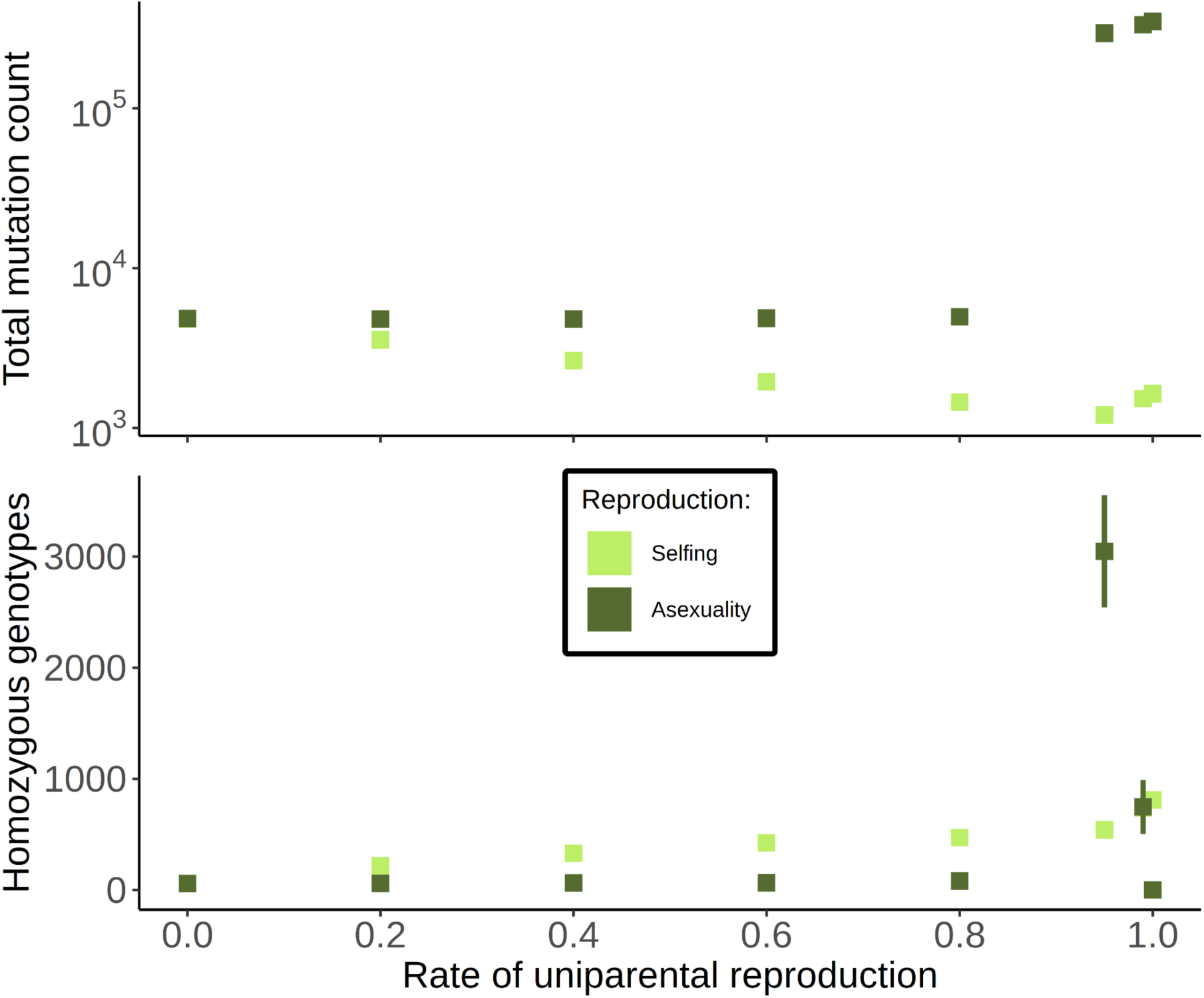
Deleterious mutation counts under different uniparental reproductive modes. As a function of the rate of uniparental reproduction (either selfing or asexual reproduction), these plots show (top) the total mutation count present in the sample of 50 individuals, and (bottom) the number of homozygous genotypes present in the sample. Error bars represent 95% confidence intervals; if they are not shown then they lie within a point. See Supplementary Figure S4 for the total count on a linear scale. **Alt text:** Graphs showing total mutation count and number of homozygote genotypes, with different points indicating different rates of either self-fertilisation or partial asexual reproduction.

### Impact of mitotic gene conversion

Given this increased mutation count under low rates of sex, we next investigated to what extent mitotic gene conversion can reduce it. Here, we used the non-WF model with the new gene conversion routine implemented. Figure 5 plots the final mean fitness for different per-base-pair initiation rates of gene conversion (with results over time shown in Supplementary Figure S6). For most values the mean fitness remains low, similar to results in the absence of gene conversion (Figure 3b). However, once mean gene conversion per site becomes sufficiently high (in this case, 4x10^-4^ and above) the mean fitness increases and starts reflecting that of outcrossing populations. These results imply that purging is indeed occurring. Purging is further validated when looking at mutation counts (Figure 6). Here, the mutation count drops while the homozygosity increases (as reflected with heightened relative homozygosity; Supplementary Figure S7), in line with purging reducing the total mutation prevalence and increasing the mean fitness.

**Figure 5:**
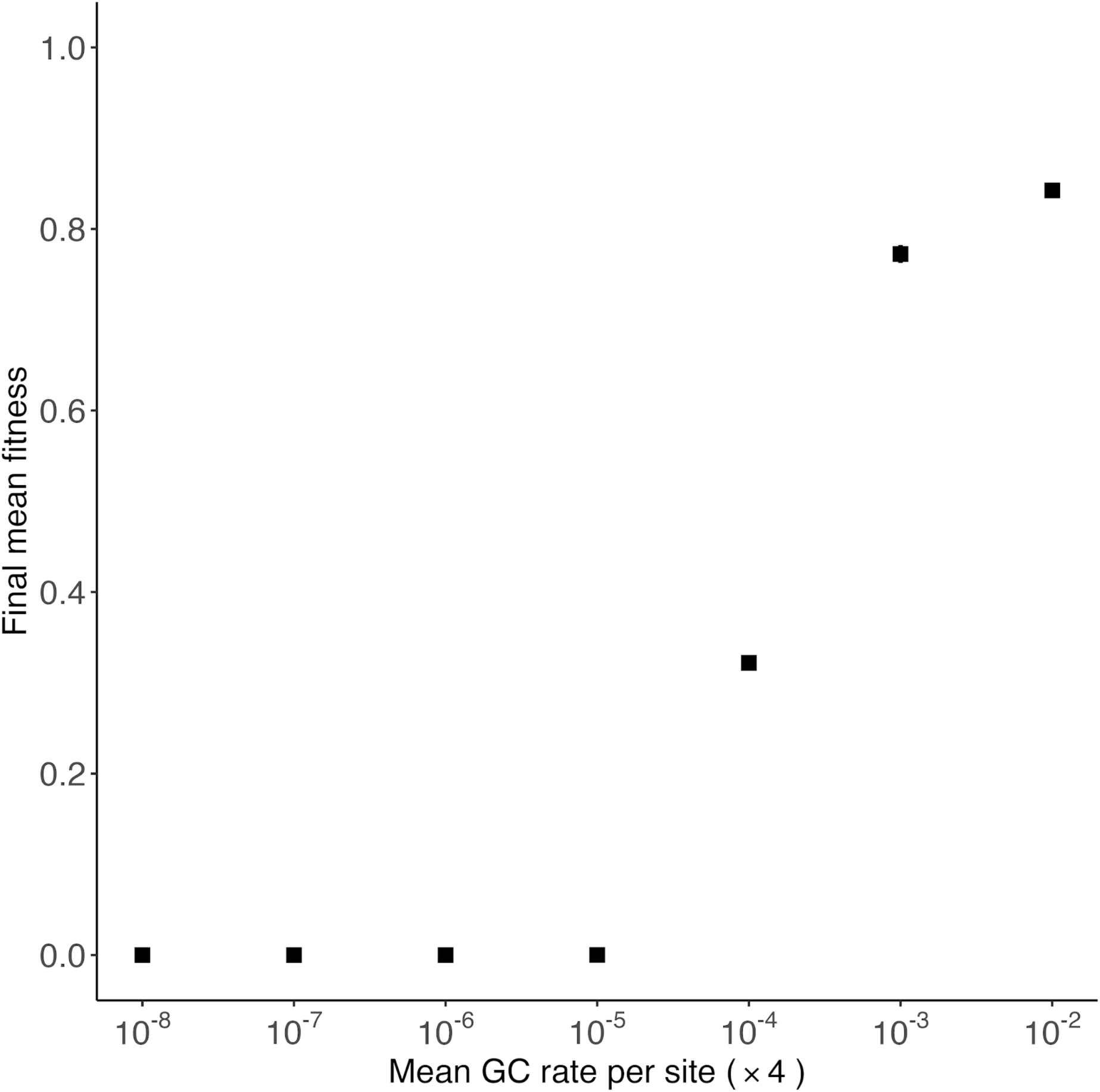
Mean fitness at the end of simulations for obligately asexual populations when mitotic gene conversion is present at different rates. Error bars for simulation results represent 95% confidence intervals; if they are not shown then they lie within a point. **Alt text:** Graphs showing mean fitness under complete asexuality, with different points indicating different rates of mitotic gene conversion.

**Figure 6:**
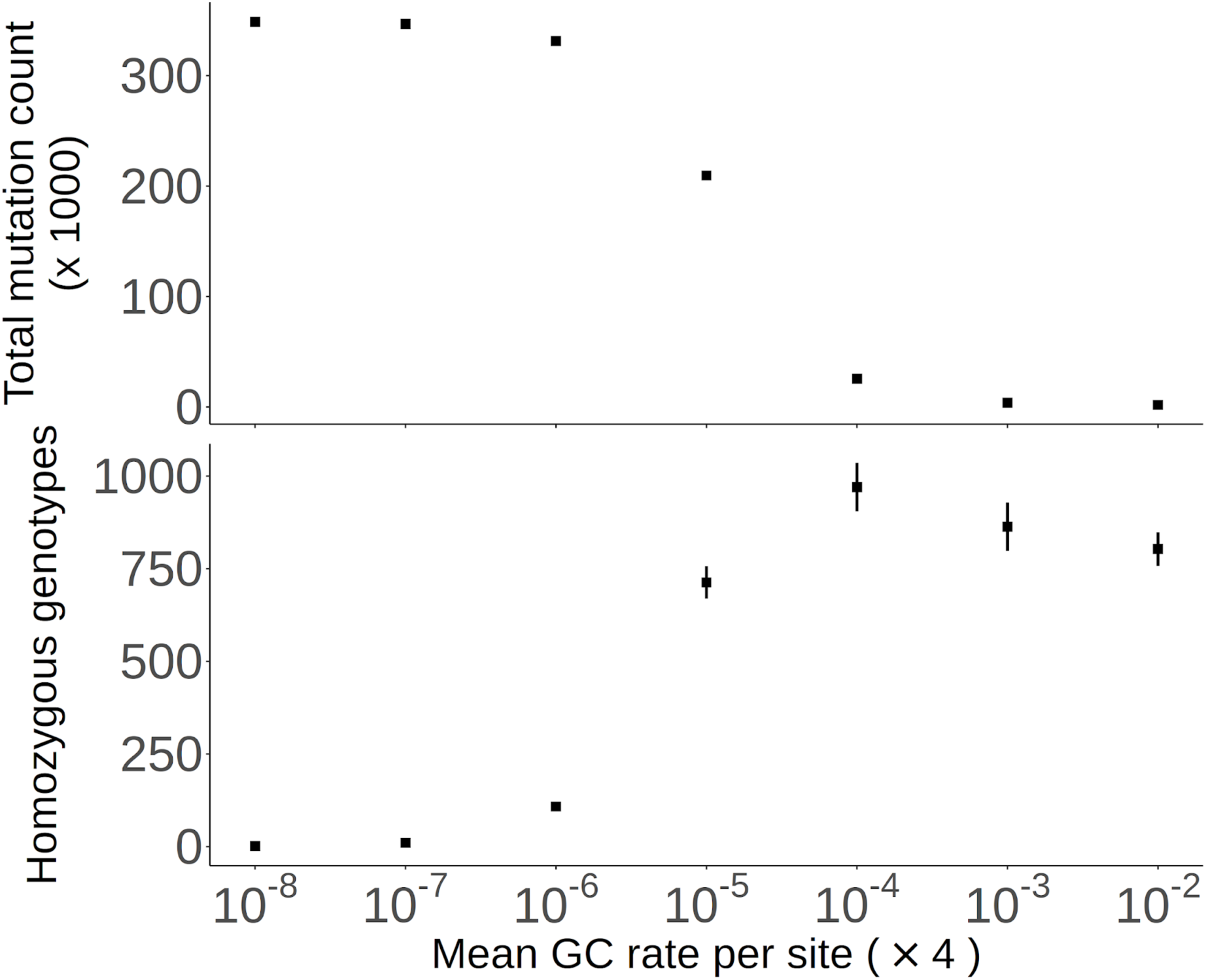
Deleterious mutation counts for obligate asexual populations with mitotic gene conversion present. Results are shown for simulations, as a function of the rate of gene conversion initiation at a single site. Plots show (top) the total mutation count present in the sample of 50 individuals, and (bottom) the number of homozygous genotypes present in the sample. Error bars for simulation results represent 95% confidence intervals; if they are not shown then they lie within a point. **Alt text:** Graph showing total mutation count and number of homozygote genotypes from simulations, with different points indicating different rates of gene conversion under complete asexual reproduction.

### Comparing mitotic gene conversion to self-fertilisation

We next compare the effects of mitotic gene conversion in asexuals to self-fertilisation, to determine to what extent the two effects are comparable. Figure 7 compares gene conversion in asexuals against a range of selfing rates. When looking at homozygosity, it seems that asexuality with the highest gene conversion rates (4x10^-2^) yields the same homozygous counts as complete selfing, implying that the two are equivalent. However, the total mutation count is lower for high levels of self-fertilisation (80% and above), compared to the highest gene conversion rate used. Hence, it seems that while gene conversion in asexuals can purge mutations, it does not produce the same benefits in terms of reducing deleterious mutation prevalence as observed under high self-fertilisation.

**Figure 7:**
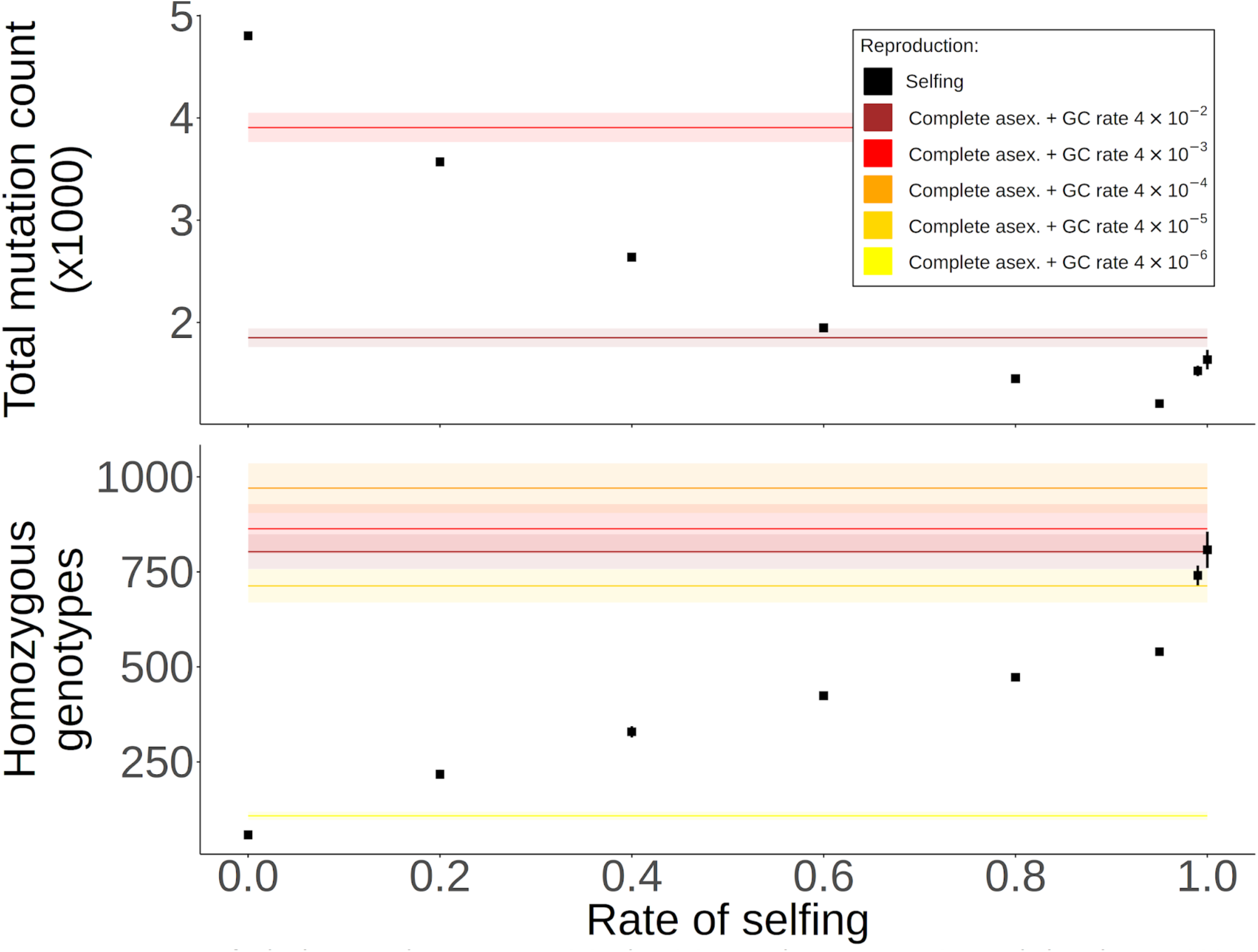
Comparing mutation counts under different levels of self-fertilisation to asexual populations subject to mitotic gene conversion. Plots show (top) the total mutation count present in the sample of 50 individuals, and (bottom) the number of homozygous genotypes present in the sample. Note that only gene conversion rates of 4x10^-2^ and 4x10^-3^ are shown in the top panel. The points show results under different levels of self-fertilisation; error bars represent 95% confidence intervals, and if they are not shown then they lie within a point. Coloured lines are results for asexuality with different levels of mitotic gene conversion, as indicated by the legend; shaded bands around lines indicate 95% confidence intervals. **Alt text:** Graph comparing total mutation count and number of homozygote genotypes, with points indicating different selfing rates, and horizontal lines are results for complete asexuality with different gene conversion rates.

### Exploring results under different parameters

#### Results with different population sizes and rescaled parameters

To test if these simulation results hold more generally, we next explored deleterious mutation prevalence under different parameters. Under classic population-genetic models, evolutionary outcomes are not just dependent on the mutation rate and selection coefficients, but on their products with the population size (Hill and Robertson 1966; Kim and Wiehe 2009; Uricchio and Hernandez 2014; but see Dabi and Schrider 2025; Johri et al. 2026; Marsh et al. 2026 for discussions into when such rescaling breaks down). It has been previously shown that the effects of mitotic gene conversion on increased homozygosity is dependent on *Nγ* if the rate of gene conversion is sufficiently rare (Hartfield et al. 2016; Hartfield et al. 2018; Hartfield 2021). Hence, if this scaling influences mutation purging, then we should see lower mutation counts arising for smaller gene conversion values if other compound parameters are kept constant. We therefore ran new simulations with different population sizes (*N* = 1,000 and 10,000 respectively) but with the mutation rate and selection coefficients scaled so that *Nμ*, *Ns* are the same as for *N* = 5,000 case.

Results for mutation counts are shown in Supplementary Figures S8, S9. It seems that the same general levels of gene conversion (i.e., 4x10^-4^ and above) lead to the largest drops in deleterious mutation count. However, for *N* = 10,000 greatly increased homozygosity is observed for 4x10^-5^, albeit with only a modest reduction in total mutation count. This result hints that purging does depend on the compound parameter *Nγ*. Hence, it seems that the observed results for *N* = 5,000 still generally hold if other compound parameters (*Nμ*, *Ns*) are kept constant, but there are indications that gene conversion can be more effective in much larger populations due to more gene conversion events in a population (larger *Nγ*).

#### Results where *s*, *h* are drawn from distributions

We also investigate simulations where *s*, *h* are drawn from probability distributions, using the same mutation rate as before. Specifically, selection coefficients are drawn from a Gamma distribution, with dominance values drawn so that strongly-selected mutations are more recessive, in line with empirical observations (see Methods for further details). Mean fitness results are generally similar to the fixed *s*, *h* case although, if looking just at selfing populations, the highest fitness is realised for complete self-fertilisation (Supplementary Figure S10). Mutation accumulation in the absence of gene conversion acts slightly differently; while the highest mutation count occurs under high levels of sexual reproduction, the number of homozygous genotypes is reduced under both high levels of selfing and asexual reproduction (Supplementary Figures S11, S12). The relative homozygosity is higher than that for fixed *s*, *h* cases under low rates of sex, but drastically drops once the rate of asexuality reaches 95% and higher (Supplementary Figure S13). Gene conversion in asexuals restores population fitness in a similar manner to the fixed *s, h* simulations (Supplementary Figure S14), with slightly lower levels of gene conversion needed to purge deleterious mutations (Supplementary Figures S15 – S17). Hence, it seems that the main results do not strongly depend on the fitness model used.

### Results with evolving gene conversion rates

In these simulations, rather than gene conversion being fixed over time, then offspring instead inherit the parental gene conversion rate plus an additional value so it evolves over time. Added values are drawn from a normal distribution with mean zero and a pre-defined standard deviation (so the rate can also decrease). The size of the standard deviation determines how ‘evolvable’ gene conversion rates are. This model mimics a scenario where gene conversion is a complex trait affected by multiple loci. More information on this model is available in the Methods.

Figure 8 shows the mean fitness under evolving gene conversion, and also how the gene conversion rate changes over time. If the standard deviation of evolving gene conversion rates are low (1x10^-8^) so values are constrained, then fitness greatly drops unless the initial gene conversion rate is high (4x10^-2^), in which case fitness also remains rather high. With higher gene conversion evolvability (standard deviation of 1x10^-7^) the fitness converges to the same values of just under a half, irrespective of the starting rate. This outcome is an improvement when the starting gene conversion rate is low (4x10^-7^ or 4x10^-4^), but detrimental if it is high (4x10^-2^). For the most part, gene conversion rates tend to reach a steady-state frequency very quickly, which is usually high (between 1x10^-4^ and 1x10^-1^). The main outlier case is when the starting gene conversion rate is high (4x10^-2^) and can more freely evolve (standard deviation of 1x10^-7^). Here the mean gene conversion rate slowly decreases over time, but there is substantial variability around mean values.

**Figure 8:**
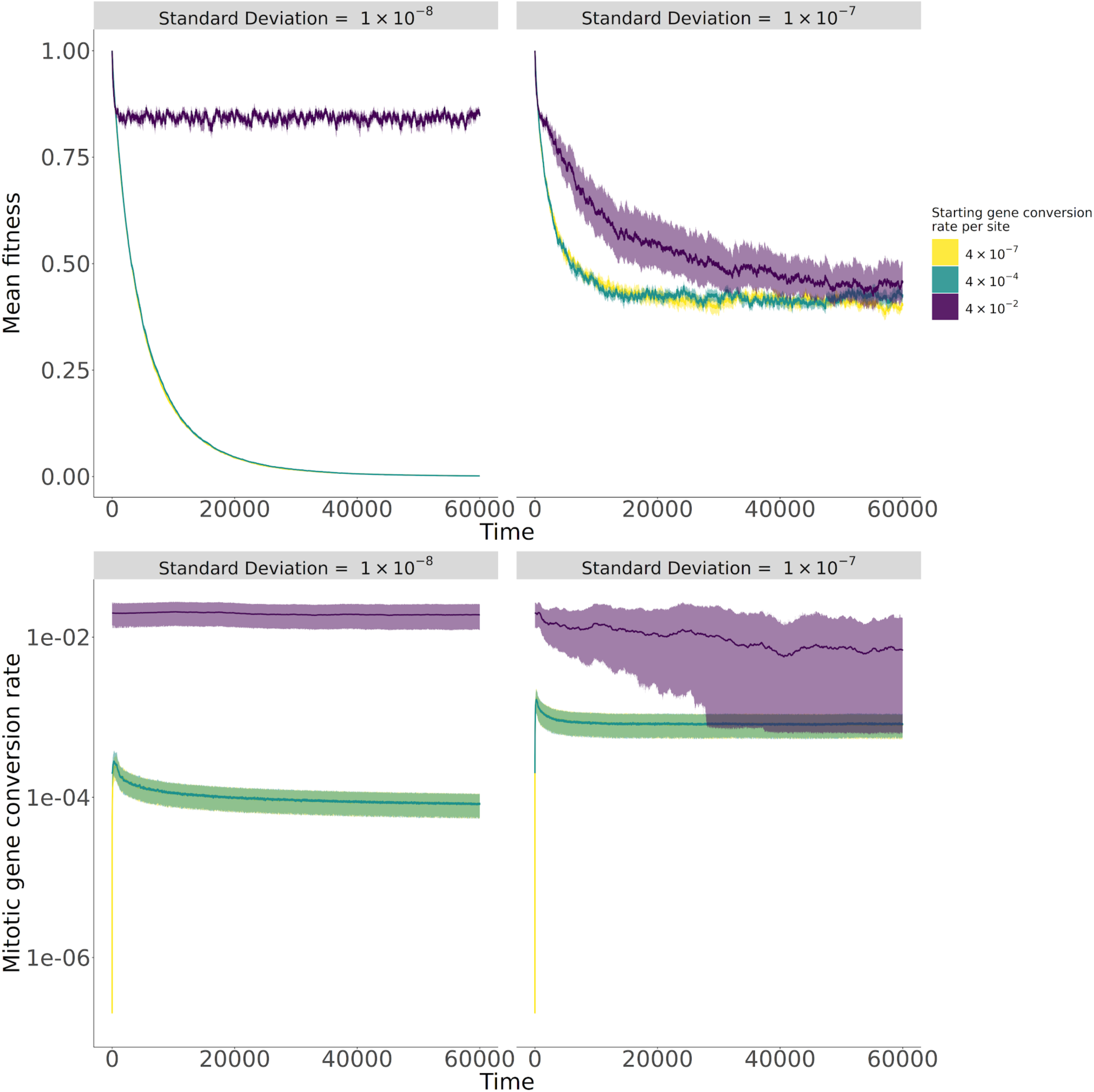
Mean fitness (top) and mitotic gene conversion rate (bottom) over time, when the rate is allowed to evolve. Lines are mean values, while bands around lines represent 95% confidence intervals. **Alt text:** Graphs showing mean fitness and mitotic gene conversion rate when the latter can evolve, with lines representing mean values and ribbons are 95% confidence intervals.

The mutation counts reflect these variable outcomes of evolving gene conversion (Figure 9, with relative homozygosity plotted in Supplementary Figure S18). When gene conversion rates are constrained (standard deviation of 1x10^-8^) then mutation counts usually increase over time in line with mutation accumulation, unless the starting rate is high (4x10^-2^) in which case the mutation count is consistently low. When gene conversion can more freely evolve (standard deviation of 1x10^-7^) then both total mutation and homozygote counts match the same values, reflecting similar convergence of mean fitness.

**Figure 9:**
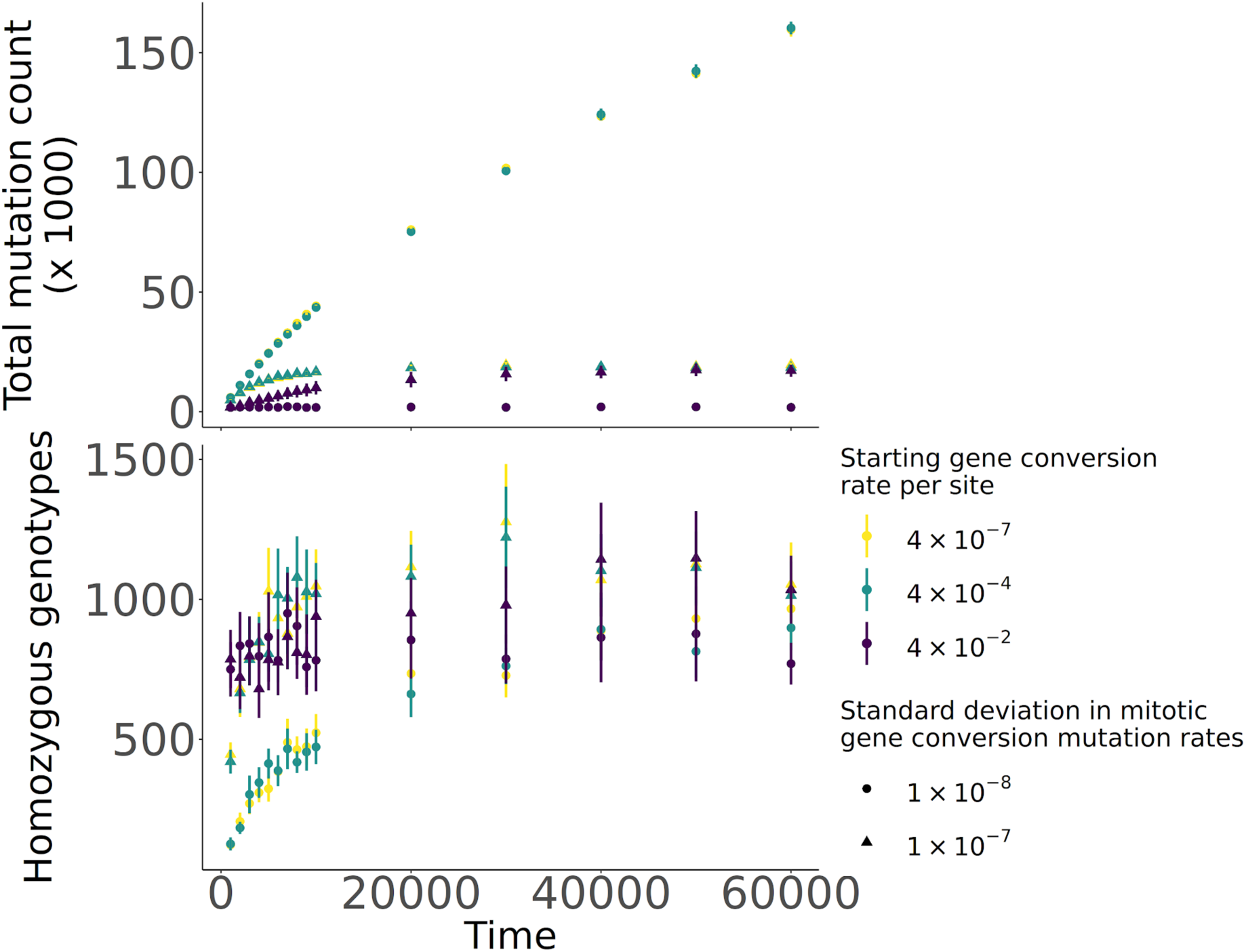
Deleterious mutation counts under complete asexuality when gene conversion is allowed to evolve. As a function of time, these plots show (top) the total mutation count present in the sample of 50 individuals, and (bottom) the number of homozygous genotypes present in the sample. Error bars represent 95% confidence intervals; if they are not shown then they lie within a point. See Supplementary Figure S18 for the relative homozygosity. **Alt text:** Graphs showing total mutation count and number of homozygote genotypes when mitotic gene conversion can evolve. Different colours indicate the starting rate, different shapes represent the extent to which gene conversion can evolve.

Overall, these results show that when gene conversion rates can evolve, they do not necessarily do so to ensure the highest fitness is attained. Evolving gene conversion is mainly beneficial when the initial rates are low, and it is more able to evolve larger values. However, if initial gene conversion rates are high, then freely-evolving rates are detrimental and lead to lower fitness than if it were fixed.

## Discussion

### Summary of results

In this study, we theoretically explored to what extent mitotic gene conversion in facultative sexual species purge deleterious mutations. We additionally compared this process to self-fertilisation, as the latter is known to purge deleterious mutations so we can compare how effective the two are. Using a single-locus model (Figure 1), we first demonstrated that mitotic gene conversion has the greatest effect in reducing mutation frequency when asexuality is common, but also when mutations are recessive. We also show that, while the process leads to elevated homozygosity (as measured by the inbreeding coefficient *F,* Figure 2), it will not be substantially elevated and match the effects of selfing unless populations are fully asexual and the gene conversion rate is high. We subsequently used multi-locus simulations to explore the effect of mitotic gene conversion acting on a genome-wide level. In the absence of gene conversion (Figures 3, 4), highly-asexual species accumulate many mutations as heterozygotes, leading to a drastic drop in fitness. When gene conversion is present in fully asexual populations (Figures 5, 6), then we do see purging: once the gene conversion rate exceeds a critical value (generally a mean rate of 4x10^-4^ per site) the total mutation count drops and homozygosity increases. These levels of gene conversion also dramatically increase the fitness of asexual populations (Figure 5, Supplementary Figure S14). Despite these benefits of gene conversion, the total mutation count is lowest under high levels of self-fertilisation (Figure 7).

Overall, both the single-locus and multi-locus results indicate that while high rates of gene conversion can purge deleterious mutations under asexual reproduction, these benefits do not always counter the costs of inefficient selection under clonality; self-fertilisation can act a stronger force at purging deleterious mutations. This conclusion is reinforced when running simulations where we let gene conversion rates evolve (Figures 8, 9). In all simulations mitotic gene conversion rates evolve to high values (between 1x10^-4^ and 1x10^-1^) but the outcomes on mean fitness differ depending on the starting rate and the ability to which it can evolve. When rates are initially low then it only reaches high-enough values to increase mean fitness if it can freely evolve (i.e., when the standard deviation in mutated values equals 1x10^-7^). However, such lability in gene conversion rates is detrimental if they are already high. We suspect the reason why high gene conversion rates are not maintained is that, once it has arisen to purge a majority of deleterious mutations, then there is not a sustained evolutionary pressure to maintain high values. A similar argument has been made for how sexual reproduction can be difficult to maintain; once sex has created favourable genetic combinations, then there can be evolutionary pressure to reduce it as otherwise it will break them up (Otto and Michalakis 1998; Otto 2009).

### Comparing mutation accumulation in self-fertilising and asexual populations

One notable feature of the asexual-selfing comparison is the threshold behaviour of mutation counts. In the absence of gene conversion, neutral genetic diversity in asexual populations is similar to outcrossers unless the rate of sex is of the order 1/*N* (Bengtsson 2003; Ceplitis 2003; Hartfield et al. 2016; Hartfield 2021; Pieszko et al. 2025); background selection also incurs noticeable reductions in diversity in species with infrequent sex (Agrawal and Hartfield 2016). We see similar results regarding the mutation count in Figure 4; it stays fairly stable under facultative sex unless populations become highly asexual. Conversely, under high selfing we generally see a drop in the mutation count and heightened homozygosity, in line with mutation purging. With very high levels of selfing, we see an elevation in both the homozygous mutation count and in the total mutation number. These increases likely arise due to higher mutation numbers caused by genetic drift, as *N_e_* can be greatly reduced in selfing populations due to background selection (Kamran-Disfani and Agrawal 2014; Roze 2016). Note that this behaviour can change with the fitness model assumptions; under a model where *s* and *h* are drawn from distributions, we instead see a large reduction in the mutation count under high selfing, in line with an outcome where genetic variation is purged (Charlesworth and Charlesworth 1995; Lande and Porcher 2015; Abu Awad and Roze 2018).

### Effects of mitotic gene conversion, and comparison with empirical estimates

Throughout this study, we compared purging via mitotic gene conversion in asexuals to that via self-fertilisation. We showed that despite the benefits of mitotic gene conversion, lower mutation counts were observed under high levels of self-fertilisation (see, e.g., Figure 7). These results indicate that self-fertilisation can be a more efficient means of purging deleterious mutations, likely because it affects the whole genome and homozygous regions can be formed more rapidly. In contrast, mitotic gene conversion as employed here only affects a fraction of the genome every generation, so purging acts in a piecemeal manner. That said, sufficiently high mitotic gene conversion could restore fitness to some degree through deleterious mutation purging. This comparison seems to hold over different mutation models, selfing and gene conversion rates; one caveat is that there could be a specific, untested combination of parameters that might lead to different results.

Our simulation model shows that a mean mitotic gene conversion rate above 4x10^-4^ per site per generation can strongly purge deleterious mutations, while a rate of 4x10^-5^ can prevent their accumulation. This is in line with previous theoretical work showing that, once gene conversion becomes sufficiently high, then it can substantially reduce genetic variation in asexual populations (Hartfield et al. 2016; Hartfield 2021). Although empirical estimates are sparse, they are usually much lower than this value, ranging between 10^-7^ and 10^-5^ depending on the study organism (Mandegar and Otto 2007; Xu et al. 2011; Flot et al. 2013; Sharp and Agrawal 2016). Potentially higher rates were observed in rice; Jia et al. (2021) inferred a mitotic recombination rate of 10^-6^ per marker with mean tract lengths ranging between 30-50kb per gene conversion event; assuming one event per mitosis, this leads to a per-locus, per-mitosis rate on the order of 10^-2^. Hence, it seems that while some empirical estimates are sufficiently high to cause extensive deleterious mutation purging, on the whole mitotic gene conversion is usually too weak a force to lead to extensive purging in nature. However, the estimates of Jia et al. (2021) consider all mitotic recombination events, including crossover and non-crossover events. Our simulation procedure only considers non-crossover events, which converts more localised genomic regions. Mitotic crossover events, in contrast, can convert much larger fractions of the genome (Jia et al. 2021). Even though the net conversion rate per site seems the most important parameter, longer conversion regions could have a stronger effect through affecting multiple sites at once, in a similar manner to self-fertilisation. We also only have estimates from a handful of species, so our empirical understanding of mitotic gene conversion is far from complete (Jinks-Robertson and Petes 2021). In addition, there has been some debate as to whether meiotic gene conversion is biased to counter deleterious mutation load, although evidence for this mechanism is mixed (Liu et al. 2018). Additional empirical estimates, along with further information on whether mitotic conversion is biased, will shed light on whether sufficiently frequent conversion rates exist that can purge deleterious mutations.

Other forms of asexual reproduction that create extensive homozygous regions, including automixis (Mirzaghaderi and Hörandl 2016; Engelstädter 2017) and endomitosis (i.e., spontaneous chromosome doubling; Lamb and Willey 1987), could also affect enough of the genome to cause large levels of mutation purging. Future research can determine whether these different factors have a strong effect on asexual fitness.

### Caveats and perspective for future work

Another factor not considered in our study is the effect of small and/or changing population sizes. In small populations, homozygotes can form more frequently due to genetic drift (Malécot 1948), which can also lead to the purging of deleterious mutations. However, this process is only effective at purging highly recessive mutations (Glémin 2003). Gene conversion and drift also interact to affect the evolution of sex (Roze and Michod 2010). Given the large diploid population size of 5,000 used in simulations, then drift should have a minimal influence on purging in the multi-locus simulations, especially since the prevalence of deleterious mutations did not strongly change under different population sizes (Supplementary Figures S8, S9). However, the population size could affect the baseline level of deleterious mutations in the absence of gene conversion. It will be a rewarding avenue of future research to determine how drift would affect these processes. Similarly, we also did not consider demographic feedbacks that are induced during purging that cause a reduction in population size (Abu Awad and Billiard 2017). The presence of these feedbacks could increase the extinction probability of populations during gene-conversion induced purging. Another avenue for further investigation is the role of adaptive evolution and/or epistasis on the relative advantage of different reproduction strategies, as they could impact the relative fitness of heterozygotes and homozygotes and hence the outcomes of the reproductive modes studied here (Kondrashov and Kondrashov 2001; Abu Awad and Roze 2020).

The model we used to simulate evolving mitotic gene conversion rates approximated it as being a complex trait, where the change in an individual’s rate is drawn from a normal distribution. Hence, there is plenty of scope for exploring this process more deeply in future work. For example, one can further investigate if results differ depending on whether a single modifier gene or a polygenic basis determines mitotic gene conversion rates. In addition, given these rates do not seem to evolve to high values, it will be of interest to investigate how mitotic gene conversion evolution, selection, and genetic drift interact to determine how likely it evolves to remove deleterious mutations.

We only considered allelic gene conversion in this model, given the focus on previous empirical studies proposing that allelic conversion could prevent mutation loads in asexual species. Non-allelic conversion, which happens along a haplotype rather than between haplotypes, can also arise, especially between genes of the same gene family. Previous theoretical papers have also shown that it can reduce mutation loads under specific conditions (Ohta 1989; Hurst and Smith 1998) and prevent Y-chromosome degeneration (Marais et al. 2010). There is empirical evidence that non-allelic conversion affects human histone evolution (Galtier 2003), but it is unknown to what extent it acts in facultatively sexual species. Hence, determining the prevalence of non-allelic conversion in these species would be welcome, along with theoretical explorations of its potential in removing deleterious variants.

We assume that populations are diploid to more cleanly compare the effect of mitotic gene conversion between sexual and asexual populations. However, polyploidy is pervasive in asexual organisms (Neiman et al. 2014). We have little biological knowledge of how mitotic gene conversion acts in polyploids, and future modelling work can investigate whether mutation purging would still be possible. We argue that the same general arguments presented here will apply to polyploid asexuals, as long as pairing and gene exchange is permissible between homologous chromosomes.

We also assumed that the mutation and gene conversion rates were the same irrespective of how species reproduced. Specifically, we focussed on mitotic gene conversion as the mitotic pathway is common to both sexually- and asexually-reproducing species. However, given that mitosis and meiosis are distinct cellular processes (Jinks-Robertson and Petes 2021), it could be the case that gene conversion and mutation rates differ between them. For example, some evidence exists that mitotic mutation rates are lower than meiotic mutation rates, but can still make a significant contribution to the total number of inherited mutations (Magni and Von Borstel 1962; Schoen and Schultz 2019; Villalba De La Peña et al. 2023). We argue that in cases where both mitotic and meiotic events affect genetic diversity in offspring (e.g., somatic mutations in plants; Schoen and Schultz 2019), then it is the totality of these rates that determine whether gene conversion can be selected to counter mutation accumulation.

An interesting case where gene conversion (either allelic or non-allelic) may be important under clonal reproduction is tumour evolution. Here, cells reproduce asexually by mitosis and can potentially accumulate deleterious mutations due to a lack of recombination (Ní Leathlobhair and Lenski 2022). Hence, the gene conversion mechanisms investigated here may prevent transmissible cancers from experiencing mutation meltdown (Ní Leathlobhair and Lenski 2022), and also create homozygote loss-of-function tumour-suppressing genes that enable carcinogenesis (Takahashi and Innan 2022).

### Multi-Locus Simulation Methods

#### Simulation Outline

Simulations were written using SLiM v4.0.1 (Haller and Messer 2023), where we used one of two types of simulation framework. We first used the Wright-Fisher framework (hereafter ‘WF model’) when exploring population fitness and mutation counts for general levels of uniparental reproduction, in the absence of mitotic gene conversion. In these simulations, the population size is fixed and the probability of choosing parents are proportional to their fitness. We next ran simulations using the non-Wright-Fisher framework (hereafter ‘non-WF model’) when investigating the effects of mitotic gene conversion in fully asexual populations. By default in non-WF models, the population size can change depending on which parents reproduce or not. However, as explained below, we modify the non-WF behaviour in SLiM so that it acts like a comparable WF model.

Simulated populations consist of 5,000 diploid individuals, each with a genome size of *L* = 25 *Mb* that approximates the length of *Arabidopsis thaliana* chromosome 1 (The Arabidopsis Genome Initiative 2000). Deleterious mutations can arise at all nucleotides, and individuals do not carry mutations at the start of the simulations. In all simulations, we let populations evolve for 60,000 generations, which is sufficient for the population variance to reach a steady-state (Supplementary Figures S19 - S21). After this time, 50 individuals are sampled and their genotypes were saved in a VCF file format for downstream analysis. The mutation rate was set to 4x10^-9^ per basepair, which is on the same order as that inferred from *Arabidopsis thaliana* (Ossowski et al. 2010). Per-base-position recombination rate was set to 4x10^-8^, reflecting estimates obtained from crosses of *Arabidopsis thaliana* (Salome et al. 2012).

Mutations were deleterious and mutation stacking (a SLiM-specific technical feature where multiple mutations can occupy the same site within an individual) is allowed by default. Two types of fitness models were investigated. They differ in whether selection and dominance coefficients were constant or drawn from a distribution (Table 1). In the first type, fixed values of deleterious mutations and dominance coefficients were chosen. In the second type, the selection and dominance coefficients for each mutation were drawn from a distribution based on Spigler et al. (2017) (see Table 1). Selection coefficient is modelled by a gamma distribution with shape parameter *α* and mean *k̄*. The dominance coefficient of a mutation with selection coefficient *s* is then drawn from a uniform distribution bounded by 0 and *e^-ks^*, where *k̄* is a constant (see Table 1 for exact values). Free parameters in the model, *k*, *s̄*, and *α*, were defined so *h* matches that used in main simulations (0.2) when *s̄* = 0.01 using a shape parameter *α* = 0.02 that reflects values inferred by (Chen et al. 2017); note that this parameterisation produces ℎ; ≈ 0.35. This model reflects empirical observations that strongly deleterious mutations tend to be more recessive (Simmons and Crow 1977; Agrawal and Whitlock 2011).

**Table 1.** Summary of model parameters and values used for simulations in the main text. Note that in the Supplementary Material, we show results with different population sizes (*N* equalling 1,000 and 10,000) but with the mutation rate and selection coefficients scaled so *Nμ, Ns* are the same as for *N* = 5,000 simulations.

| Parameter | Values for simulations with f. e.* |  |
| --- | --- | --- |
|  | constant | non-constant |
| Genome size | Diploid, L = 25 Mbp |  |
| Per-base mutation rate $\mu$ | $4 \times 10^{-9}$ | |
| Per-position recombination rate $r$ | $4 \times 10^{-8}$ | |
| Population size $N$ | 5 000 | |
| Sample size | 50 |  |
| Selection | $s = 0.01$ | $s \sim \text{Gamma}(\bar{s}, \alpha)$<br>mean $\bar{s} = 0.01$<br>shape $\alpha = 0.2$ |
| Dominance | $h = 0.2$ | $h \sim U(0, e^{-ks})$<br>$k = 91.63$ |
| Mean gene conversion tract length | $\lambda = 4\,000$ bp | |
| Standard Deviation in gene conversion rate (when it evolves) | $1 \times 10^{-8}; 1 \times 10^{-7}$ | |

\* f. e. - fitness effects; selection and dominance coefficients

### Implementing mitotic gene conversion in asexual reproduction

By default, recombination in SLiM (including gene conversion) only acts exclusively when meiosis is involved. Therefore, the reproduction cycle was manually modified through a reproduction callback (i.e., a SLiM function for governing reproduction), so that mitotic gene conversion in asexuals can be simulated; this can only be done in the non-WF model. For simplicity, the model applies only to a completely asexual population. The development of the code was facilitated by a post on the SLiM discussion board (Dussert and Haller 2019) and was extended based on the gene conversion model under SLiM Wright-Fisher simulations (Haller and Messer 2016).

When an offspring is generated, then the number of gene conversion events that acts within it is drawn from a Poisson distribution with a mean *L*γ where γ denotes the rate of gene conversion initiating at a site. For each gene conversion event, the central position is first drawn uniformly through the chromosome. Then, the tract is extended to the left and right independently by drawing each length from a geometric distribution with parameter *p:*

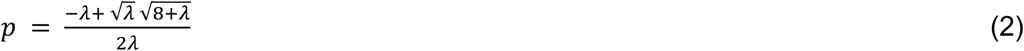

where *λ* is the total mean tract length we want to use in simulations. This parameterization is used to account for the fact that gene conversion tracts with length zero are discarded. This redefined value of *p* means that the mean tract length is λ after accounting for discarding these tracts (see Supplementary *Mathematica* notebook for further information). Gene conversion events that overlap with the chromosome ends are truncated so they stop at the chromosome edges.

After the gene conversion start- and end-points are drawn, they are checked and potentially modified as follows: tracts of length 0 are removed; gene conversion tracts may have the same start or end positions, in which case, the start (or end) of one of them is shifted to the right (left), so that gene conversion tracts overlap; or two gene conversion tracts overlap, in which case two interior gene conversion positions are removed so that the resulting gene conversion tract is a union of the two smaller ones. Since multiple gene conversion tracts can overlap, this refinement process is repeated until all start and end positions are unique and alternate after sorting.

Gene conversion can occur on either of the homologous chromosomes with equal probability and results in the selected part of one chromosome being copied onto the same position at the other chromosome. Note that for each offspring-production event, all gene conversion tracts are in the same direction, i.e., material is transferred solely from one strand to another during each reproduction event. Since the ‘initiation’ strand is randomly chosen for each reproduction event, then this effect should not lead to a major bias in results. The mean length of a gene conversion event is set to 4,000 base pairs; this value accounts for the fact that empirical studies of mitotic gene conversion tends to find that gene conversion tracts are bimodally distributed, with a combination of short (on the order of 1kb) and long tracts (on the order of 10kb) (Yim et al. 2014; Chumki et al. 2016; Jia et al. 2021). Using a mean length of 4,000bp will thus enable a wide distribution of tract lengths to be drawn that cover both these possibilities. We report simulation results in terms of the ‘mean gene conversion rate per site’, which is equal to the initiation rate times the mean tract length. This value is approximately equivalent to the gene conversion parameter *γ* in the single-locus model (Frisse et al. 2001).

To enable fair comparison between simulations, further changes to the non-WF model were made so that it behaves as a standard Wright-Fisher model with a fixed population size over time (which is explained further in the SLiM Manual; Haller and Messer 2016). Specifically, a survival callback (i.e., a SLiM function for regulating individual mortality) limits the age of an individual to a single generation, and a reproduction callback samples a constant number of parents with probability weighted by their fitness. By comparing measurements of population fitness (Supplementary Figure S22) and mutation count (Supplementary Figure S23) between the two types of simulations, we see that this new routine yields similar behaviour under low gene conversion rates to the original WF model with complete asexuality. 100 replicate simulations were run if *s*, *h* were fixed; 20 replicate simulations were run for WF simulations (i.e., with partial selfing and cloning) where *s*, *h* are drawn from distributions; and 10 replicate simulations were run for non-WF simulations (i.e., with asexuality and gene conversion) where *s*, *h* are drawn from distributions. Fewer simulations replicates were used in the latter two cases due to their longer runtime.

### Mutational Load Measurement

Although we can measure the genetic load exactly from our simulations, in practice they are hard to measure from empirical data, as they usually require assumptions of the underlying parameters (e.g., selection and dominance coefficients) that have considerable uncertainty around them (Henn et al. 2016; Robinson et al. 2023; Kyriazis and Lohmueller 2024). Instead, empirical studies often use proxy measurements for load, specifically the total number of deleterious mutations, and those present as homozygotes (Marsden et al. 2015; Henn et al. 2016; Robinson et al. 2016; Laenen et al. 2018; Grossen et al. 2020; Hoffman et al. 2024). These measurements are simple ways to gauge deleterious mutation prevalence, and their approximate contributions to fitness reductions in a way that can be easily measured from genome data and compared across species with different reproductive modes in nature. For example, Laenen et al. (2018) used these measurements in an analysis of *Arabis alpina* populations across Europe, to compare deleterious mutation prevalence in populations with different degrees of inbreeding via self-fertilisation. Hence, we present deleterious mutation prevalence from simulations in the same way here, to better enable empirical comparisons with these theoretical results and to understand the relative contributions of heterozygotes and homozygotes to deleterious mutation prevalence.

From the sampled populations, we calculated (i) the total number of mutations present in the population, and (ii) the number of homozygous mutant genotypes. These measurements are proportional to the population’s genetic load, assuming a fixed selection coefficient, and the realised load for recessive mutations with a dominance coefficient of zero (Bertorelle et al. 2022). We also calculated the relative homozygosity for each simulation, defined as 2*(number of homozygous mutations)/(total number of mutations), which can be interpreted as the proportion of deleterious mutations that are in homozygous state, across all sites and individuals. Simulation outputs were analysed using R 4.2.1 (R Core Team 2024).

### Implementing Evolving Gene Conversion

We also ran simulations where the gene conversion rate was a property of individuals and could evolve in the population. These simulations were only run for cases where *s*, *h* were fixed. Gene conversion was set as a ‘tag’ variable, which is a SLiM construct for assigning values to each individual. The initial gene conversion rate was set to one of three values, to test if the starting rate had an effect on its ability to evolve: 4x10^-7^, 4x10^-4^, or 4x10^-2^ (after accounting for the mean tract length of 4,000 base pairs). Every generation, the gene conversion rate of an offspring was shifted relative to the parent by adding a value drawn from a normal distribution with mean zero, and a standard deviation of 1x10^-8^ or 1x10^-7^ (so the added value could also be negative, to reduce the gene conversion rate). If the resulting rate was negative, it was then set to zero. Simulations were left to evolve for 60,000 generations as before. Mutation counts were sampled every 1,000 generations for the first 10,000 generations, then every 10,000 generations thereafter. Denser mutation sampling was used at the start of simulations so we could better track how mutation accumulation was affected by evolving gene conversion rates. We tested this model by running it without gene conversion evolution, and obtained the same mean fitness and mutation count values as original simulations (Supplementary Figure S24). 20 replicate simulations were run in each case.

## Supporting information

Supplementary Mathematica File

## Data availability statement

Simulation code and Conda environments with all used packages, together with all scripts have been archived on GitHub (https://github.com/dominik-kopcak/SelfClonGC).

## Author contributions

MH designed the project with input from DK. DK wrote and ran the simulations, and analysed the resulting outputs with feedback from MH. MH developed and analysed the single-locus model. MH and DK wrote and edited the manuscript.

## Acknowledgements

We would like to thank Anders Poulsen Charmouh for advising on how to implement the non-constant *s*, *h* model, and members of the Hartfield lab and anonymous reviewers for providing feedback on the work.

## Supplementary Text

### Appendix 1: Deriving single-locus results for the frequency of deleterious mutations and inbreeding coefficients

In the single-locus model, we assume a lifecycle consisting of selection, reproduction (either self-fertilisation or facultative sex), mutation, then mitotic gene conversion. We derive recursions for the frequency of the three genotypes (*aa*, *Aa*, and *AA* for *a* the wild-type mutations and *A* the derived, deleterious mutation), convert them to allele frequencies and inbreeding coefficients, then solve for steady-state values. We let *p_g_* denote the frequency of genotype *g* (e.g., *p_AA_* is the frequency of genotype *AA*).

#### Selection

Let fitness for the three genotypes (*aa*, *Aa*, and *AA*), which we denote *W_g_*, be 1, 1–*hs* and 1–*s* respectively. The mean fitness equals W_m_ = *p_aa_* + *p_Aa_* (1–*hs*) + *p_AA_* (1–*s*). The change in frequency of genotype *g* over a single generation of selection is:

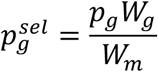

#### Reproduction

We can partition reproduction into three outcomes; via outcrossing, via selfing, or via clonal reproduction. For outcrossing, we assume allele frequencies segregate independently. Hence, for example, the frequency of ancestral homozygotes after outcrossing

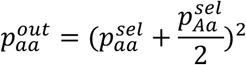

with a similar calculation for derived homozygotes *AA*. The frequency of heterozygotes after outcrossing is:

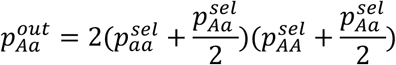

Under selfing, homozygotes will only produce homozygote offspring, while heterozygotes will produce heterozygotes with probability ½ and either of the homozygote with frequency ¼ each. This yields the recursion equation for the change in homozygotes following selfing (with the example given below for *aa* homozygotes):

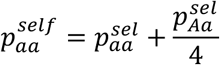

The change in heterozygote frequency is:

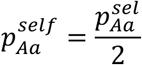

Under clonal reproduction, then genotype frequencies are unchanged between parent and offspring.

If self-fertilisation occurs with frequency *S*, then we can calculate the genotype frequencies after reproduction by weighing the contribution from outcrossing and selfing accordingly:

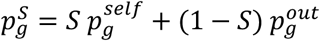

Similarly, we can calculate the change in genotype frequencies following facultative sex, given a frequency of sex *σ* and clonal reproduction 1– *σ*:

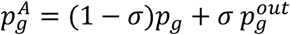

In subsequent sections we use 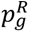 to denote the allele frequency following any kind of reproduction.

#### Mutation

We assume unidirectional mutation, i.e. allele *a* changes to *A* with probability *μ*. Hence, *aa* mutates to *Aa* with probability 2 *μ* (1– *μ*), *aa* mutates to *AA* with probability *μ*^2^, and *Aa* mutates to *AA* with probability *μ*. This process yields the following recursions for 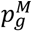, the change in the frequency of genotype *g* after mutation:

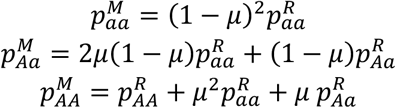

#### Mitotic gene conversion

Mitotic gene conversion acts with probability *γ*. It only affects heterozygotes, and turns them into either homozygotes with probability *γ*/2. This process yields the following recursions, which we denote 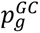:

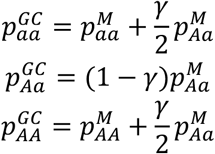

#### Solving for steady-states

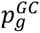 denotes the final change in genotype frequency following one generation. We can use the change in genotype frequencies to derive the change in frequency for allele *A*, which equals 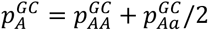. We can also derive the change in inbreeding coefficient:

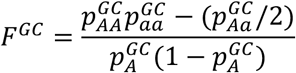

To then convert genotypes into allele frequencies, we can write each genotype in terms of allele frequencies, as described in the main text. For example, the frequency of *AA* can be written as *p_A_ + p_A_(1– p_A_)F*. These equations can be complicated, so we simplify them by taking a Taylor series assuming that the mutation rate *μ* and *p_A_* are both small (details are given in the Supplementary *Mathematica* file). From these simplified equations, we jointly solve for 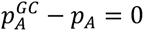 and *F^GC^* − *F* = 0, yielding Equations 1–3 in the main text.

### Comparison with the model used by Flot et al. (2013)

There are some differences between the model used here and that employed by Flot *et al*. (2013). First, when modelling mutation we initially model the exact process, but subsequently linearise equations to remove terms of order *μ*^2^ and above. This linearisation is also employed by Flot *et al*. (2013). Second, we use gamma for the overall gene conversion rate, with a rate 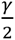 that a heterozygote is converted into each homozygote. Conversely, Flot *et al*. (2013) use *γ* to denote the conversion rate from heterozygote to each homozygote, so is effectively twice as large as what is used here. Finally, Flot *et al*. (2013) considered a stochastic model of mutation accumulation, while we use a deterministic model that does not take population-size effects into account.

**Figure S1:**
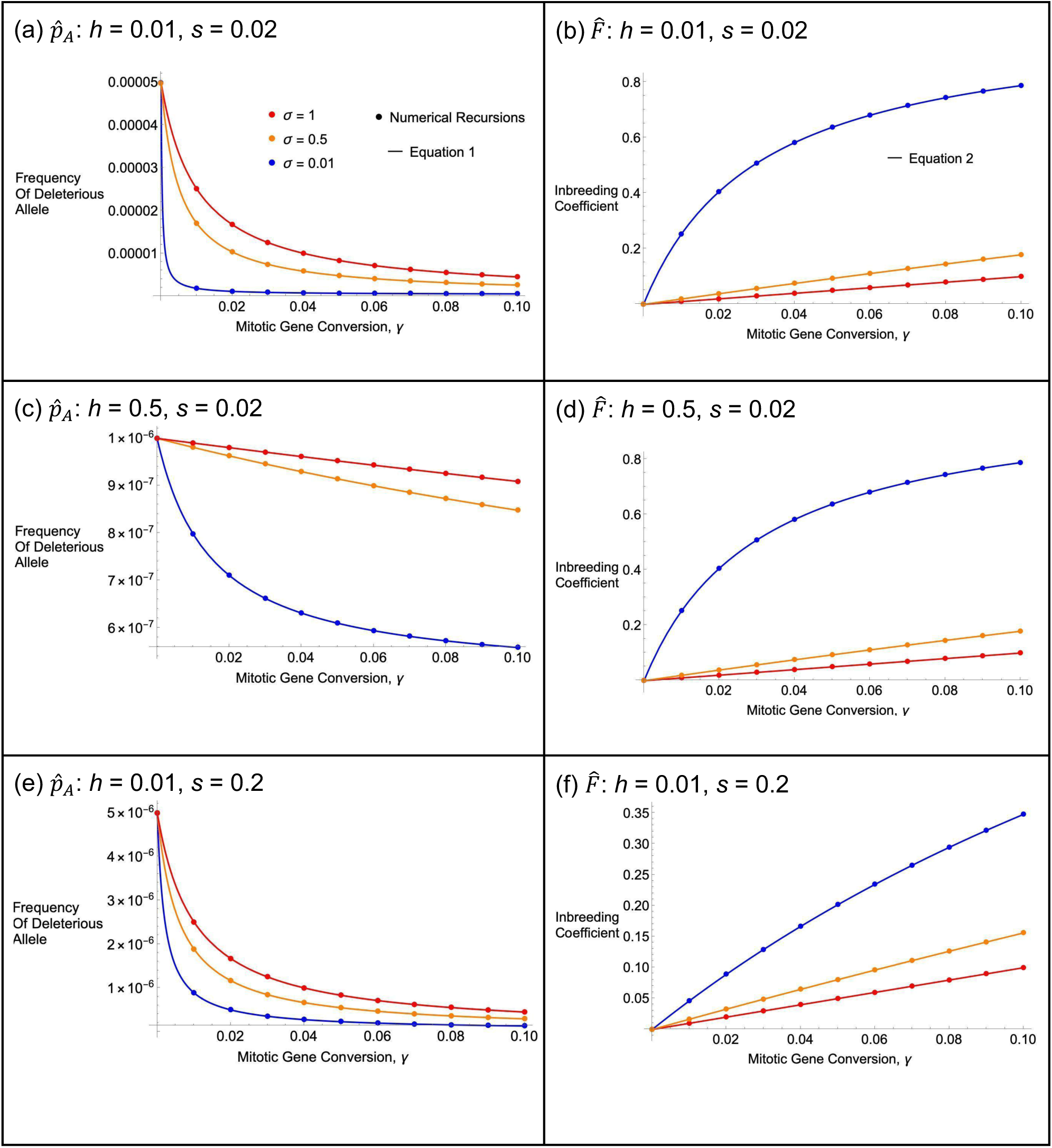
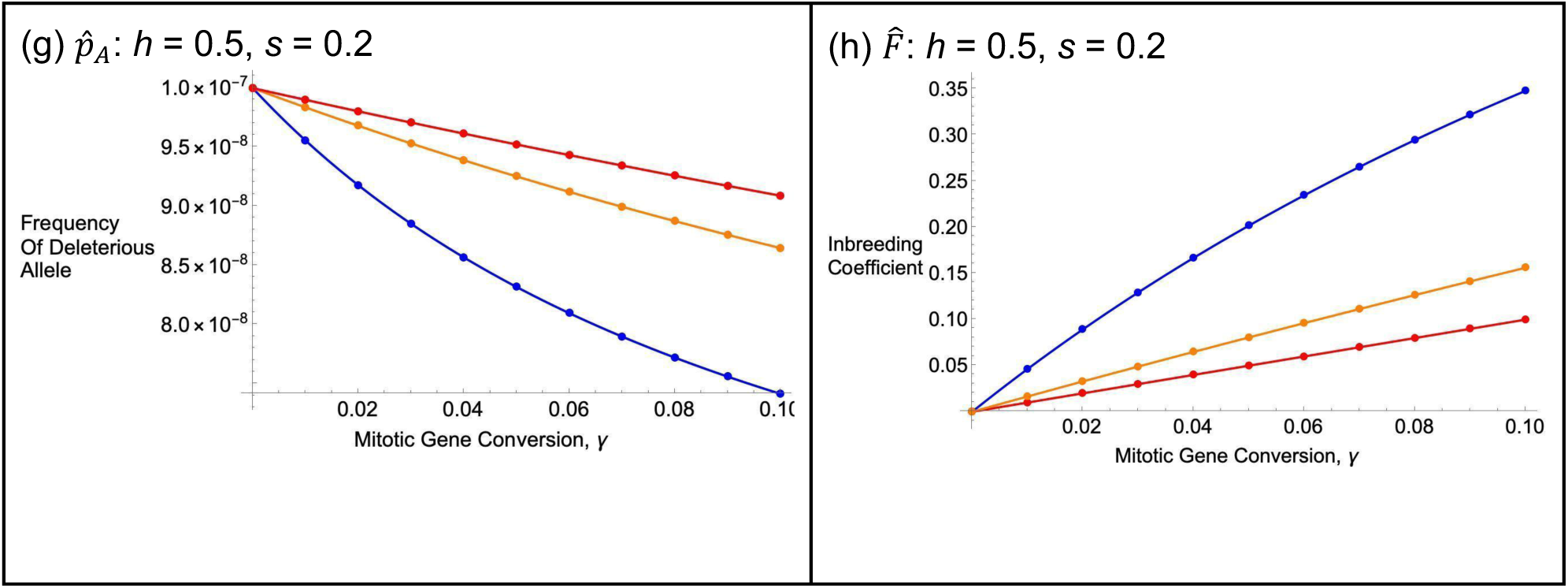
Comparing analytical equations against exact numerical recursions, for varying rates of sex. As a function of the rate of gene conversion, (a), (c), (e), and (g) show the frequency of the deleterious mutation (Equation 1 in the main text), and (b), (d), (f) and (h) show the corresponding inbreeding coefficient F (Equation 2 in the main text). Closed circles gave values for the numerical recursions. Results are shown for different frequencies of asexual reproduction, as shown in the legend for panel (a). Other parameters are μ = 10^-8^, and in the recursions the initial frequency of genotype Aa is 0.001 and AA is zero, and recursions were run for 50,000 generations.

**Figure S2:**
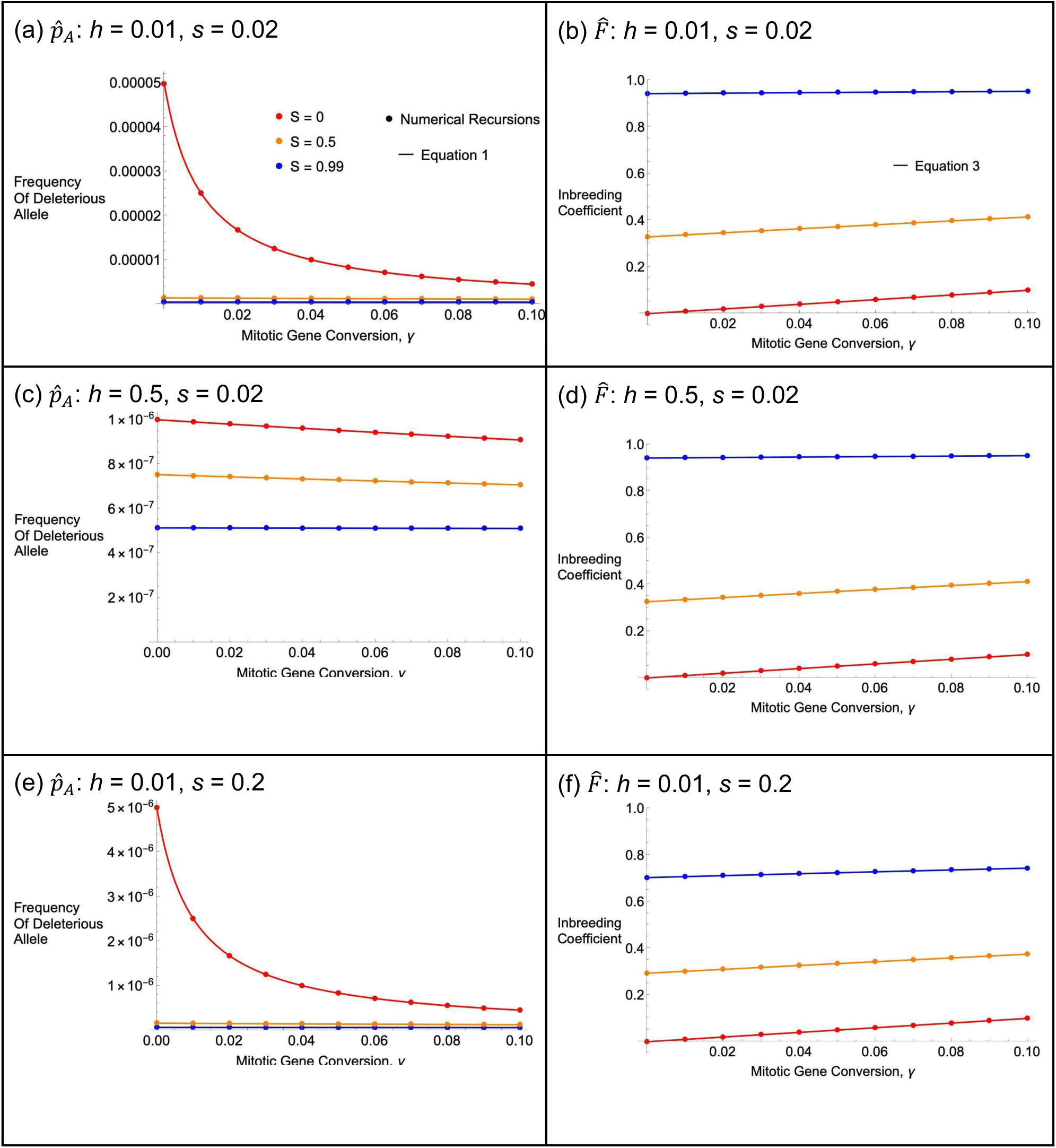
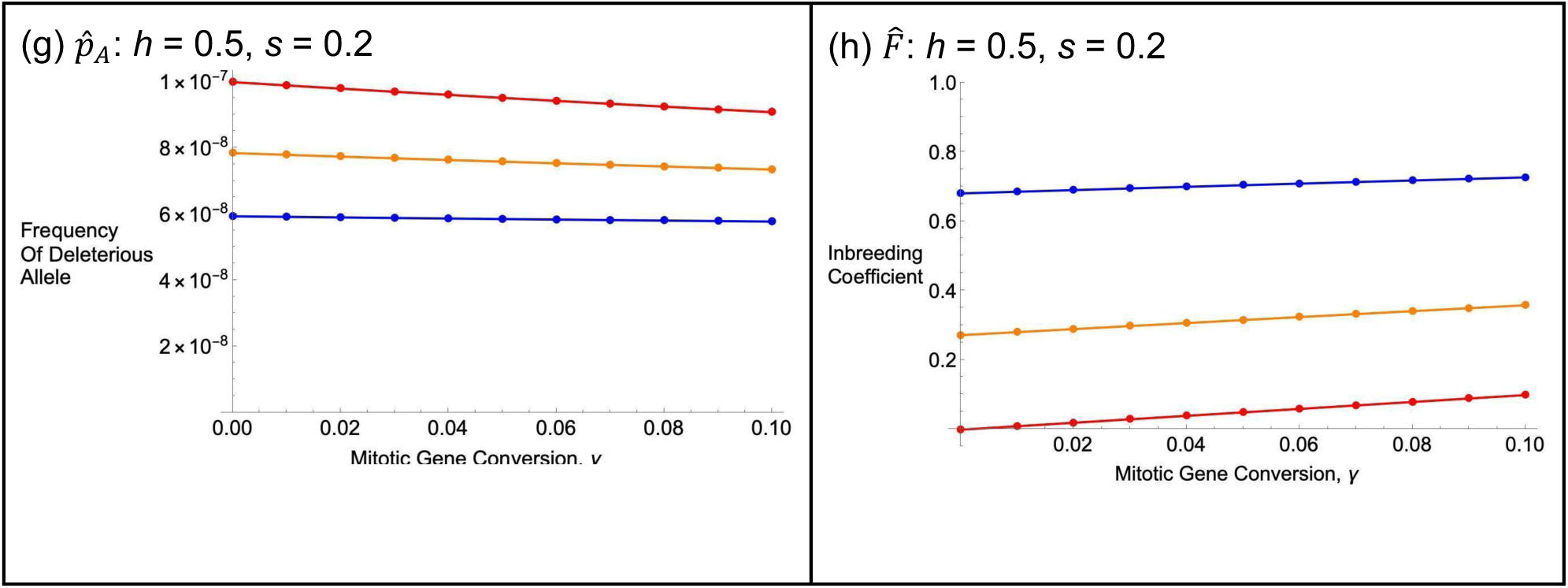
As Figure S1 but computed over different rates of self-fertilisation. The plotted inbreeding coefficient is Equation 3 in the main text; closed circles give results from numerical recursions.

**Figure S3:**
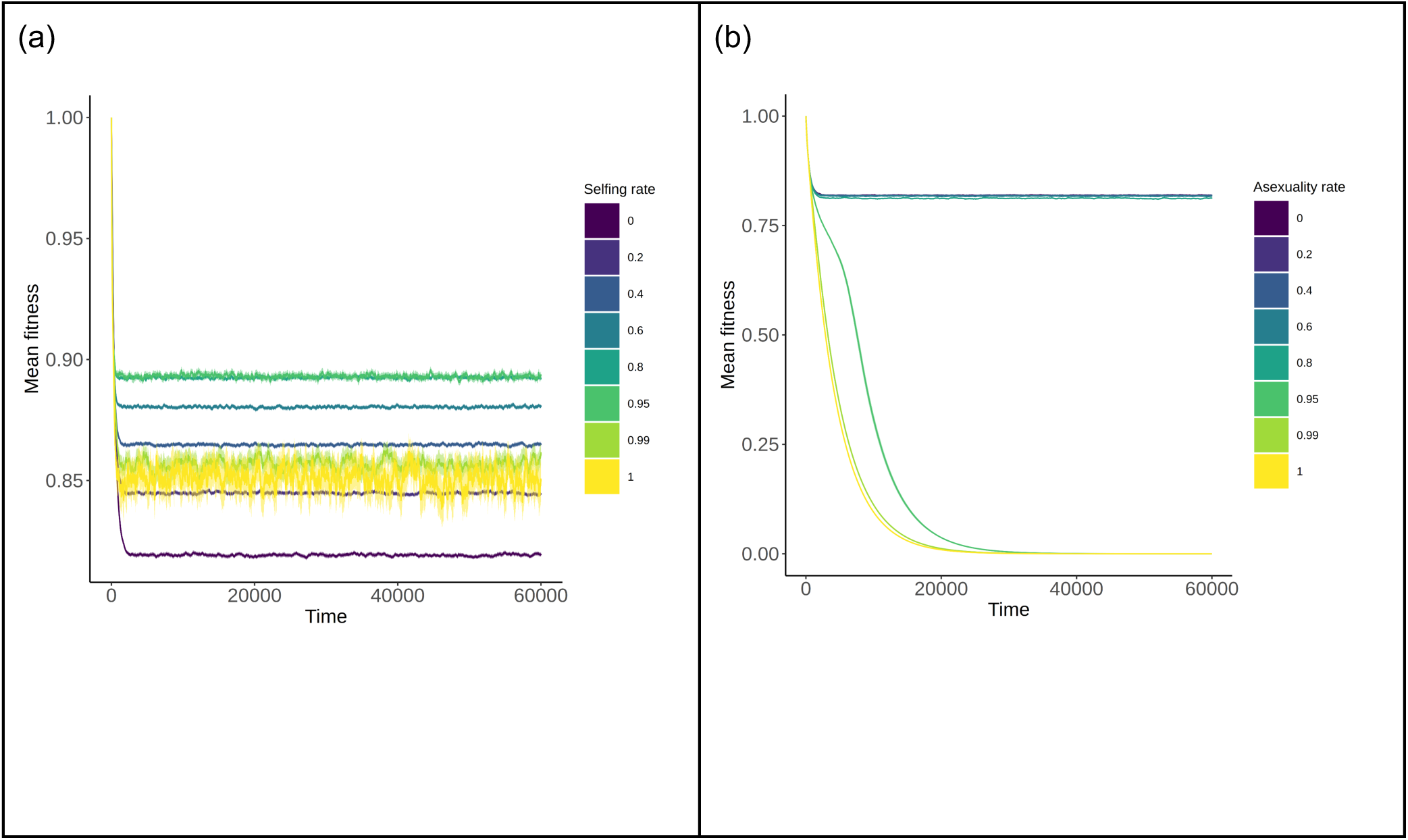
Mean fitness for different modes of uniparental reproduction. Results are shown as a function of time, for (a) different rates of self-fertilisation or (b) asexual reproduction. Thick lines are mean values, while thinner shading around lines show 95% confidence intervals. Notice the change of scale on the y-axis between (a) and (b).

**Figure S4:**
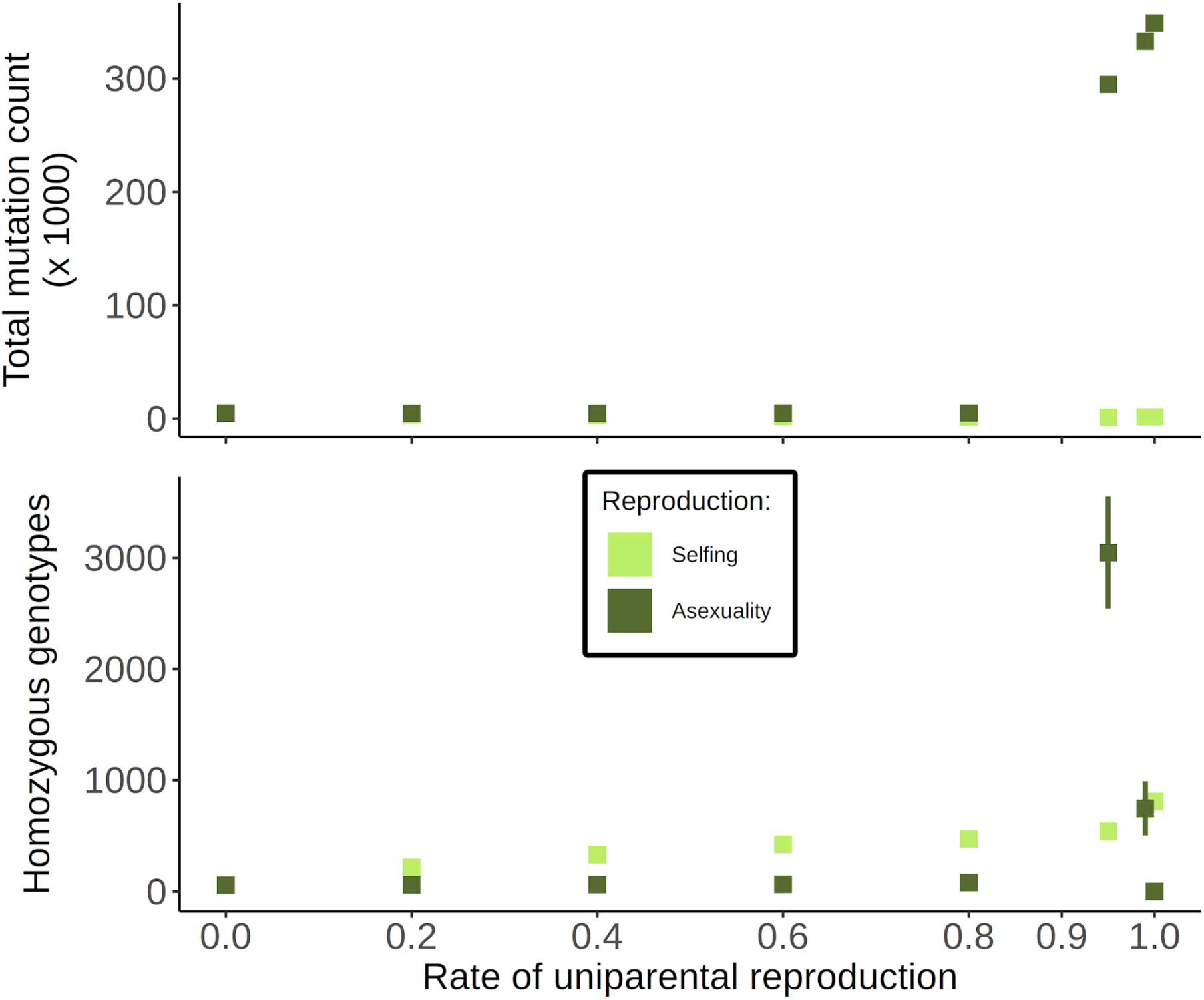
As Figure 4 in the main text, but with total count on a linear scale.

**Figure S5:**
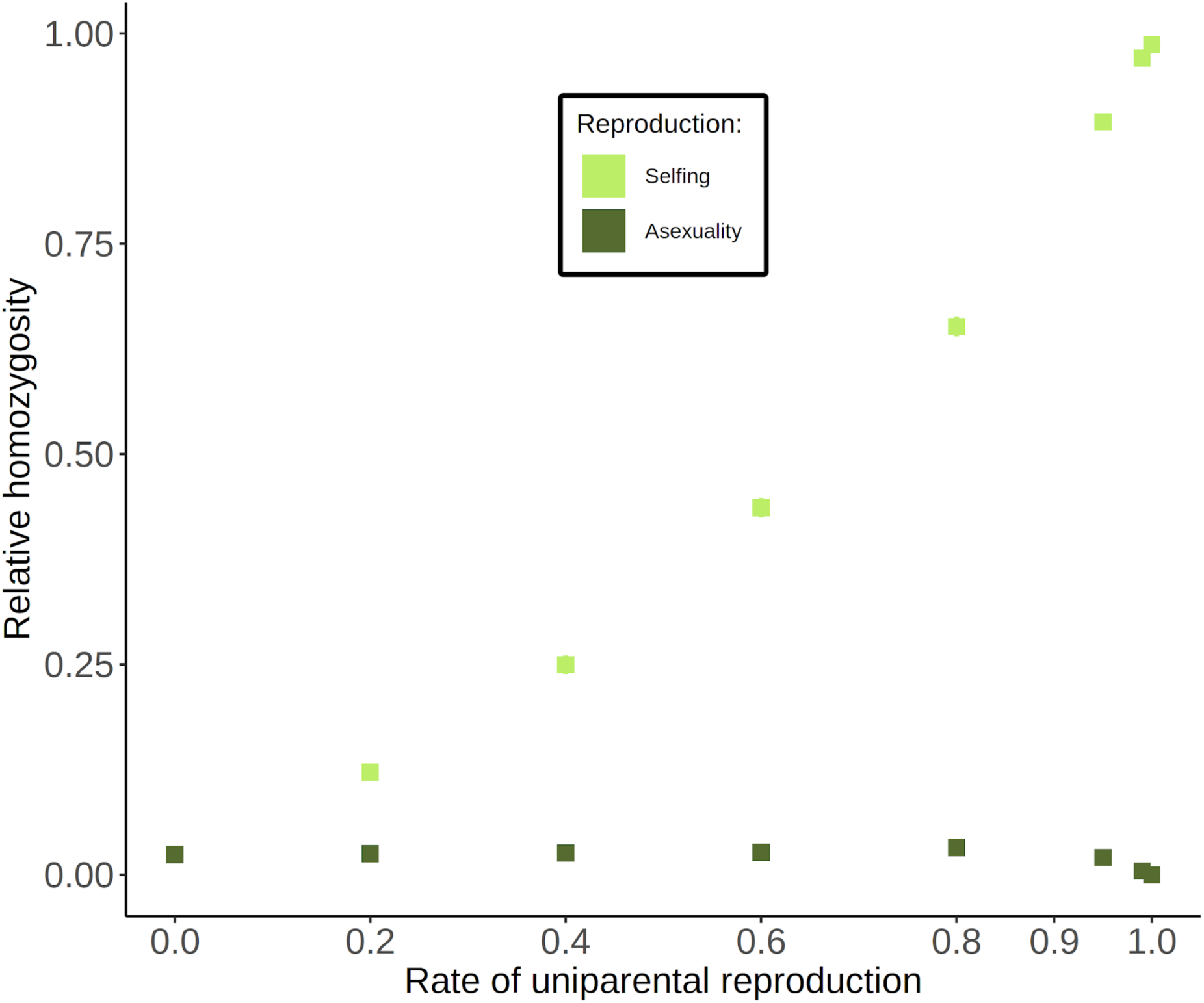
Relative homozygosity as a function of uniparental reproduction rates for fixed s, h. Points are averages across all simulations. Error bars represent 95% confidence intervals; if they are not shown then they lie within a point.

**Figure S6:**
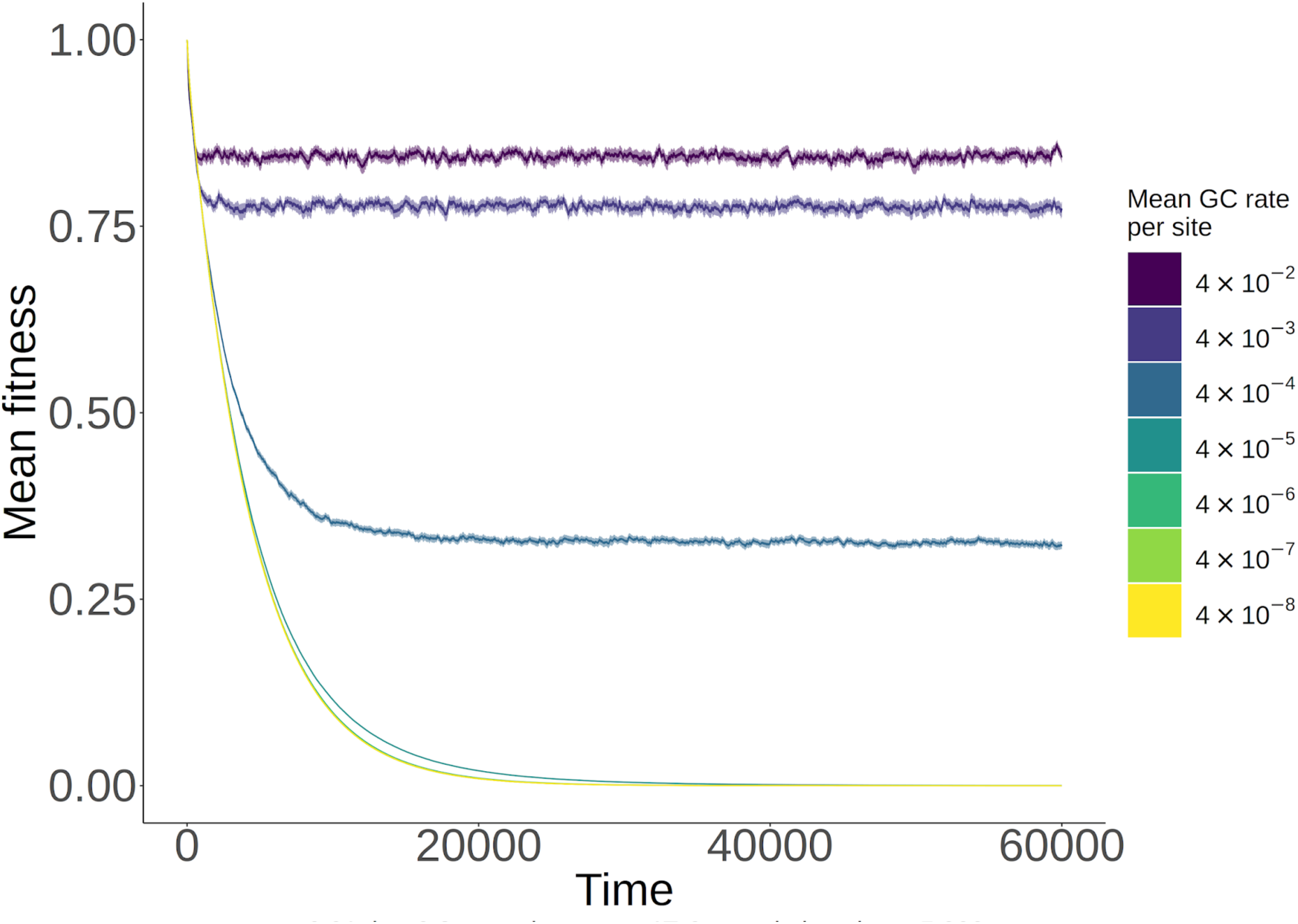
Mean fitness for obligately asexual populations when mitotic gene conversion is present at different rates. Results are shown as a function of time. Thick lines are mean values, while thinner shading around lines show 95% confidence intervals.

**Figure S7:**
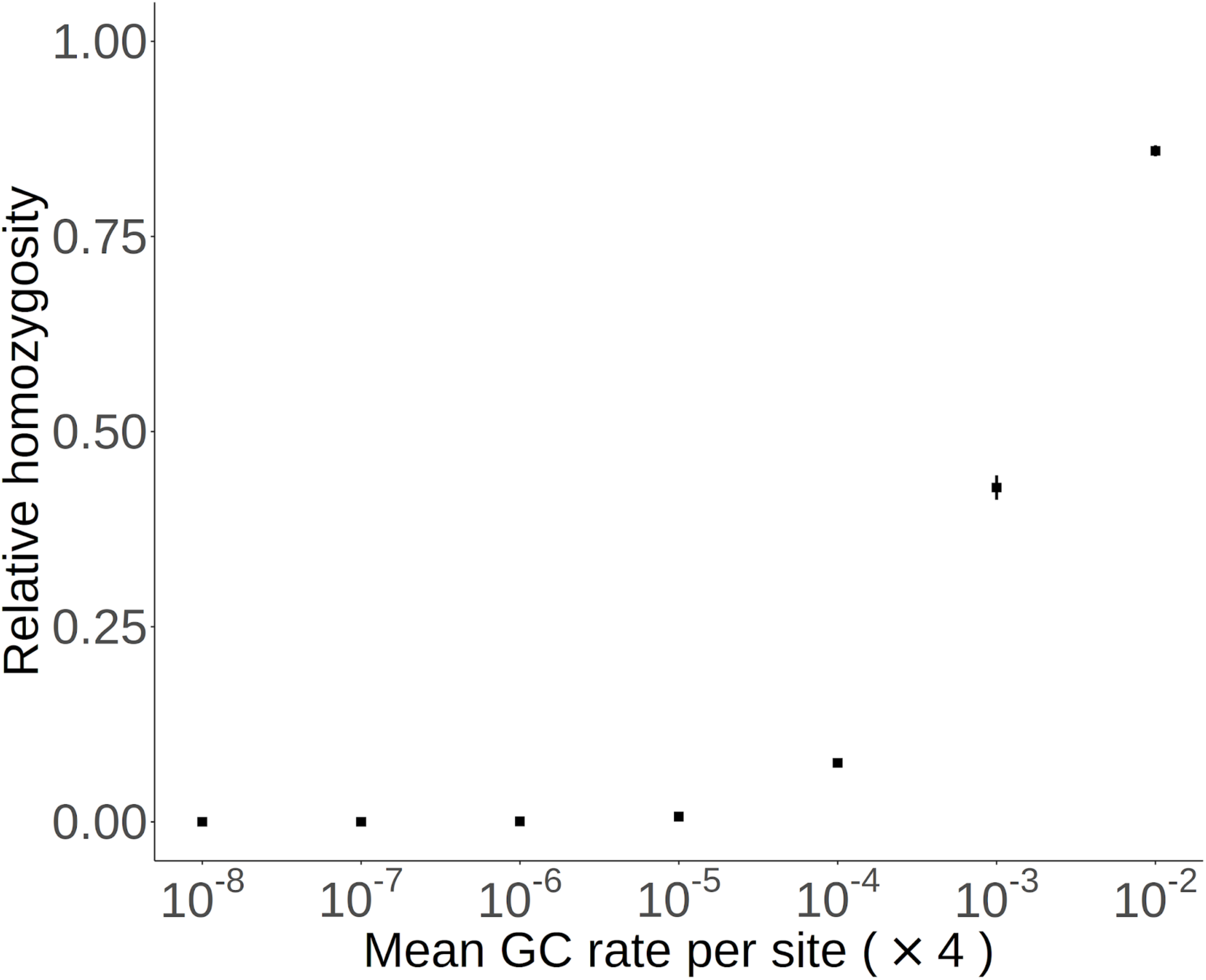
Relative homozygosity for obligate asexual populations with different rates of mitotic gene conversion, for fixed s, h. Points are averages across all simulations. Error bars represent 95% confidence intervals; if they are not shown then they lie within a point.

**Figure S8:**
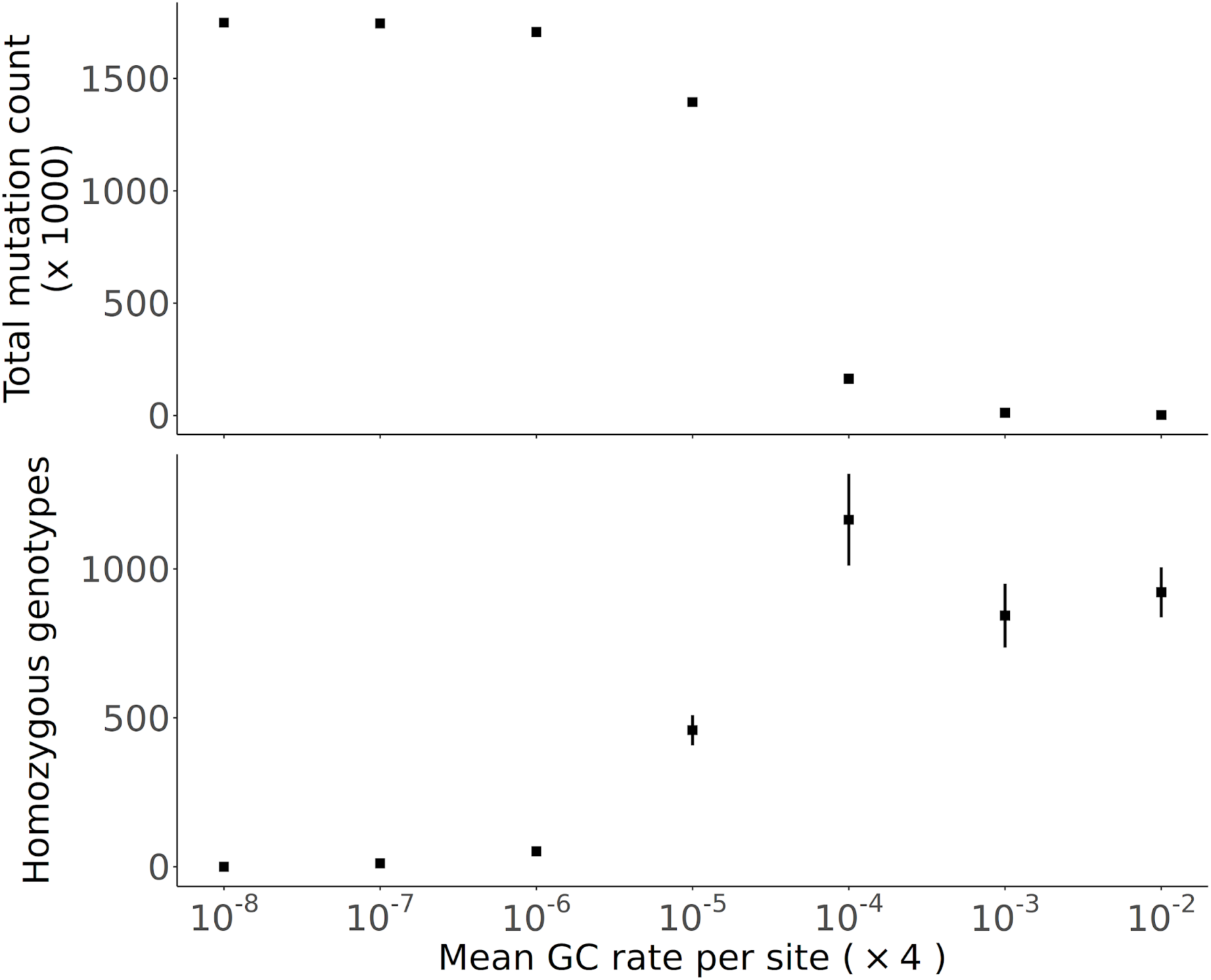
As Figure 6 in the main text but with a population size of N = 1,000, and μ, s increased 5-fold.

**Figure S9:**
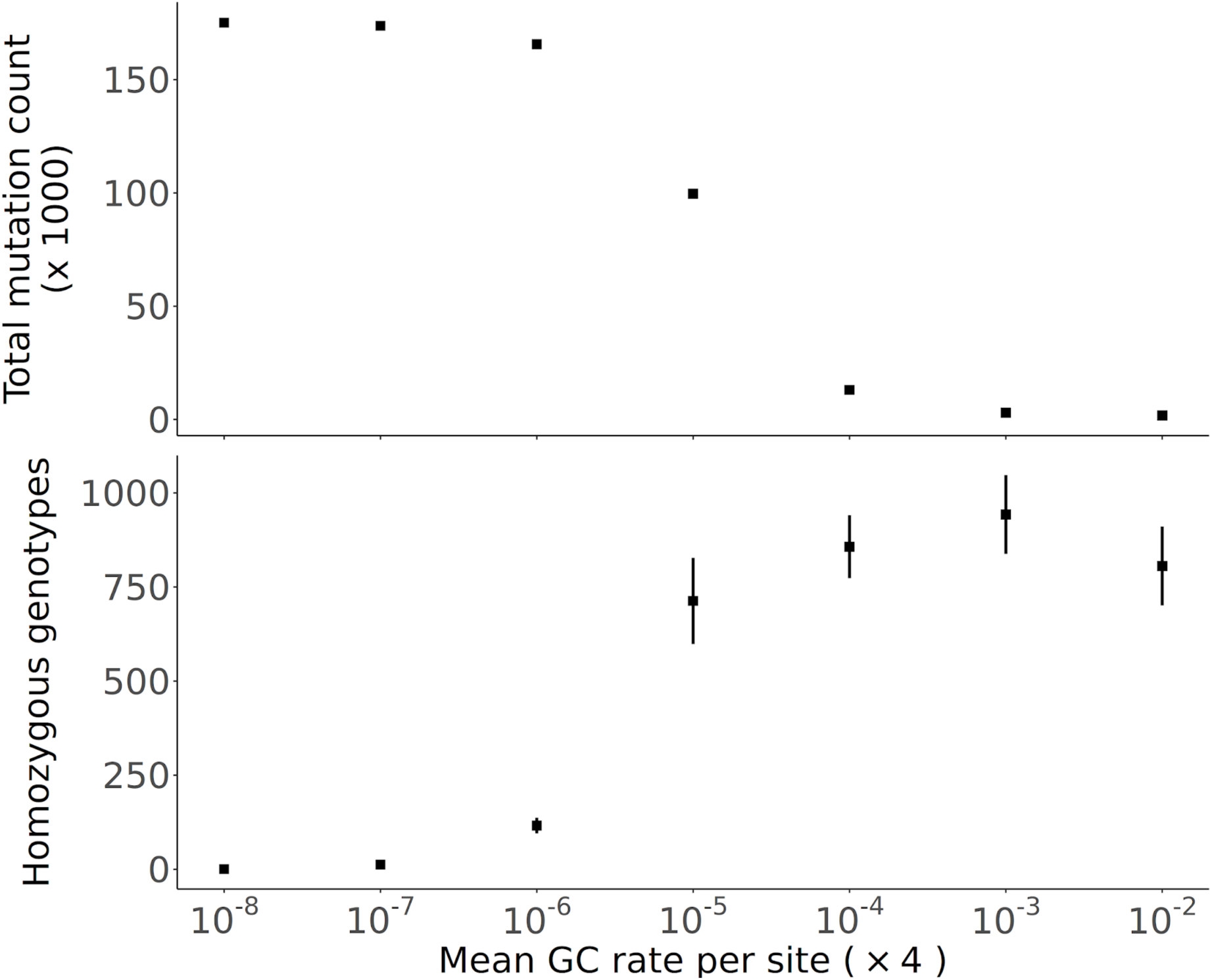
As Figure 6 in the main text but with a population size of N = 10,000, and μ, s decreased 2-fold.

**Figure S10:**
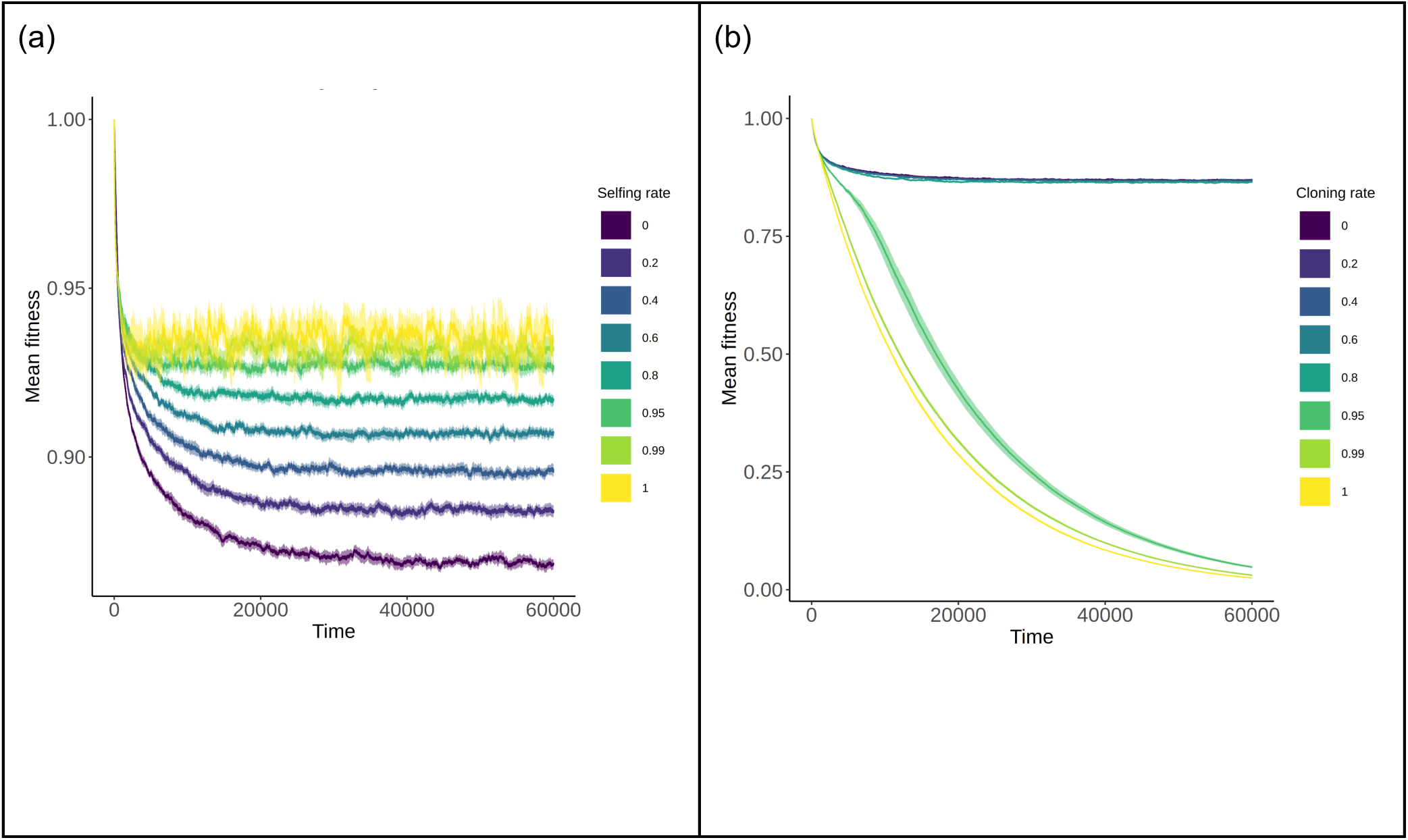
Mean fitness for different modes of uniparental reproduction, when s, h are drawn from a distribution. Results are shown as a function of time, for (a) different rates of self-fertilisation or (b) asexual reproduction under the original WF model. Thick lines are mean values, while thinner shading around lines show 95% confidence intervals. Notice the change of scale on the y-axis between (a) and (b).

**Figure S11:**
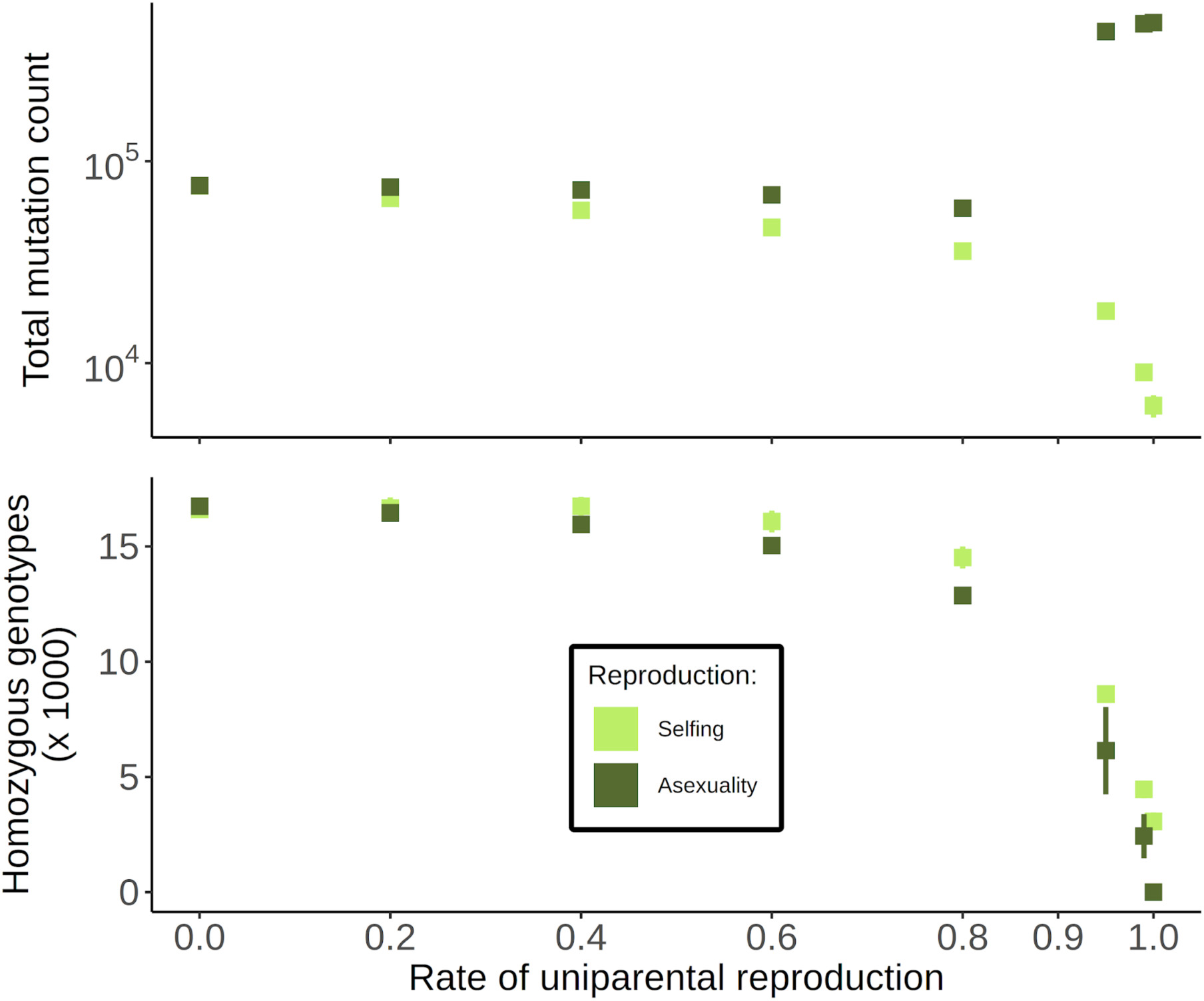
Deleterious mutation counts under different uniparental reproductive modes, when s, h are drawn from a distribution. As a function of the rate of uniparental reproduction (either selfing or asexual reproduction), these plots show (top) the total mutation count present in the sample of 50 individuals, and (bottom) the number of homozygous genotypes present in the sample. Error bars represent 95% confidence intervals; if they are not shown then they lie within a point.

**Figure S12:**
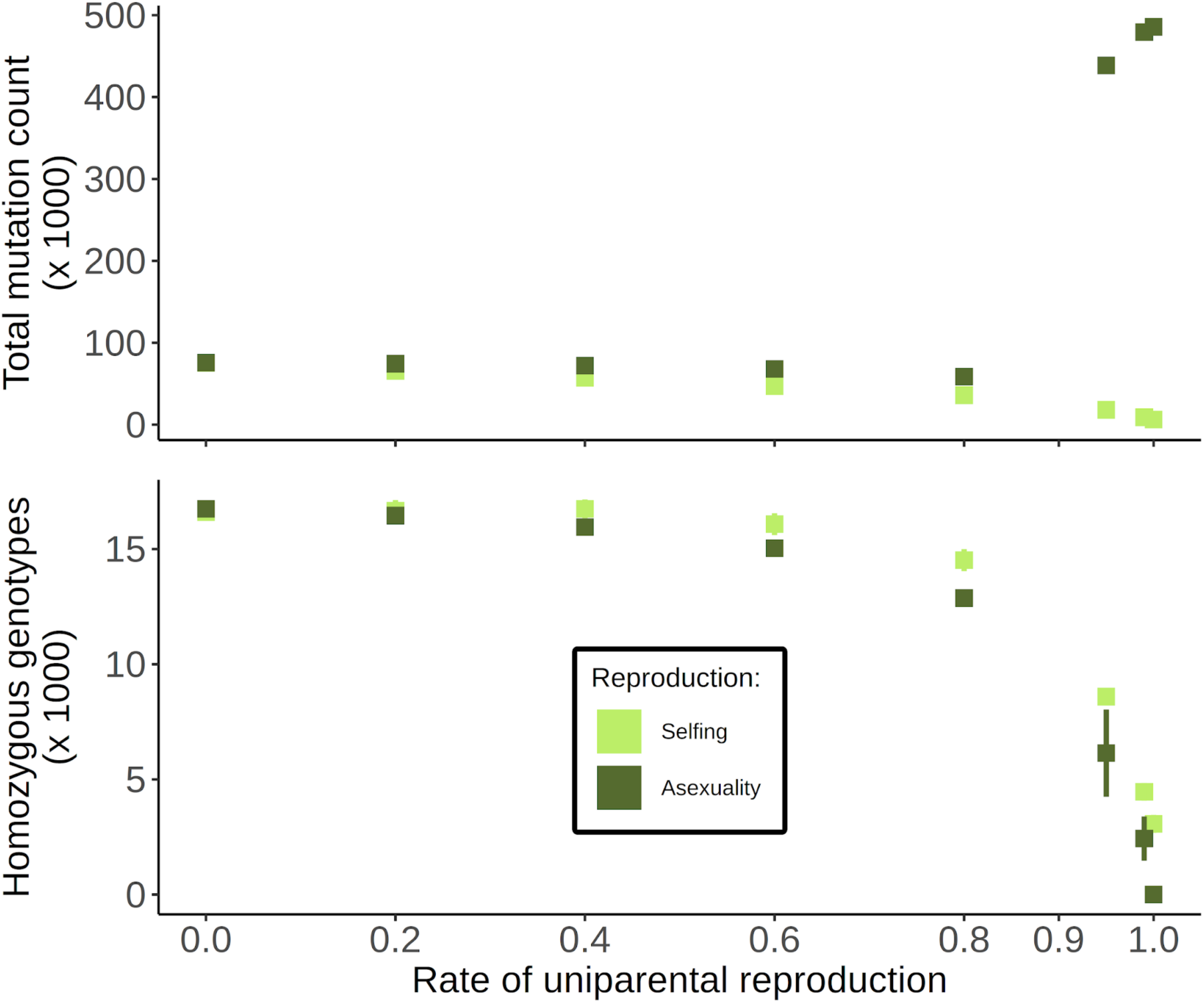
As Figure S11, but with the total mutation count on a linear scale.

**Figure S13:**
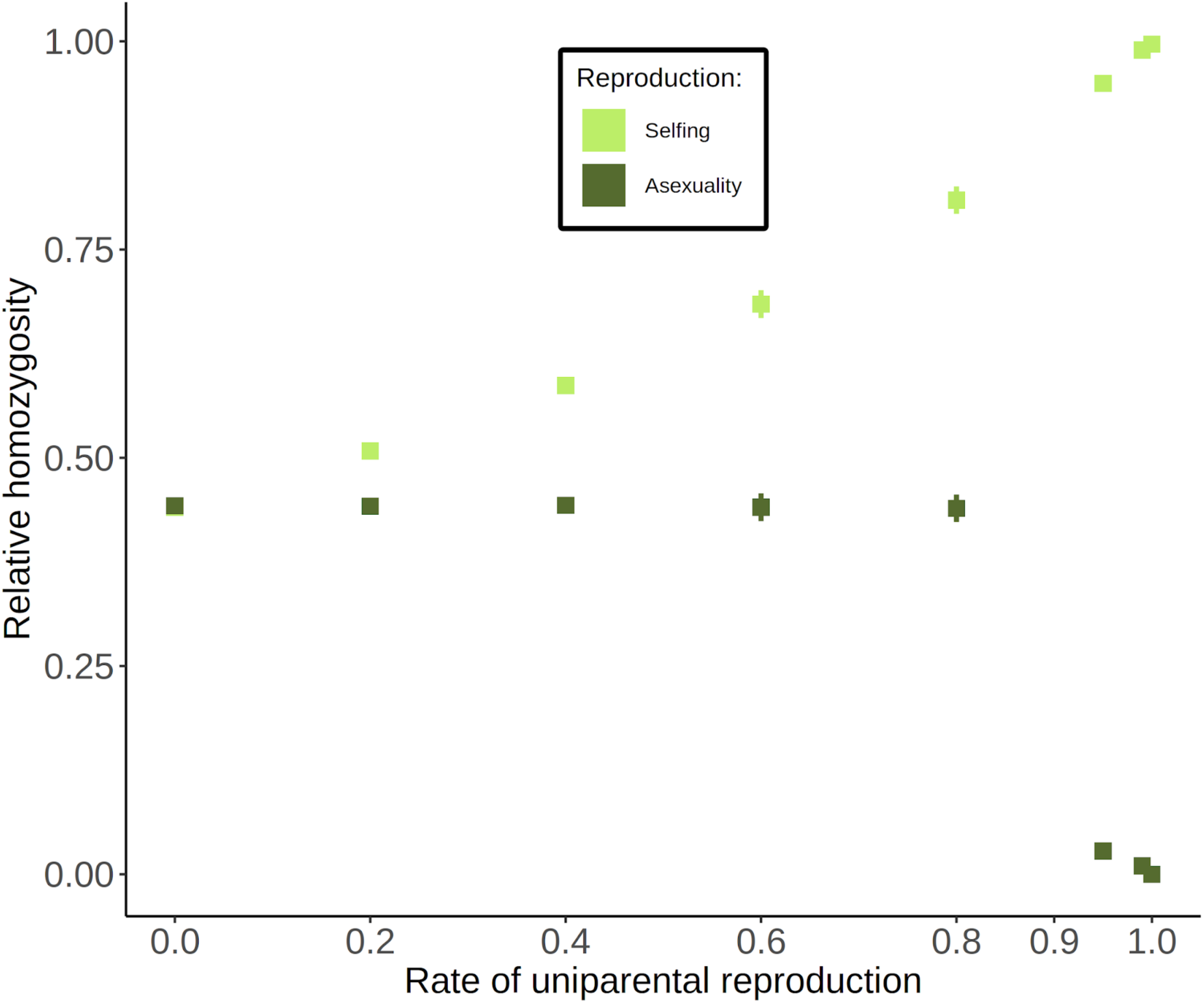
Relative homozygosity as a function of uniparental reproduction rates, when s, h are drawn from a distribution. Points are averages across all simulations. Error bars represent 95% confidence intervals; if they are not shown then they lie within a point.

**Figure S14:**
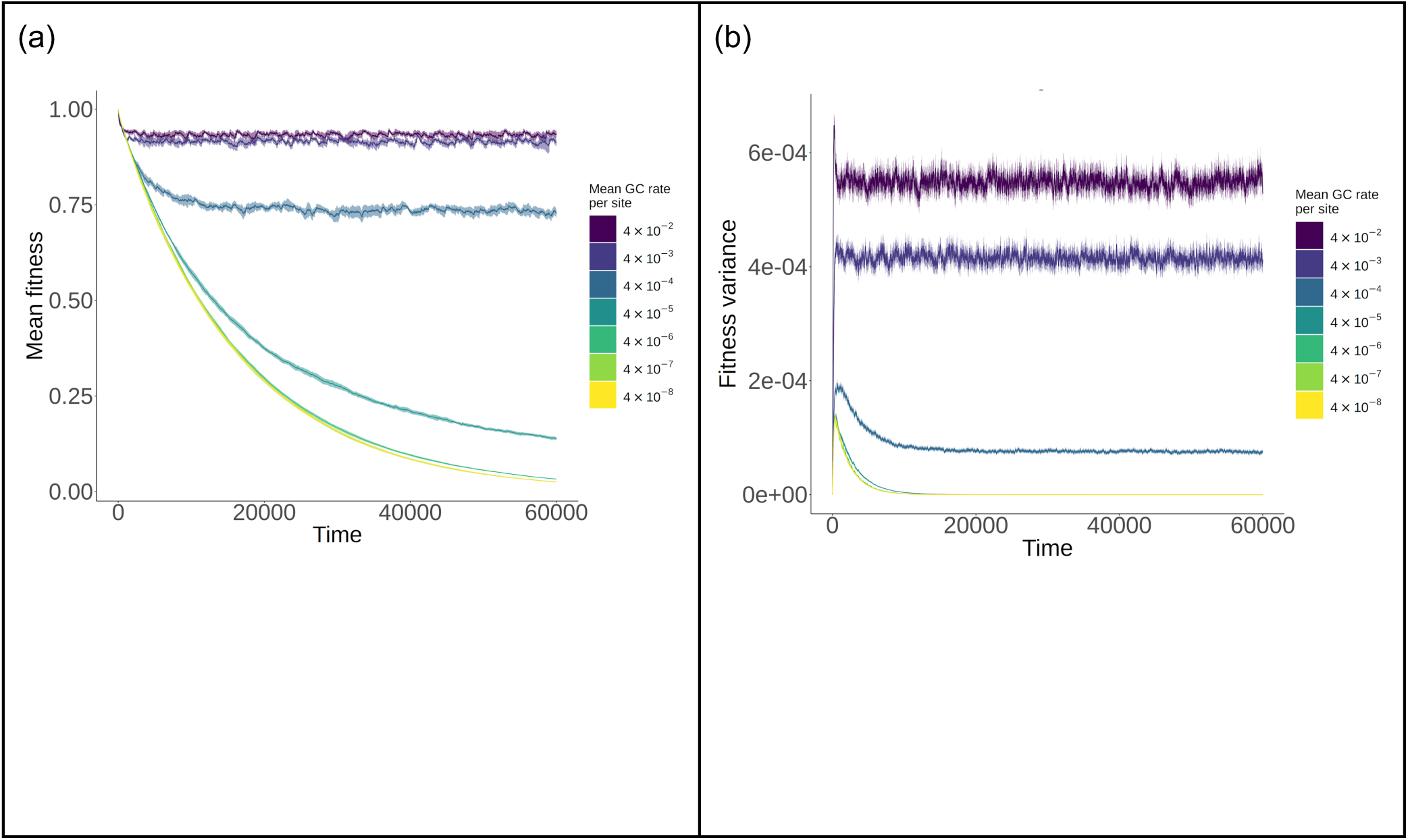
Fitness measurements as a function of time when s, h are drawn from a distribution. Shown are (a) mean fitness, and (b) variance in fitness for simulations of an asexual population subject to different levels of gene conversion. Thick lines are mean values, while thinner shading around lines show 95% confidence intervals.

**Figure S15:**
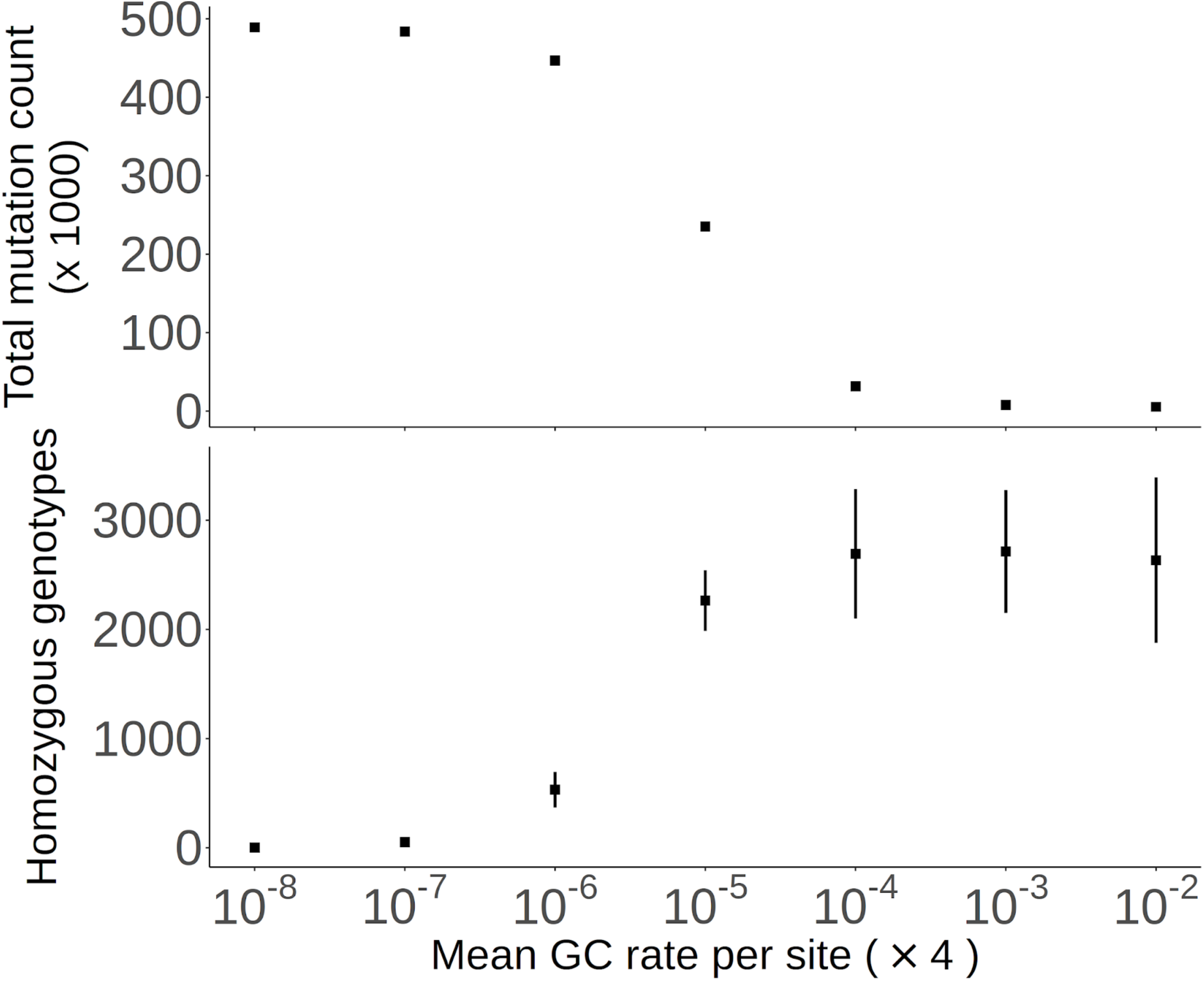
Deleterious mutation counts for obligate asexual populations with mitotic GC present, when s, h are drawn from a distribution. Results are shown as a function of the rate of gene conversion initiation at a single site. Plots show (top) the total mutation count present in the sample of 50 individuals, and (bottom) the number of homozygous genotypes present in the sample. Error bars represent 95% confidence intervals; if they are not shown then they lie within a point.

**Figure S16:**
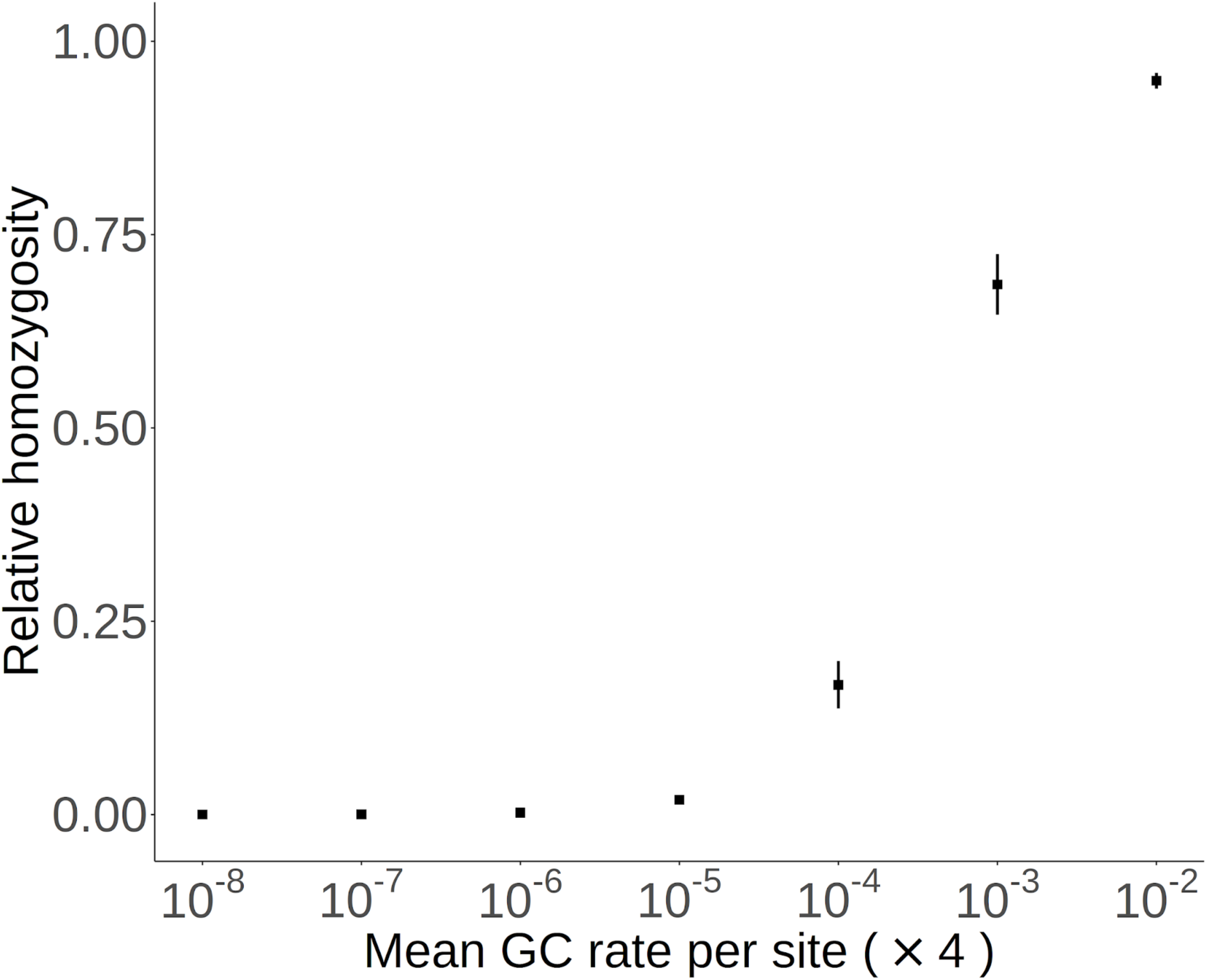
Relative homozygosity for obligate asexual populations with different rates of mitotic gene conversion, when s, h are drawn from a distribution. Points are averages across all simulations. Error bars represent 95% confidence intervals; if they are not shown then they lie within a point.

**Figure S17:**
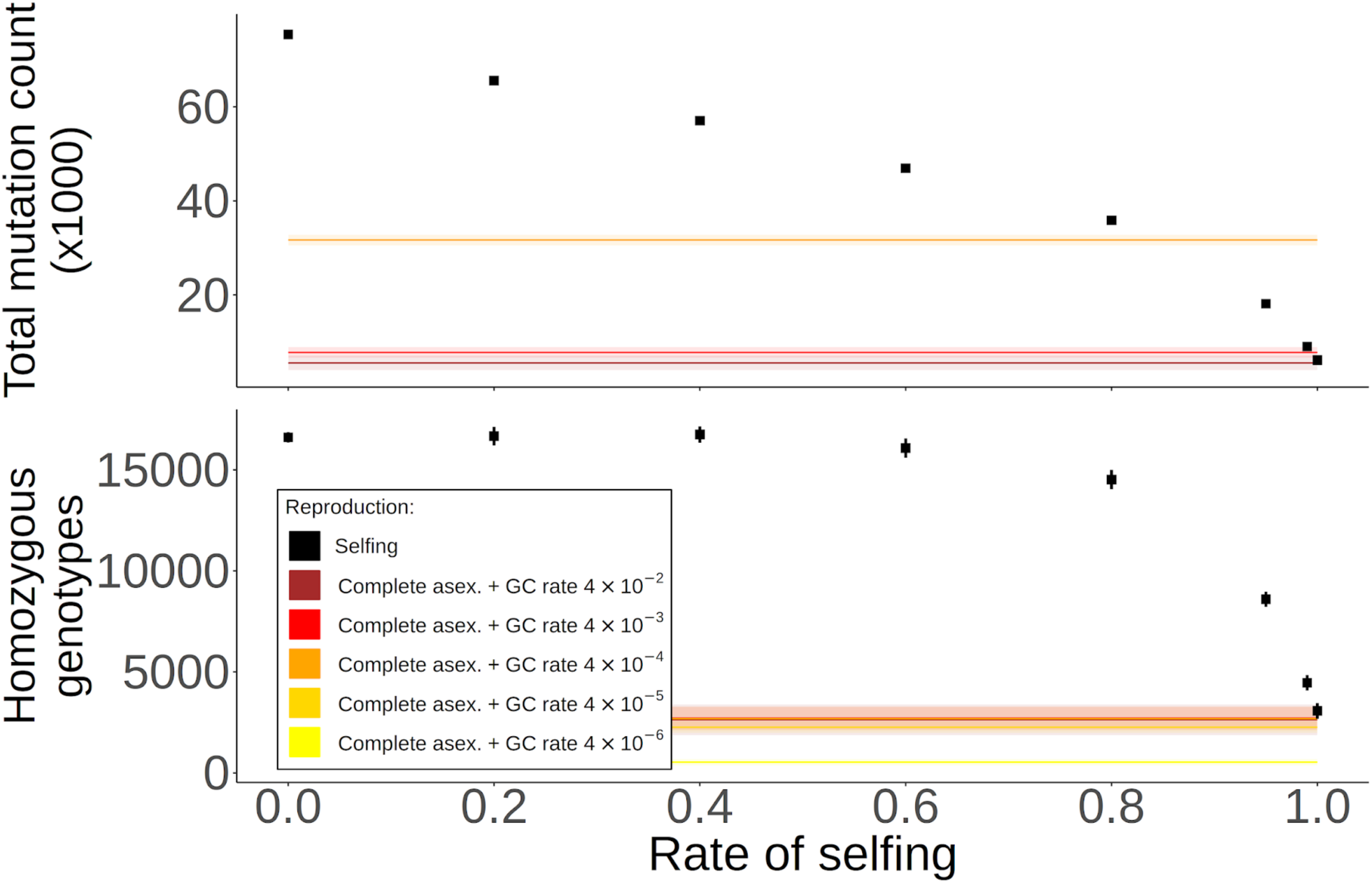
As Figure 7 in the main text, but when s, h are drawn from a distribution.

**Figure S18:**
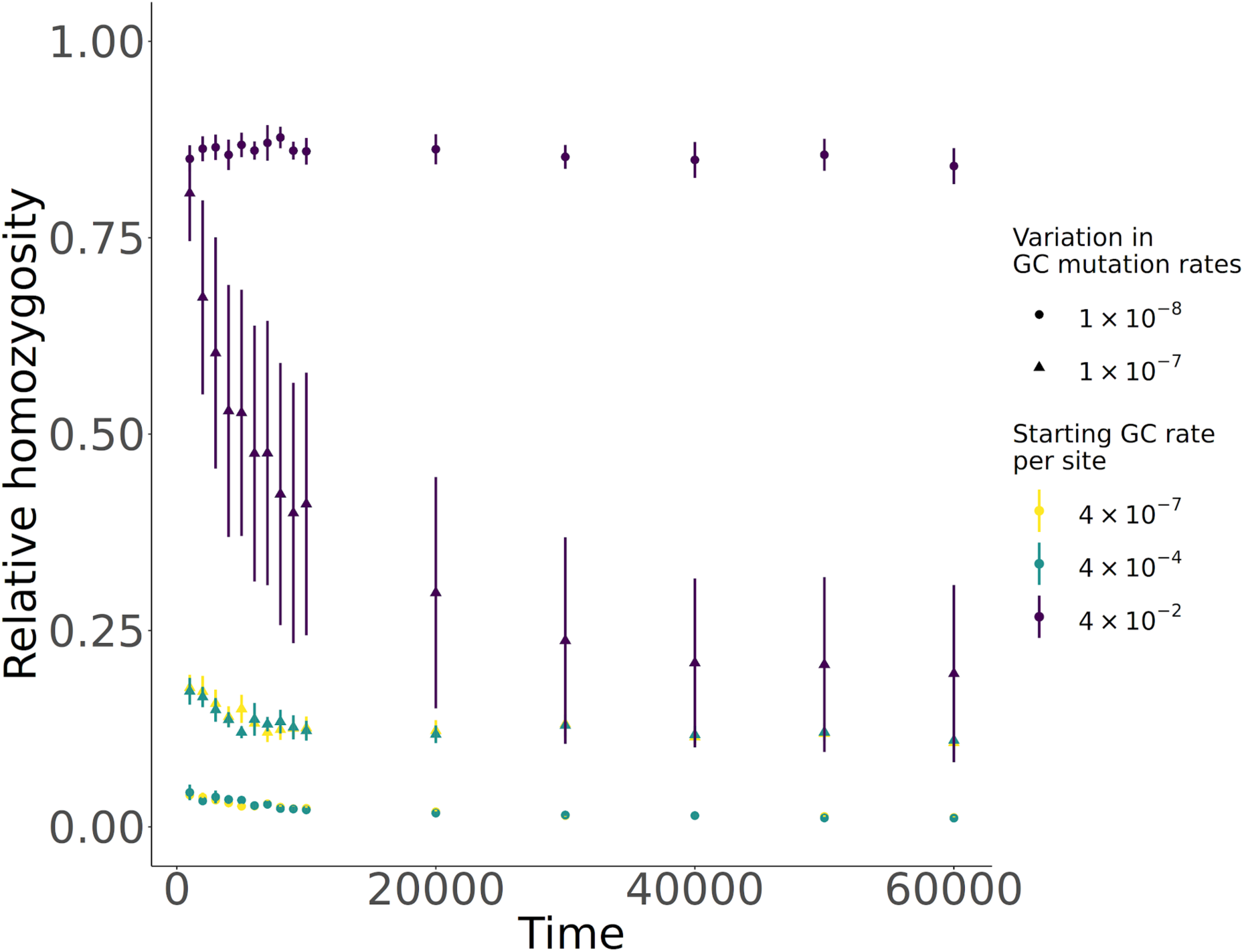
As Figure S7 but when mitotic gene conversion rates are allowed to evolve.

**Figure S19:**
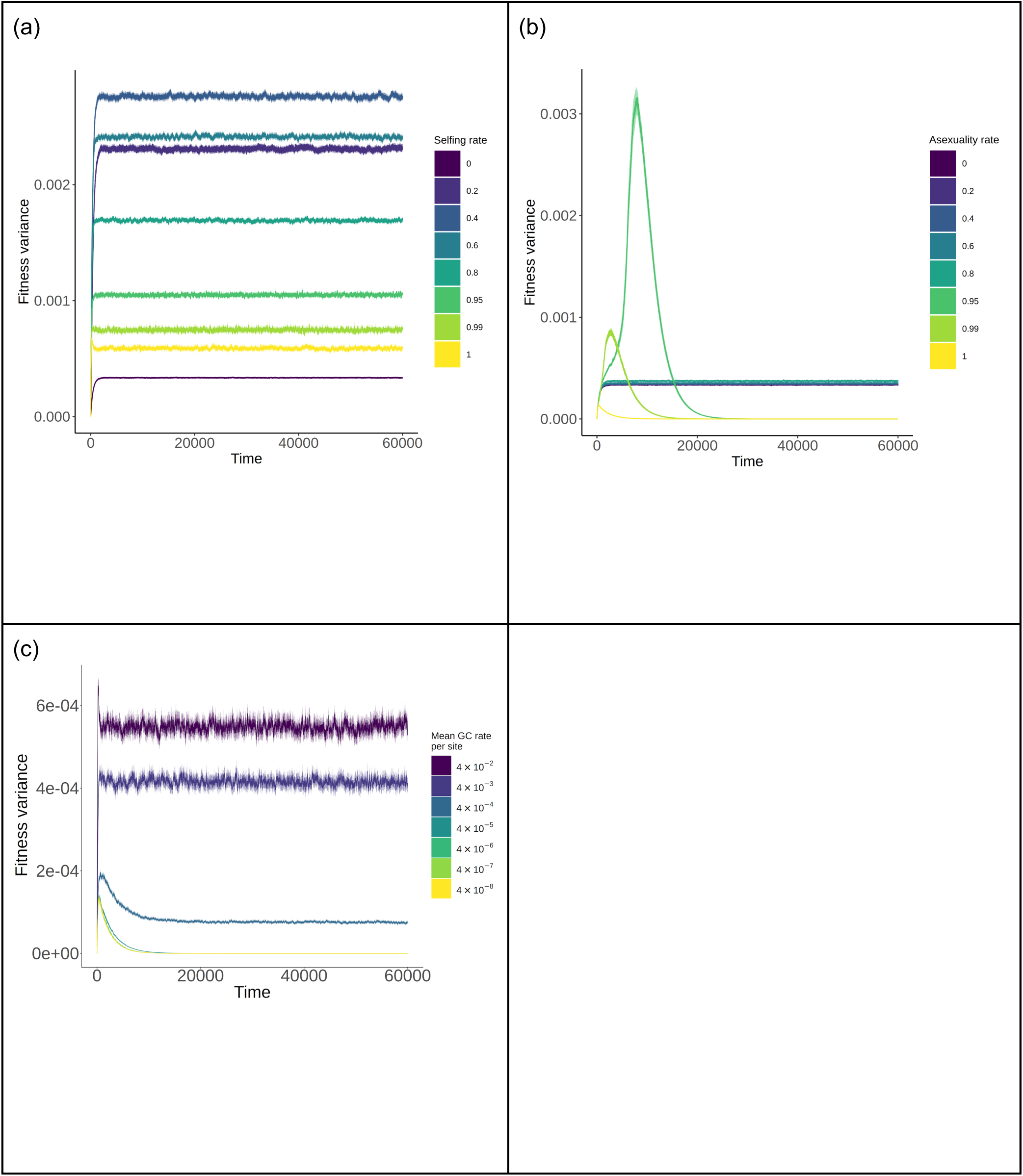
Fitness variance as a function of time, for fixed s, h values. Results are for different levels of (a) self-fertilisation, (b) asexual reproduction for the original Wright-Fisher simulation, or (c) GC in the modified non-WF routine. Thick lines are mean values, while thinner shading around lines show 95% confidence intervals.

**Figure S20:**
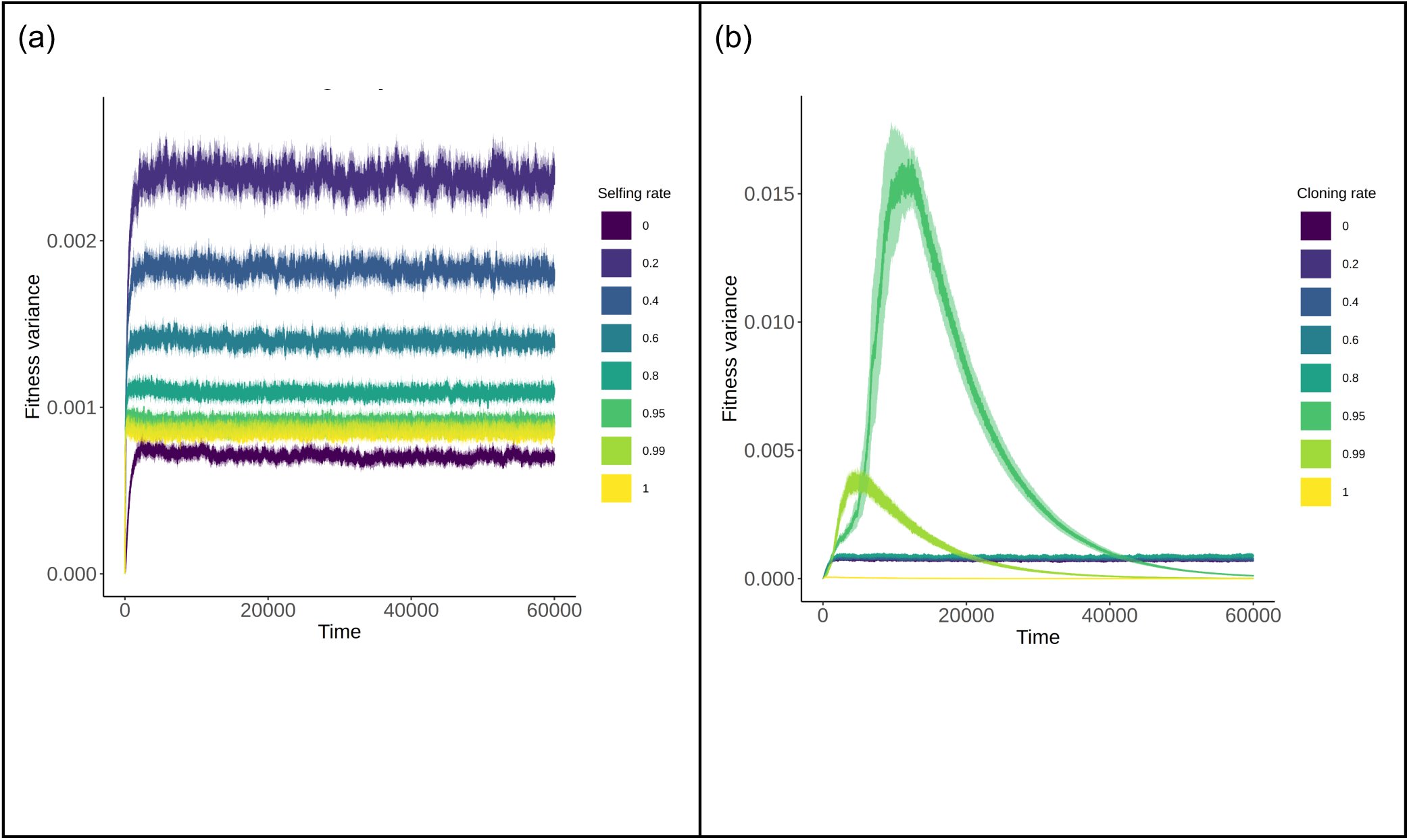
Fitness variance as a function of time when s, h are drawn from a distribution. Results are for different levels of (a) self-fertilisation, (b) asexual reproduction for the original Wright-Fisher simulation. Thick lines are mean values, while thinner shading around lines show 95% confidence intervals.

**Figure S21:**
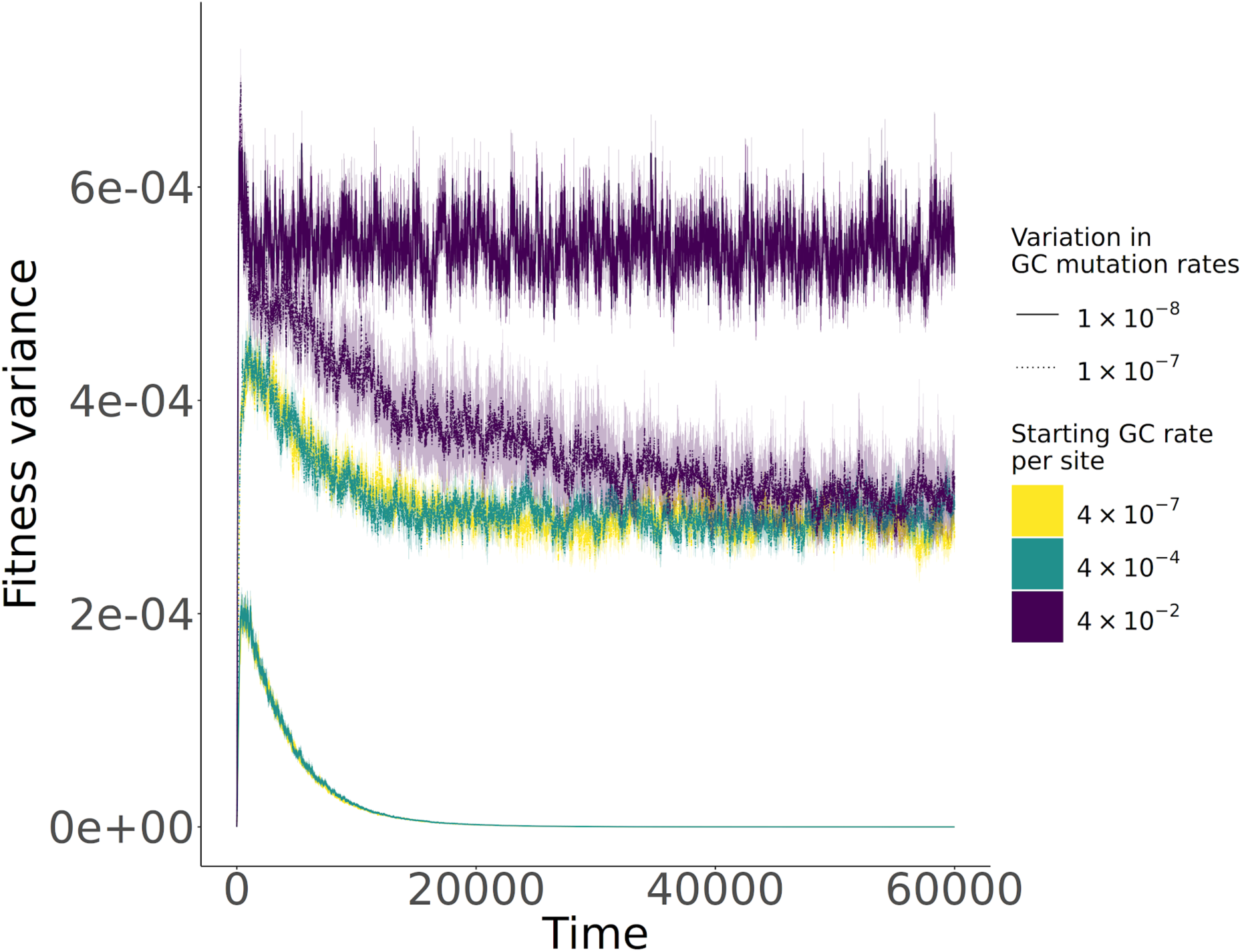
Fitness variance as a function of time, for fixed s, h values and when mitotic gene conversion is allowed to evolve. Thick lines are mean values, while thinner shading around lines show 95% confidence intervals.

**Figure S22:**
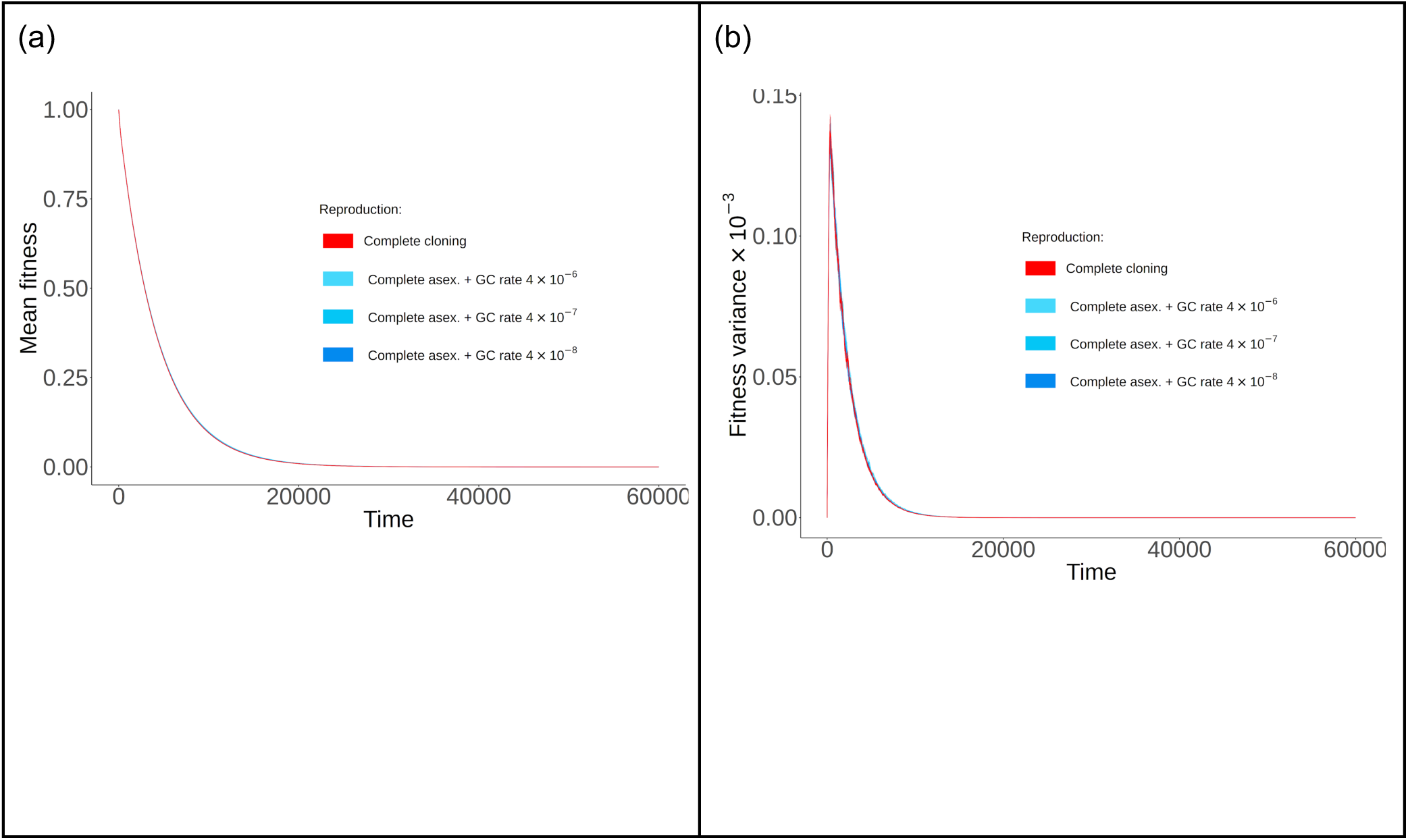
Comparing fitness measurements between the original WF model, and the new routine that implements GC into asexuals. Plots show either (a) mean fitness or (b) fitness variance as a function of time. They compare these measurements under complete cloning in the WF model, and in the new non-WF-based routine under low levels of GC. Plots show both the mean value and 95% CI, although the latter are too small to see.

**Figure S23:**
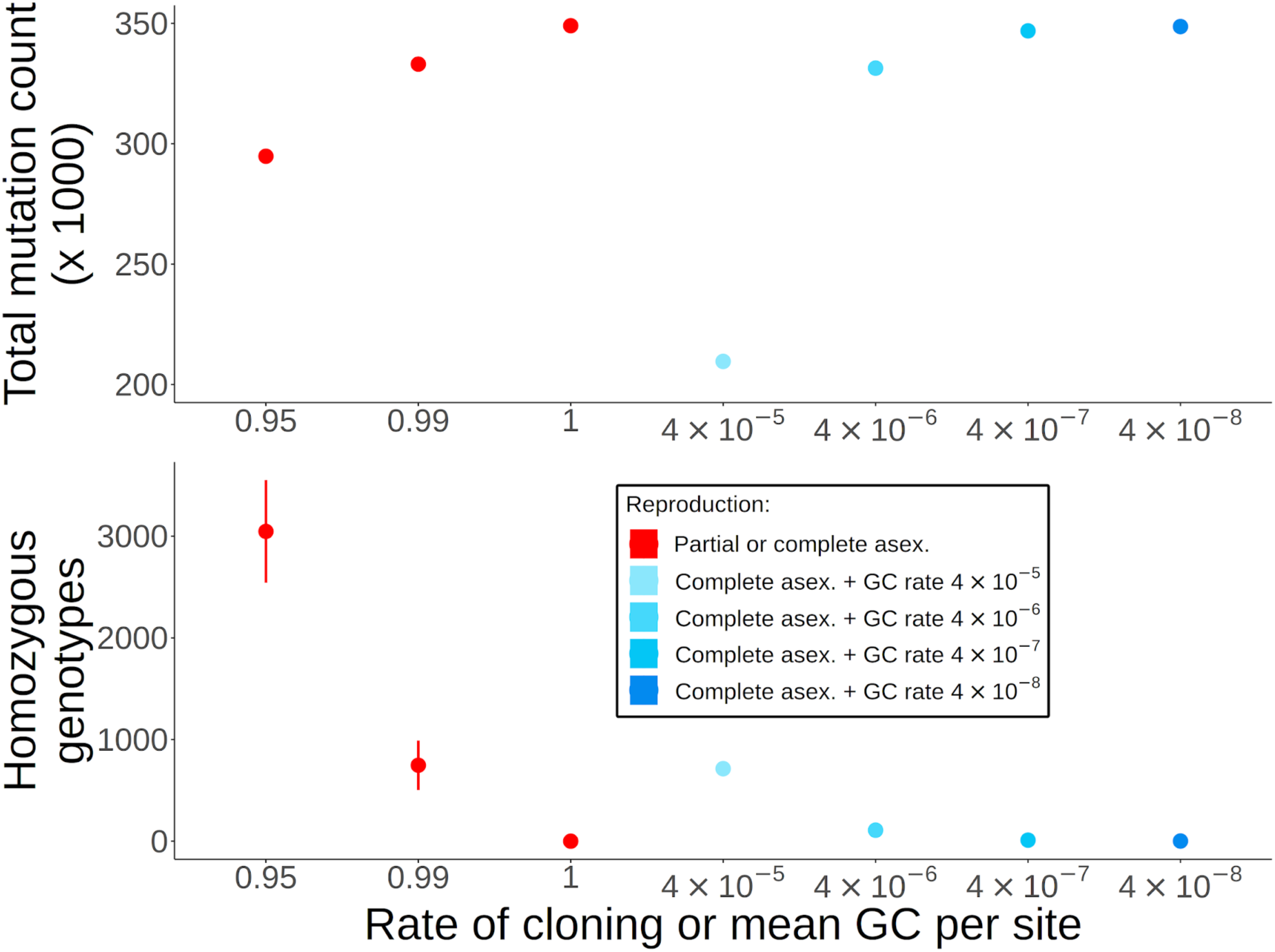
Comparing mutation accumulation between the original WF model, and the new routine that implements GC into asexuals. Plots show either (top) total mutation count, or (bottom) number of homozygous genotypes. Plots compare high levels of asexuality under the original WF model (red points), to results obtained under the new non-WF-based GC routine (blue points). Bars represent 95% confidence intervals; if they are not seen then they lie within the point. Note that the total mutation counts and homozygous genotype counts under complete cloning in the original WF simulations (red dots, left-hand panels) match up with those for the new non-WF routine when gene conversion is low (blue dots, right-hand panels), demonstrating the equivalence of the two routines.

**Figure S24:**
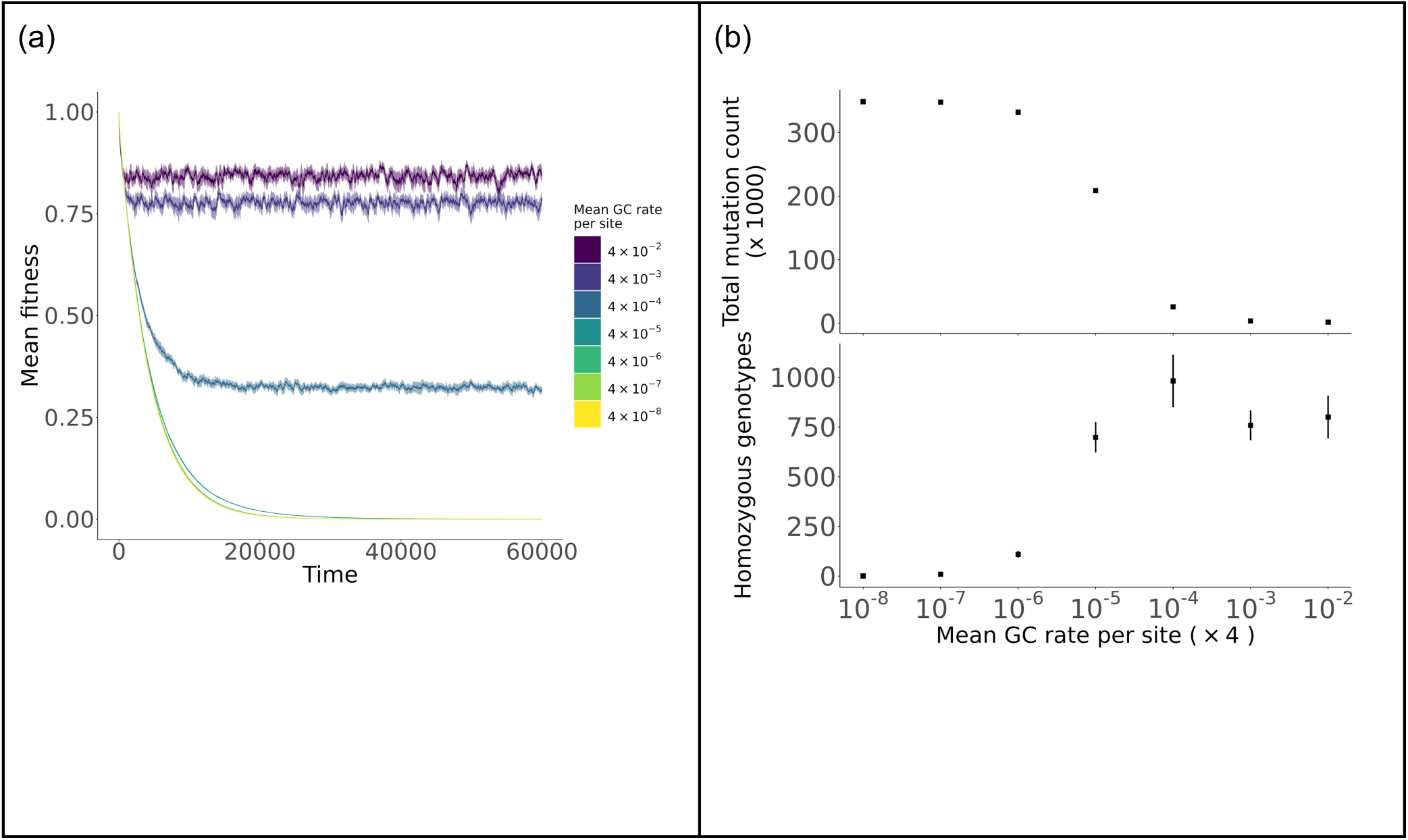
Testing the evolving gene conversion simulation. Results show (a) the mean fitness as a function of time, and (b) mutation counts as a function of the mean gene conversion rate. Bands, lines are 95% confidence intervals. Compare with Figures S6 and 6 in the main text, respectively.

