## Supplementary Mathematica File for "High levels of mitotic gene conversion are needed to effectively purge deleterious mutations in asexual organisms": Supplementary_Mathematica_File.pdf

---

#### Section A: Single-locus model, mutation-selection balance under facultative sex and gene conversion

In[ ]:=
```
(* This function clears existing commands from memory *)
Clear["`*"];
```

##### Recursions for asex with gene conversion, and selfing

This is a simple model for change in deleterious allele frequencies at a single locus, if affected by partial sex, mutation and gene conversion.
Let a be the wild-type and A the deleterious site.
Denote gaa, gAa, gAA the frequencies of aa, Aa and AA genotypes.
Note that gaa + gAa + gAA = 1.

###### Change in frequencies by selection, assuming mutations are deleterious

In[ ]:=
```
(* Individual fitness defintions *)
Waa = 1;
WAa = 1 - h s;
WAA = 1 - s;
(* Mean fitness *)
Wm = gaa * Waa + gAa * WAa + gAA * WAA
```

Out[ ]=
```
gaa + gAA (1 - s) + gAa (1 - h s)
```

In[ ]:=
```
gaas = gaa * Waa / Wm;
gAas = gAa * WAa / Wm;
gAAs = gAA * WAA / Wm;
```

Checking post-selection frequencies sum to 1:

In[ ]:=
```
gaas + gAas + gAAs // Simplify
```

Out[ ]=
```
1
```

#### Reproduction *with selfing*

Via sex (outcrossing):

In[ ]:=

$$gaaO = \left(gaas + \frac{gAas}{2}\right)^2;$$
$$gAaO = 2\left(gaas + \frac{gAas}{2}\right)\left(gAAs + \frac{gAas}{2}\right);$$
$$gAAO = \left(gAAs + \frac{gAas}{2}\right)^2;$$

Via selfing:

In[ ]:=

$$gaaS = gaas + \frac{gAas}{4};$$
$$gAaS = \frac{gAas}{2};$$
$$gAAS = gAAs + \frac{gAas}{4};$$

Total reproductive frequencies, assuming a mix of outcrossing and selfing (where S = frequency of selfing):

In[ ]:=
```
gaaRS = (1 - S) gaaO + S gaaS;
gAaRS = (1 - S) gAaO + S gAaS;
gAARS = (1 - S) gAAO + S gAAS;
```

Checking if summing to one:

In[ ]:=
```
gaaRS + gAaRS + gAARS // Simplify
```

Out[ ]=
```
1
```

#### Reproduction *via facultative sex*

Via asex (clonal reproduction):

In[ ]:=
```
gaaA = gaas;
gAaA = gAas;
gAAA = gAAs;
```

Total reproductive frequencies, assuming a mix of sex and asex (let $\sigma$ = frequency of sex):

```
In[•]:= gaaRA = σ gaa0 + (1 - σ) gaaA;
gAaRA = σ gAa0 + (1 - σ) gAaA;
gAARA = σ gAA0 + (1 - σ) gAAA;
```

Checking if summing to one:

```
In[•]:= gaaRA + gAaRA + gAARA // Simplify
Out[•]=
1
```

#### Mutation

Assume unidirectional mutations, so a -> A with probability $\mu$ and no back-mutation probability.
Hence aa -> aA with probability $2\mu(1-\mu)$, aa -> AA with probability $\mu^2$, and aA -> AA with probability $\mu$.
This yields the following recursions:

```
In[•]:= gaaMS = (1 - μ)^2 gaaRS;
gAaMS = 2 μ (1 - μ) gaaRS + (1 - μ) gAaRS;
gAAMS = gAARS + μ^2 gaaRS + μ gAaRS;
```

Checking if summing to one:

```
In[•]:= gaaMS + gAaMS + gAAMS // Simplify
Out[•]=
1
```

Same but for facultative sex:

```
In[•]:= gaaMA = (1 - μ)^2 gaaRA;
gAaMA = 2 μ (1 - μ) gaaRA + (1 - μ) gAaRA;
gAAMA = gAARA + μ^2 gaaRA + μ gAaRA;
```

```
In[•]:= gaaMA + gAaMA + gAAMA // Simplify
Out[•]=
1
```

#### Gene conversion

Gene conversion acts with probability $\gamma$.
It only affects heterozygotes, and turns them into homozygotes with probability $\gamma$ (each homozygote with probability 1/2).
First for selfing:

```
In[•]:= gaaGCS = gaaMS + γ/2 gAaMS;
gAaGCS = (1 - γ) gAaMS;
gAAGCS = gAAMS + γ/2 gAaMS;
```

Checking if summing to one:

```
In[•]:= gaaGCS + gAaGCS + gAAGCS // Simplify
Out[•]=
1
```

Next for facultative sex:

In[∘]:=
```
gaaGCA = gaaMA + γ/2 gAaMA;
gAaGCA = (1 - γ) gAaMA;
gAAGCA = gAAMA + γ/2 gAaMA;
```

In[∘]:=
```
gaaGCA + gAaGCA + gAAGCA // Simplify
```

Out[∘]=
```
1
```

### Steady-state solutions

#### Analytical solutions

This function converts genotype frequencies to allele frequencies while also considering inbreeding (F):

In[∘]:=
```
GtoP = {gaa → (1 - pA)^2 + pA (1 - pA) F,
   gAa → 2 pA (1 - pA) (1 - F), gAA → pA^2 + pA (1 - pA) F}
```

Out[∘]=
```
{gaa → (1 - pA)^2 + F (1 - pA) pA, gAa → 2 (1 - F) (1 - pA) pA, gAA → F (1 - pA) pA + pA^2}
```

We look for steady-state frequencies of pA and F, assuming small $\mu$ and pA. That way, we can use a series expansion to simplify the recursions while capturing the main effects of the reproductive mode and gene conversion.

Here's the appropriate expansion for pA following a single generation, given partial selfing:

In[∘]:=
```
Series[gAAGCS + gAaGCS/2 /. GtoP /. {pA → pA * ξ, μ → μ * ξ}, {ξ, 0, 1}] // Simplify //
   Normal;
% /. ξ → 1 // Simplify
```

Out[∘]=
```
pA + F (-1 + h) pA s - h pA s + μ
```

We obtain the same result under facultative sex:

In[∘]:=
```
Series[gAAGCA + gAaGCA/2 /. GtoP /. {pA → pA * ξ, μ → μ * ξ}, {ξ, 0, 1}] // Simplify //
   Normal;
% /. ξ → 1 // Simplify
```

Out[∘]=
```
pA + F (-1 + h) pA s - h pA s + μ
```

This gives us a system of equations for the change in frequency of pA; denote it pA'.
Hence we can write a general solution for the steady-state frequency of the deleterious allele (i.e. pA' - pA = 0), given an inbreeding coefficient F:

In[•]:=
```
Solve[pA + F (-1 + h) pA s - h pA s + μ - pA == 0, pA]
```
Out[•]=

$$\left\{\left\{pA \to -\frac{\mu}{(-F - h + F\,h)\,s}\right\}\right\}$$

Rewriting:

$$\frac{\mu}{(F + h - F\,h)\,s}$$

This equation matches previous results for allele frequencies under non-random mating (e.g. Glémin 2003).

Below are the equations for change in F following a single generation, assuming first selfing, and subsequetly for facultative sex:

In[•]:=
```
Series[(gAAGCS gaaGCS - (gAaGCS/2)^2)/((gAAGCS + gAaGCS/2) (gaaGCS + gAaGCS/2)) /. GtoP /. {pA → pA * ξ, μ → μ * ξ},
    {ξ, 0, 0}] // Simplify // Normal;
% /. ξ → 1 // Simplify
```
Out[•]=

$$\frac{-pA\,(1 + F + F\,(-2 + h)\,s - h\,s)\,S\,(-1 + \gamma) + 2\,pA\,(1 + F\,(-1 + h)\,s - h\,s)\,\gamma + 2\,\gamma\,\mu}{2\,(pA + F\,(-1 + h)\,pA\,s - h\,pA\,s + \mu)}$$

In[•]:=
```
Series[(gAAGCA gaaGCA - (gAaGCA/2)^2)/((gAAGCA + gAaGCA/2) (gaaGCA + gAaGCA/2)) /. GtoP /. {pA → pA * ξ, μ → μ * ξ},
    {ξ, 0, 0}] // Simplify // Normal;
% /. ξ → 1 // Simplify
```
Out[•]=

$$\frac{\gamma\,(pA - h\,pA\,s + \mu) + F\,pA\,((-1 + \gamma)\,(-1 + \sigma) + s\,(-1 + h\,\gamma + \sigma - \gamma\,\sigma))}{pA + F\,(-1 + h)\,pA\,s - h\,pA\,s + \mu}$$

Now we can simultaneously solve for pA’ - pA = 0 and F’ - F = 0 to find the inbreeding coefficient under each mating-system. First for selfing:

In[•]:=
```
Solve[{(pA + F (-1 + h) pA s - h pA s + μ) - pA == 0,
    ((-pA (1 + F + F (-2 + h) s - h s) S (-1 + γ) + 2 pA (1 + F (-1 + h) s - h s) γ + 2 γ μ)/(2 (pA + F (-1 + h) pA s - h pA s + μ))) - F == 0},
  {pA, F}] // Simplify
```
Out[•]=

$$\left\{\left\{pA \to -\frac{(2 + (1 + (-2 + h)\,s)\,S\,(-1 + \gamma))\,\mu}{s\,(S\,(-1 + \gamma) + h\,(2 + (-2 + s)\,S)\,(-1 + \gamma) - 2\,\gamma)},\ F \to \frac{(-1 + h\,s)\,S\,(-1 + \gamma) + 2\,\gamma}{2 + (1 + (-2 + h)\,s)\,S\,(-1 + \gamma)}\right\}\right\}$$

Rewriting the pA, F terms to look nicer:

In[ ]:=
```
(2 - (1 - (2 - h) s) S (1 - γ)) μ / (s (S (1 - γ) + h (2 - (2 - s) S) (1 - γ) + 2 γ)) -
  {-(2 + (1 + (-2 + h) s) S (-1 + γ)) μ / (s (S (-1 + γ) + h (2 + (-2 + s) S) (-1 + γ) - 2 γ))} // Simplify
```
Out[ ]=
```
{0}
```

In[ ]:=
```
((1 - h s) S (1 - γ) + 2 γ) / (2 - (1 - (2 - h) s) S (1 - γ)) - {((-1 + h s) S (-1 + γ) + 2 γ) / (2 + (1 + (-2 + h) s) S (-1 + γ))} // Simplify
```
Out[ ]=
```
{0}
```

Note we can also write pA as $\frac{\mu}{(F+h-F h) s}$ but with F equal to the solution above:

In[ ]:=
```
(2 - (1 - (2 - h) s) S (1 - γ)) μ / (s (S (1 - γ) + h (2 - (2 - s) S) (1 - γ) + 2 γ)) -
  {μ / ((F + h - F h) s) /. F → ((1 - h s) S (1 - γ) + 2 γ) / (2 - (1 - (2 - h) s) S (1 - γ))} // Simplify
```
Out[ ]=
```
{0}
```

Next we find the steady-state values of pA, F under facultative sex:

In[ ]:=
```
Solve[{(pA + F (-1 + h) pA s - h pA s + μ) - pA == 0,
   ((γ (pA - h pA s + μ) + F pA ((-1 + γ) (-1 + σ) + s (-1 + h γ + σ - γ σ))) / (pA + F (-1 + h) pA s - h pA s + μ)) - F == 0}, {pA,
   F}] // Simplify
```
Out[ ]=

$$\left\{\left\{pA \to \frac{\mu\ (\gamma + s\ (-1+\gamma)\ (-1+\sigma) + \sigma - \gamma\,\sigma)}{s\ (\gamma + h\ (-1+\gamma)\ (s\ (-1+\sigma) - \sigma))},\ F \to \frac{\gamma}{\gamma + s\ (-1+\gamma)\ (-1+\sigma) + \sigma - \gamma\,\sigma}\right\}\right\}$$

Rewriting the pA, F terms to look nicer:

In[ ]:=
```
μ (γ + (1 - γ) (s (1 - σ) + σ)) / (s (γ + h (1 - γ) (s (1 - σ) + σ))) - {μ (γ + s (-1 + γ) (-1 + σ) + σ - γ σ) / (s (γ + h (-1 + γ) (s (-1 + σ) - σ)))} // Simplify
```
Out[ ]=
```
{0}
```

In[ ]:=
```
γ / (γ + (1 - γ) (s (1 - σ) + σ)) - {γ / (γ + s (-1 + γ) (-1 + σ) + σ - γ σ)} // Simplify
```
Out[ ]=
```
{0}
```

Here too we can write pA as $\frac{\mu}{(F+h-F h) s}$ but with F equal to the steady-state value:

In[ ]:=
```
μ (γ + (1 - γ) (s (1 - σ) + σ)) / (s (γ + h (1 - γ) (s (1 - σ) + σ))) -
  {μ / ((F + h - F h) s) /. F → γ / (γ + (1 - γ) (s (1 - σ) + σ))} // Simplify
```
Out[ ]=
```
{0}
```

#### Defining numerical functions, for single loci and multiple loci

Below we define functions for pA:

In[∘]:= $\text{pAnumSelf}[\mu\_, s\_, h\_, S\_, \gamma\_] := \frac{(2 - (1 - (2 - h)\, s)\, S\, (1 - \gamma))\, \mu}{s\, (S\, (1 - \gamma) + h\, (2 - (2 - s)\, S)\, (1 - \gamma) + 2\, \gamma)}$

$\text{pAnumAsex}[\mu\_, s\_, h\_, \sigma\_, \gamma\_] := \frac{\mu\, (\gamma + (1 - \gamma)\, (s\, (1 - \sigma) + \sigma))}{s\, (\gamma + h\, (1 - \gamma)\, (s\, (1 - \sigma) + \sigma))}$

Writing F as functions:

In[∘]:= $\text{FnumSelf}[s\_, S\_, \gamma\_, h\_] := \frac{(1 - h\, s)\, S\, (1 - \gamma) + 2\, \gamma}{2 - (1 - (2 - h)\, s)\, S\, (1 - \gamma)}$

$\text{FnumAsex}[s\_, \sigma\_, \gamma\_] := \frac{\gamma}{\gamma + (1 - \gamma)\, (s\, (1 - \sigma) + \sigma)}$

#### Contour plots for manuscript

```
In[∘]:= SetDirectory[NotebookDirectory[]];
```

```
In[•]:= pAAsex = Labeled[Labeled[ContourPlot[{ pAnumAsex[10^-8, 0.02, 0.01, σ, γ] / pAnumAsex[10^-8, 0.02, 0.01, 1, 0],
     pAnumAsex[10^-8, 0.02, 0.2, σ, γ] / pAnumAsex[10^-8, 0.02, 0.2, 1, 0], pAnumAsex[10^-8, 0.02, 0.5, σ, γ] / pAnumAsex[10^-8, 0.02, 0.5, 1, 0]},
    {σ, 0, 1}, {γ, 0, 0.1}, PlotLayout → {"Column", 3}, ContourStyle → Black,
    PlotLegends → BarLegend[{Automatic, {0.0, 1.0}}],
    FrameTicks → {{{0.00, 0.02, 0.04, 0.06, 0.08, 0.10}, None},
      {{0, 0.2, 0.4, 0.6, 0.8, 1.0}, None}}, ImagePadding → {{30, 5}, {30, 5}},
    LabelStyle → Directive[FontFamily → "Arial", FontSize → 14, FontColor → Black],
    Method → {Spacings → {30, 0}}, Mesh → None, PlotRange → All],
   {Text@TraditionalForm@Style["Frequency of Sex, σ", 16],
    Text@TraditionalForm@Style["Mitotic Gene Conversion, γ", 16], Text@
     TraditionalForm@Style["(a) h = 0.01                                        (b) h =
        0.2                                        (c) h = 0.5             ", 16]},
   {Bottom, Left, Top}, RotateLabel → True], Text@TraditionalForm@
   Style["Frequency of deleterious mutation p̂_A, scaled by μ/hs \n", 16], Top]
```

Out[•]=

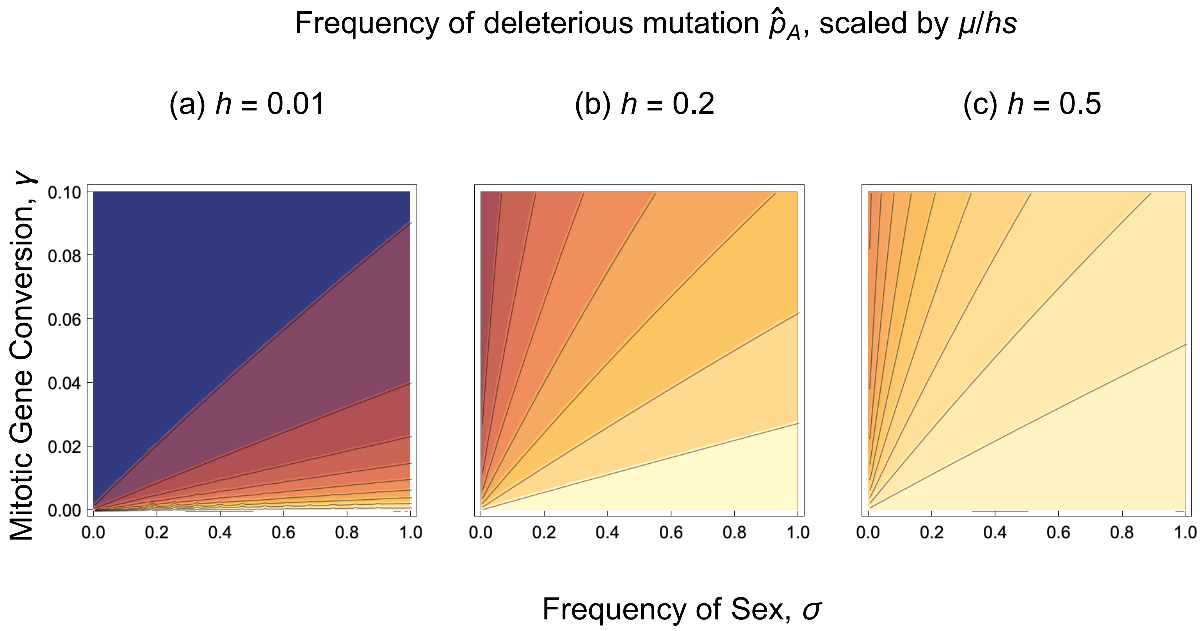

In[∘]:=
```
pASelf =
 Labeled[ContourPlot[{pAnumSelf[10^-8, 0.02, 0.01, S, γ] / pAnumSelf[10^-8, 0.02, 0.01, 0, 0],
   pAnumSelf[10^-8, 0.02, 0.2, S, γ] / pAnumSelf[10^-8, 0.02, 0.2, 0, 0], pAnumSelf[10^-8, 0.02, 0.5, S, γ] / pAnumSelf[10^-8, 0.02, 0.5, 0, 0]},
   {S, 0, 1}, {γ, 0, 0.1}, PlotLayout → {"Column", 3}, ContourStyle → Black,
   PlotLegends → BarLegend[{Automatic, {0.0, 1.0}}],
   FrameTicks → {{{0.00, 0.02, 0.04, 0.06, 0.08, 0.10}, None},
     {{0, 0.2, 0.4, 0.6, 0.8, 1.0}, None}}, ImagePadding → {{30, 5}, {30, 5}},
   LabelStyle → Directive[FontFamily → "Arial", FontSize → 14, FontColor → Black],
   Method → {Spacings → {30, 0}}, Mesh → None, PlotRange → All],
  {Text@TraditionalForm@Style["Frequency of Selfing, S", 16],
   Text@TraditionalForm@Style["Mitotic Gene Conversion, γ", 16],
   Text@TraditionalForm@Style["(a)  h  = 0.01                                    (b)  h
      = 0.2                                   (c)  h  = 0.5            ",
     16]}, {Bottom, Left, Top}, RotateLabel → True]
```

Out[∘]=

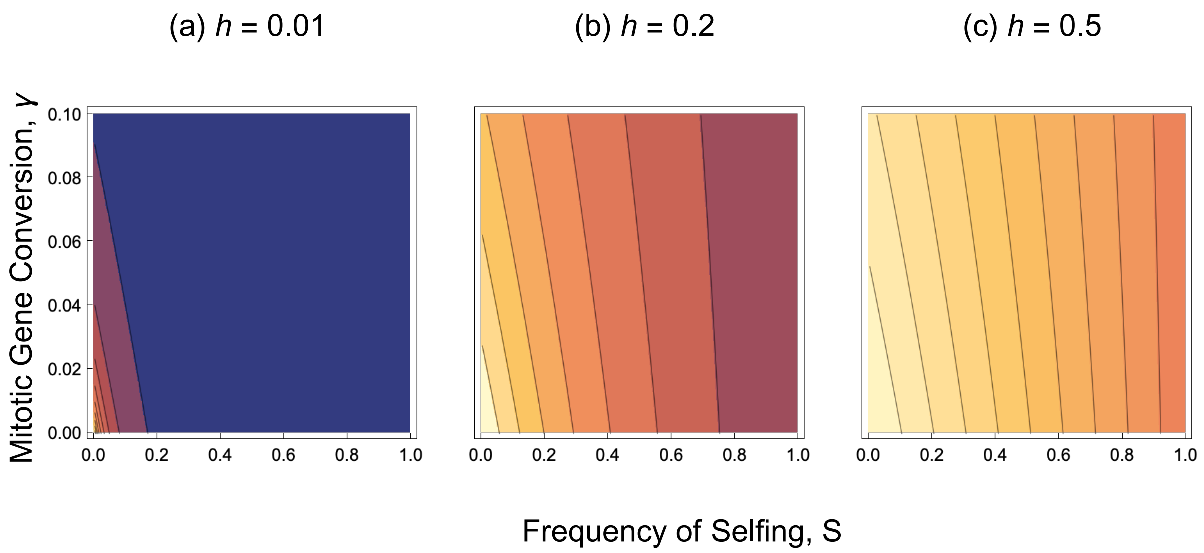

In[ ]:=
```
pFAsex = Labeled[Labeled[ContourPlot[{FnumAsex[0.02, σ, γ], FnumAsex[0.2, σ, γ]},
    {σ, 0, 1}, {γ, 0, 0.1}, ContourStyle → Black, PlotLayout → {"Column", 2},
    PlotLegends → BarLegend[{Automatic, {0, 1.0}}],
    FrameTicks → {{{0.00, 0.02, 0.04, 0.06, 0.08, 0.10}, None},
      {{0, 0.2, 0.4, 0.6, 0.8, 1.0}, None}}, ImagePadding → {{30, 5}, {30, 5}},
    LabelStyle → Directive[FontFamily → "Arial", FontSize → 14, FontColor → Black],
    Method → {Spacings → {30, 0}}, PlotRange → All],
   {Text@TraditionalForm@Style["Frequency of Sex, σ           ", 16],
    Text@TraditionalForm@Style["Mitotic Gene Conversion, γ", 16],
    Text@TraditionalForm@
      Style["(a) s = 0.02                                        (b) s =
         0.2      ", 16]}, {Bottom, Left, Top}, RotateLabel → True],
  Text@TraditionalForm@Style["Reduction in Heterozygosity F̂", 16],
  Top]
```

Out[ ]=

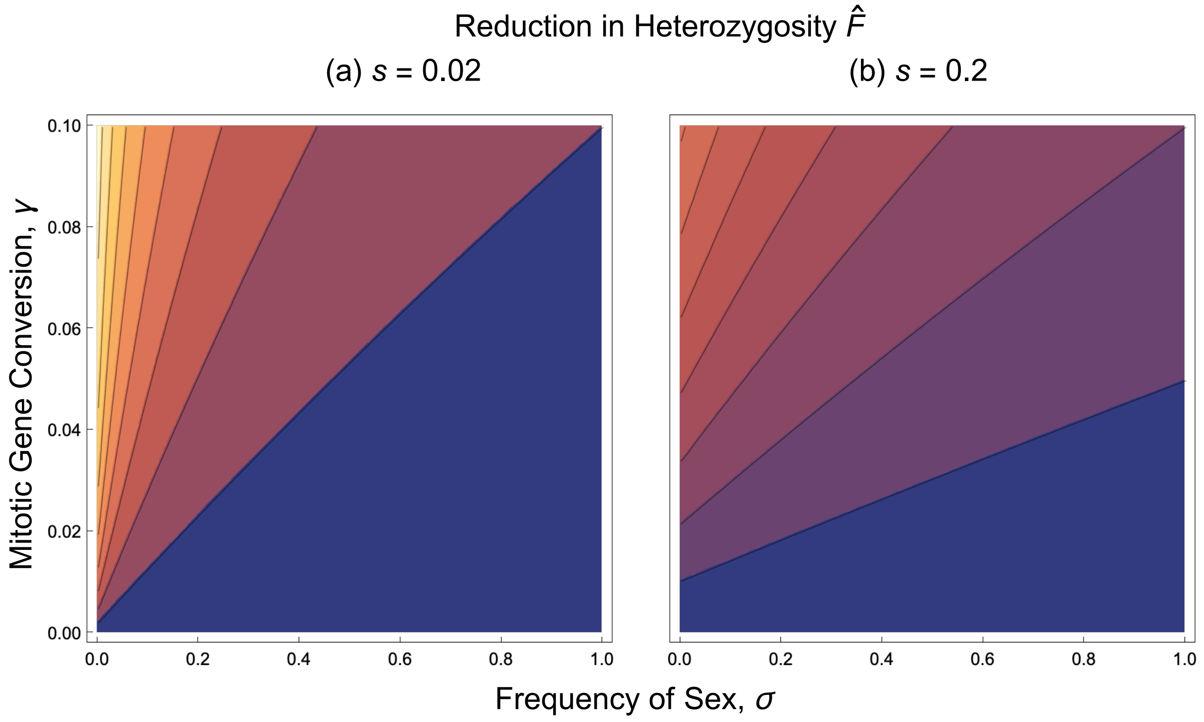

In[∘]:=
```
pFSelf = Labeled[ContourPlot[{FnumSelf[0.02, S, γ, 0.5], FnumSelf[0.2, S, γ, 0.5]},
    {S, 0, 1}, {γ, 0, 0.1}, ContourStyle → Black, PlotLayout → {"Column", 2},
    PlotLegends → BarLegend[{Automatic, {0, 1.0}}],
    FrameTicks → {{{0.00, 0.02, 0.04, 0.06, 0.08, 0.10}, None},
      {{0, 0.2, 0.4, 0.6, 0.8, 1.0}, None}}, ImagePadding → {{30, 5}, {30, 5}},
    LabelStyle → Directive[FontFamily → "Arial", FontSize → 14, FontColor → Black],
    Method → {Spacings → {30, 0}}, PlotRange → All],
   {Text@TraditionalForm@Style["Frequency of Selfing, S           ", 16],
    Text@TraditionalForm@Style["Mitotic Gene Conversion, γ", 16],
    Text@TraditionalForm@
      Style["(c) s = 0.02                                     (d) s
        = 0.2      ", 16]}, {Bottom, Left, Top}, RotateLabel → True]
```

Out[∘]=

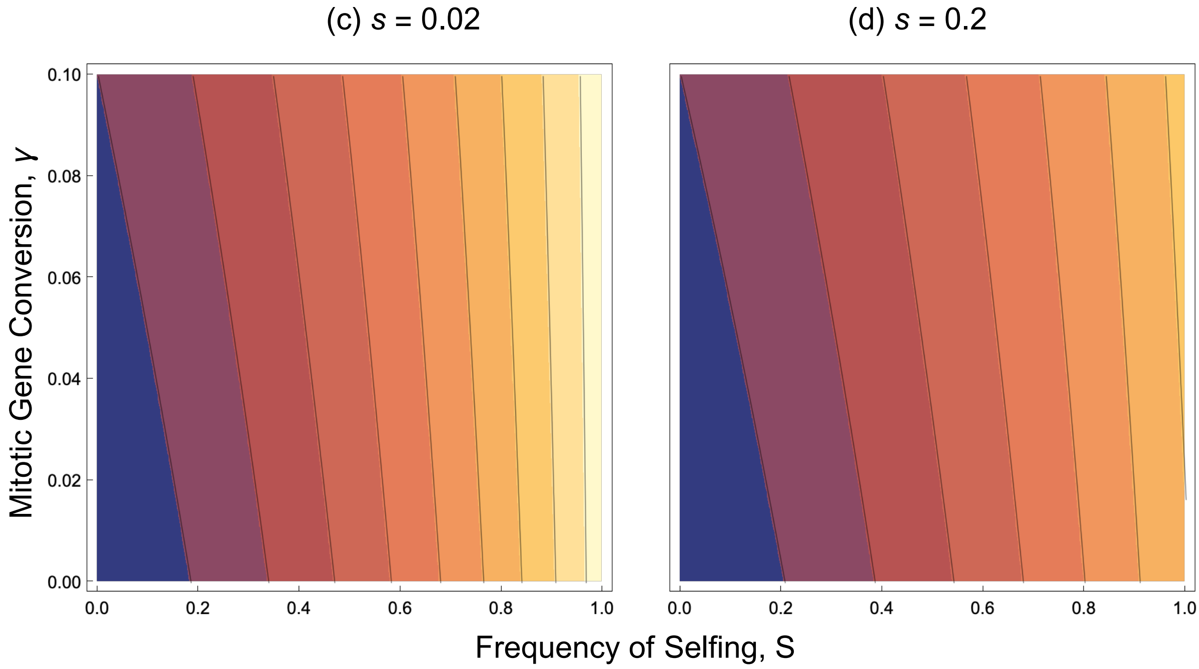


```
(* Extra command to remove tiling effects when exporting as PDF;
see https://mathematica.stackexchange.com/questions/2629/avoiding-
  white-lines-inside-filled-area-in-regionplot-exported-as-pdf-or-ps *)
```

In[∘]:=
```
pAAsex = pAAsex /. {EdgeForm[], r_?ColorQ, i___} :> {EdgeForm[r], r, i};
Export["Figures_R2/ContourPlots/pAAsex_Plot.pdf", pAAsex];
Export["Figures_R2/ContourPlots/pAAsex_Plot.jpg", pAAsex];
```

In[∘]:=
```
pASelf = pASelf /. {EdgeForm[], r_?ColorQ, i___} :> {EdgeForm[r], r, i};
Export["Figures_R2/ContourPlots/pASelf_Plot.pdf", pASelf];
Export["Figures_R2/ContourPlots/pASelf_Plot.jpg", pASelf];

pFAsex = pFAsex /. {EdgeForm[], r_?ColorQ, i___} :> {EdgeForm[r], r, i};
Export["Figures_R2/ContourPlots/FISAsex_plot.pdf", pFAsex];
Export["Figures_R2/ContourPlots/FISAsex_plot.jpg", pFAsex];

pFSelf = pFSelf /. {EdgeForm[], r_?ColorQ, i___} :> {EdgeForm[r], r, i};
Export["Figures_R2/ContourPlots/FISSelf_plot.pdf", pFSelf];
Export["Figures_R2/ContourPlots/FISSelf_plot.jpg", pFSelf];
```

### Numerical recursions, facultative sex

Below is the function for a change in genotype frequencies over a single generation:

```
In[∘]:= RecurGCA[{gaa_, gAa_, gAA_}] :=
  {-(((-1 + μ) (4 gaa^2 (1 + (-1 + γ) μ) + gaa (-2 gAa (-1 + h s) (2 + γ - 2 μ + 2 γ μ) +
           4 gAA (-1 + s) (-1 + (-1 + γ) μ (-1 + σ) + σ - γ σ)) +
        gAa (-1 + h s) (2 gAA (-1 + s) γ + gAa (-1 + h s) (σ - μ σ + γ (2 + (-1 + μ) σ))))) /
     (4 (gaa + gAa + gAA - gAA s - gAa h s)^2)), ((-1 + γ) (-1 + μ)
     (4 gaa^2 μ + gaa (-2 gAa (-1 + h s) (1 + 2 μ) + 4 gAA (-1 + s) (μ (-1 + σ) - σ)) +
       gAa (-1 + h s) (2 gAA (-1 + s) + gAa (-1 + h s) (2 + (-1 + μ) σ)))) /
   (2 (gaa + gAa + gAA - gAA s - gAa h s)^2), (4 gAA^2 (-1 + s)^2 + 4 gaa^2 μ (γ + μ - γ μ) -
     2 gAa gAA (-1 + s) (-1 + h s) (γ (-1 + μ) - 2 (1 + μ)) +
     2 gaa gAa (-1 + h s) (-2 μ (1 + μ) + γ (-1 - μ + 2 μ^2)) -
     4 gaa gAA (-1 + s) (1 + γ (-1 + μ) (μ (-1 + σ) - σ) - μ^2 (-1 + σ) - σ + 2 μ σ) -
     gAa^2 (-1 + h s)^2 (2 μ (-2 + σ) - σ - μ^2 σ + γ (-1 + μ) (2 + (-1 + μ) σ))) /
   (4 (gaa + gAa + gAA - gAA s - gAa h s)^2)};
```

Defining range of $\gamma$, $\sigma$ to be evaluated over:

```
In[∘]:= γit = Table[0 + 0.1/10 j, {j, 0, 10}];
σit = {1, 0.5, 0.01};
```

#### h = 0.01, s = 0.02 plots

```
In[∘]:= Clear[freqAa, μ, s, h, RecTime]
freqAa = 0.001;
μ = 10^-8;
s = 0.02;
h = 0.01;
RecTime = 50 000;
```

```
In[•]:= Clear[Vrin];
σ = σit[[1]];
Vrin = Table[Clear[RecurRes, γ];
   γ = γit[[j + 1]];
   RecurRes = NestList[RecurGCA, {1 - freqAa, freqAa, 0}, RecTime] // N;
   {γ, RecurRes[[RecTime]][[3]] + RecurRes[[RecTime]][[2]] / 2,
    (RecurRes[[RecTime]][[3]] * RecurRes[[RecTime]][[1]] - (RecurRes[[RecTime]][[2]] / 2)^2) /
     ((RecurRes[[RecTime]][[3]] + RecurRes[[RecTime]][[2]] / 2)
       (RecurRes[[RecTime]][[1]] + RecurRes[[RecTime]][[2]] / 2))}, {j, 0, 10}];
pArecσ1 = Partition[Drop[Flatten[Vrin], {3, Length[Flatten[Vrin]], 3}], 2];
Frecσ1 = Partition[Drop[Flatten[Vrin], {2, Length[Flatten[Vrin]] - 1, 3}], 2];

Clear[Vrin];
σ = σit[[2]];
Vrin = Table[Clear[RecurRes, γ];
   γ = γit[[j + 1]];
   RecurRes = NestList[RecurGCA, {1 - freqAa, freqAa, 0}, RecTime] // N;
   {γ, RecurRes[[RecTime]][[3]] + RecurRes[[RecTime]][[2]] / 2,
    (RecurRes[[RecTime]][[3]] * RecurRes[[RecTime]][[1]] - (RecurRes[[RecTime]][[2]] / 2)^2) /
     ((RecurRes[[RecTime]][[3]] + RecurRes[[RecTime]][[2]] / 2)
       (RecurRes[[RecTime]][[1]] + RecurRes[[RecTime]][[2]] / 2))}, {j, 0, 10}];
pArecσ05 = Partition[Drop[Flatten[Vrin], {3, Length[Flatten[Vrin]], 3}], 2];
Frecσ05 = Partition[Drop[Flatten[Vrin], {2, Length[Flatten[Vrin]] - 1, 3}], 2];

Clear[Vrin];
σ = σit[[3]];
Vrin = Table[Clear[RecurRes, γ];
   γ = γit[[j + 1]];
   RecurRes = NestList[RecurGCA, {1 - freqAa, freqAa, 0}, RecTime] // N;
   {γ, RecurRes[[RecTime]][[3]] + RecurRes[[RecTime]][[2]] / 2,
    (RecurRes[[RecTime]][[3]] * RecurRes[[RecTime]][[1]] - (RecurRes[[RecTime]][[2]] / 2)^2) /
     ((RecurRes[[RecTime]][[3]] + RecurRes[[RecTime]][[2]] / 2)
       (RecurRes[[RecTime]][[1]] + RecurRes[[RecTime]][[2]] / 2))}, {j, 0, 10}];
pArecσ001 = Partition[Drop[Flatten[Vrin], {3, Length[Flatten[Vrin]], 3}], 2];
Frecσ001 = Partition[Drop[Flatten[Vrin], {2, Length[Flatten[Vrin]] - 1, 3}], 2];
```

In[ ]:=
```
p1σ1 = ListPlot[pArecσ1, PlotStyle → {Red, PointSize[Large]}, PlotLegends →
    Placed[PointLegend[{Directive[Red, AbsolutePointSize[8]], Directive[Orange,
        AbsolutePointSize[8]], Directive[Blue, AbsolutePointSize[8]]},
      {"σ = 1", "σ = 0.5", "σ = 0.01"}], {0.4, 0.7}], LabelStyle → 18];
p1σ05 = ListPlot[pArecσ05, PlotStyle → {Orange, PointSize[Large]},
   PlotLegends → Placed[PointLegend[{Directive[Black, AbsolutePointSize[8]]},
      {"Numerical Recursions"}], {0.8, 0.8}], LabelStyle → 18];
p1σ001 = ListPlot[pArecσ001, PlotStyle → {Blue, PointSize[Large]}];

p2σ1 = Plot[{pAnumAsex[μ, s, h, 1, γin]},
   {γin, 0, 0.1}, PlotRange → Full, PlotStyle → {{Red}}, PlotLegends →
    Placed[LineLegend[{Black}, {"Equation 1"}, LabelStyle → 18], {0.75, 0.65}]];
p2σ05 = Plot[{pAnumAsex[μ, s, h, 0.5, γin]},
   {γin, 0, 0.1}, PlotRange → Full, PlotStyle → {{Orange}}];
p2σ001 = Plot[{pAnumAsex[μ, s, h, 0.01, γin]},
   {γin, 0, 0.1}, PlotRange → Full, PlotStyle → {{Blue}}];
```

In[ ]:=
```
plotpA1 = Labeled[Show[p1σ1, p2σ1, p1σ001, p2σ001, p1σ05, p2σ05, ImageSize → 550,
   LabelStyle → {FontFamily → "Arial", FontSize → 18, FontColor → Black}],
  {Text@TraditionalForm@Style["Frequency\nOf Deleterious\nAllele", 18], Text@
    TraditionalForm@Style["Mitotic Gene Conversion, γ", 18]}, {Left, Bottom}]
```

Out[ ]=

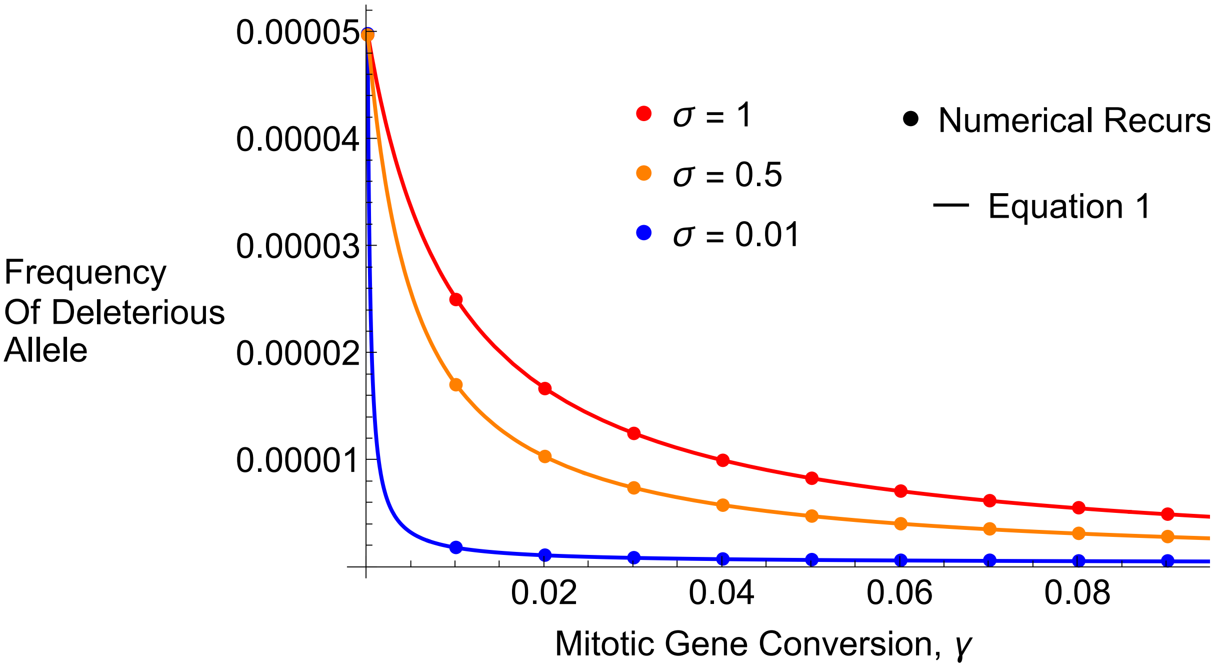


In[ ]:=
```
Export["Figures_Jan2025/RecursionsAsex/RecPlotpA_h001_s002.pdf", plotpA1];
Export["Figures_Jan2025/RecursionsAsex/RecPlotpA_h001_s002.jpg", plotpA1];
```

```
In[ ]:= p1Fσ1 = ListPlot[Frecσ1, PlotStyle → {Red, PointSize[Large]}];
        p1Fσ05 = ListPlot[Frecσ05, PlotStyle → {Orange, PointSize[Large]}];
        p1Fσ001 = ListPlot[Frecσ001, PlotStyle → {Blue, PointSize[Large]}];

        p2Fσ1 = Plot[{FnumAsex[s, 1, γin]}, {γin, 0, 0.1},
           PlotRange → Full, PlotStyle → {{Red}}, PlotLegends →
            Placed[LineLegend[{Black}, {"Equation 2"}, LabelStyle → 18], {0.75, 0.65}]];
        p2Fσ05 = Plot[{FnumAsex[s, 0.5, γin]},
           {γin, 0, 0.1}, PlotRange → Full, PlotStyle → {{Orange}}];
        p2Fσ001 = Plot[{FnumAsex[s, 0.01, γin]},
           {γin, 0, 0.1}, PlotRange → Full, PlotStyle → {{Blue}}];
```

```
In[ ]:= plotF1 =
         Labeled[Show[p1Fσ001, p2Fσ001, p1Fσ1, p2Fσ1, p1Fσ05, p2Fσ05, ImageSize → 550,
           LabelStyle → {FontFamily → "Arial", FontSize → 18, FontColor → Black}],
          {Text@TraditionalForm@Style["Inbreeding\nCoefficient", 18], Text@
            TraditionalForm@Style["Mitotic Gene Conversion, γ", 18]}, {Left, Bottom}]
```

Out[ ]=

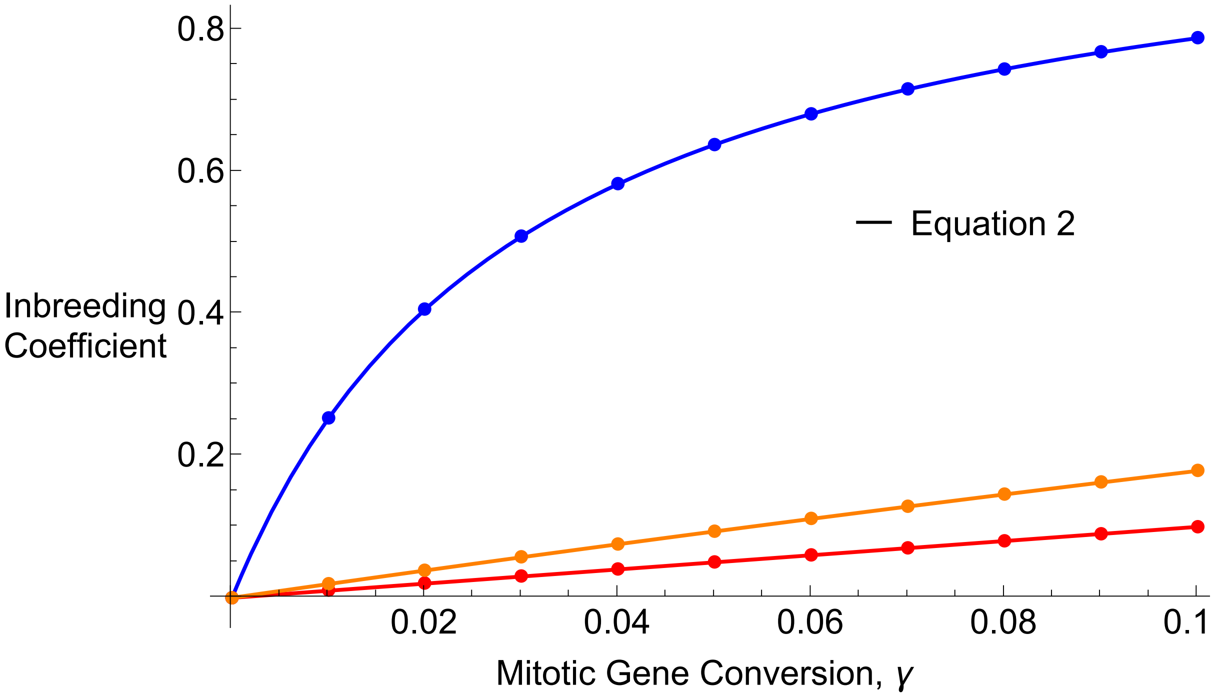


```
In[ ]:= Export["Figures_Jan2025/RecursionsAsex/RecPlotF_h001_s002.pdf", plotF1];
        Export["Figures_Jan2025/RecursionsAsex/RecPlotF_h001_s002.jpg", plotF1];
```

##### h = 0.5, s = 0.02 plots

```
In[ ]:= Clear[freqAa, μ, s, h, RecTime]
        freqAa = 0.001;
        μ = 10^-8;
        s = 0.02;
        h = 0.5;
        RecTime = 50 000;
```

```
In[•]:= Clear[Vrin];
σ = σit[[1]];
Vrin = Table[Clear[RecurRes, γ];
   γ = γit[[j + 1]];
   RecurRes = NestList[RecurGCA, {1 - freqAa, freqAa, 0}, RecTime] // N;
   {γ, RecurRes[[RecTime]][[3]] + RecurRes[[RecTime]][[2]] / 2,
    (RecurRes[[RecTime]][[3]] * RecurRes[[RecTime]][[1]] - (RecurRes[[RecTime]][[2]] / 2)^2) /
     ((RecurRes[[RecTime]][[3]] + RecurRes[[RecTime]][[2]] / 2)
       (RecurRes[[RecTime]][[1]] + RecurRes[[RecTime]][[2]] / 2))}, {j, 0, 10}];
pA2recσ1 = Partition[Drop[Flatten[Vrin], {3, Length[Flatten[Vrin]], 3}], 2];
F2recσ1 = Partition[Drop[Flatten[Vrin], {2, Length[Flatten[Vrin]] - 1, 3}], 2];

Clear[Vrin];
σ = σit[[2]];
Vrin = Table[Clear[RecurRes, γ];
   γ = γit[[j + 1]];
   RecurRes = NestList[RecurGCA, {1 - freqAa, freqAa, 0}, RecTime] // N;
   {γ, RecurRes[[RecTime]][[3]] + RecurRes[[RecTime]][[2]] / 2,
    (RecurRes[[RecTime]][[3]] * RecurRes[[RecTime]][[1]] - (RecurRes[[RecTime]][[2]] / 2)^2) /
     ((RecurRes[[RecTime]][[3]] + RecurRes[[RecTime]][[2]] / 2)
       (RecurRes[[RecTime]][[1]] + RecurRes[[RecTime]][[2]] / 2))}, {j, 0, 10}];
pA2recσ05 = Partition[Drop[Flatten[Vrin], {3, Length[Flatten[Vrin]], 3}], 2];
F2recσ05 = Partition[Drop[Flatten[Vrin], {2, Length[Flatten[Vrin]] - 1, 3}], 2];

Clear[Vrin];
σ = σit[[3]];
Vrin = Table[Clear[RecurRes, γ];
   γ = γit[[j + 1]];
   RecurRes = NestList[RecurGCA, {1 - freqAa, freqAa, 0}, RecTime] // N;
   {γ, RecurRes[[RecTime]][[3]] + RecurRes[[RecTime]][[2]] / 2,
    (RecurRes[[RecTime]][[3]] * RecurRes[[RecTime]][[1]] - (RecurRes[[RecTime]][[2]] / 2)^2) /
     ((RecurRes[[RecTime]][[3]] + RecurRes[[RecTime]][[2]] / 2)
       (RecurRes[[RecTime]][[1]] + RecurRes[[RecTime]][[2]] / 2))}, {j, 0, 10}];
pA2recσ001 = Partition[Drop[Flatten[Vrin], {3, Length[Flatten[Vrin]], 3}], 2];
F2recσ001 = Partition[Drop[Flatten[Vrin], {2, Length[Flatten[Vrin]] - 1, 3}], 2];

In[•]:= p3σ1 = ListPlot[pA2recσ1, PlotStyle → {Red, PointSize[Large]}];
p3σ05 = ListPlot[pA2recσ05, PlotStyle → {Orange, PointSize[Large]}];
p3σ001 = ListPlot[pA2recσ001, PlotStyle → {Blue, PointSize[Large]}];

p4σ1 = Plot[{pAnumAsex[μ, s, h, 1, γin]},
   {γin, 0, 0.1}, PlotRange → Full, PlotStyle → {{Red}}];
p4σ05 = Plot[{pAnumAsex[μ, s, h, 0.5, γin]},
   {γin, 0, 0.1}, PlotRange → Full, PlotStyle → {{Orange}}];
p4σ001 = Plot[{pAnumAsex[μ, s, h, 0.01, γin]},
   {γin, 0, 0.1}, PlotRange → Full, PlotStyle → {{Blue}}];
```

In[ ]:=
```
plotpA2 = Labeled[Show[p4σ001, p3σ001, p3σ05, p4σ05, p3σ1, p4σ1, ImageSize → 550,
    LabelStyle → {FontFamily → "Arial", FontSize → 18, FontColor → Black}],
  {Text@TraditionalForm@Style["Frequency\nOf Deleterious\nAllele", 18], Text@
    TraditionalForm@Style["Mitotic Gene Conversion, γ", 18]}, {Left, Bottom}]
```

Out[ ]=

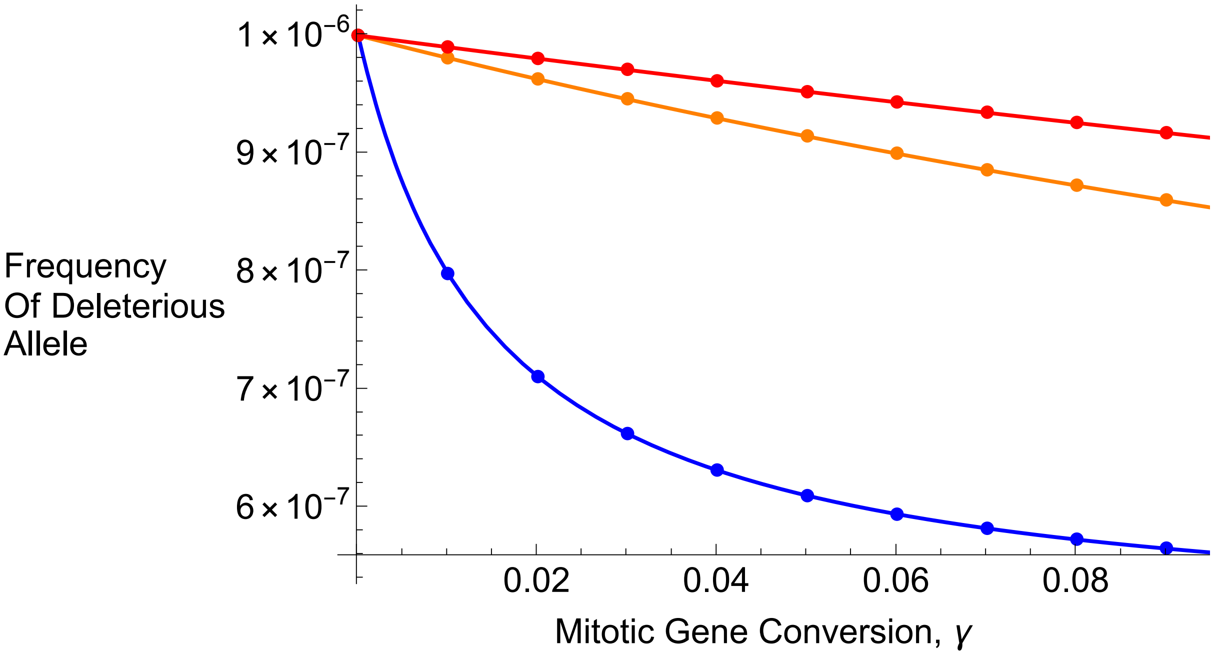


In[ ]:=
```
p3Fσ1 = ListPlot[F2recσ1, PlotStyle → {Red, PointSize[Large]}];
p3Fσ05 = ListPlot[F2recσ05, PlotStyle → {Orange, PointSize[Large]}];
p3Fσ001 = ListPlot[F2recσ001, PlotStyle → {Blue, PointSize[Large]}];

p4Fσ1 = Plot[{FnumAsex[s, 1, γin]},
   {γin, 0, 0.1}, PlotRange → Full, PlotStyle → {{Red}}];
p4Fσ05 = Plot[{FnumAsex[s, 0.5, γin]},
   {γin, 0, 0.1}, PlotRange → Full, PlotStyle → {{Orange}}];
p4Fσ001 = Plot[{FnumAsex[s, 0.01, γin]},
   {γin, 0, 0.1}, PlotRange → Full, PlotStyle → {{Blue}}];
```

In[ ]:=
```
plotF2 =
  Labeled[Show[p4Fσ001, p3Fσ001, p3Fσ1, p4Fσ1, p3Fσ05, p4Fσ05, ImageSize → 550,
    LabelStyle → {FontFamily → "Arial", FontSize → 18, FontColor → Black}],
   {Text@TraditionalForm@Style["Inbreeding\nCoefficient", 18], Text@
     TraditionalForm@Style["Mitotic Gene Conversion, γ", 18]}, {Left, Bottom}]
```

Out[ ]=

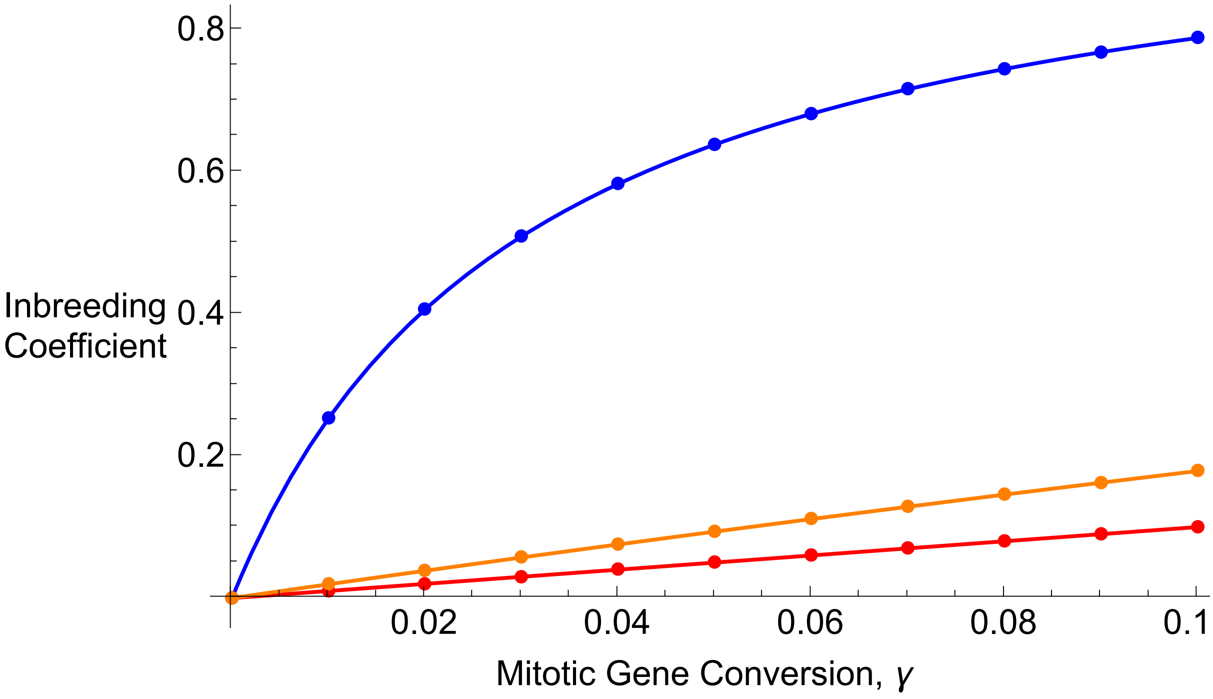


In[ ]:=
```
Export["Figures_Jan2025/RecursionsAsex/RecPlotpA_h05_s002.pdf", plotpA2];
Export["Figures_Jan2025/RecursionsAsex/RecPlotpA_h05_s002.jpg", plotpA2];

Export["Figures_Jan2025/RecursionsAsex/RecPlotF_h05_s002.pdf", plotF2];
Export["Figures_Jan2025/RecursionsAsex/RecPlotF_h05_s002.jpg", plotF2];
```

#### h = 0.01, s = 0.2 plots

In[ ]:=
```
Clear[freqAa, μ, s, h, RecTime]
freqAa = 0.001;
μ = 10^-8;
s = 0.2;
h = 0.01;
RecTime = 50 000;
```

```
In[ ]:= Clear[Vrin];
σ = σit[[1]];
Vrin = Table[Clear[RecurRes, γ];
   γ = γit[[j + 1]];
   RecurRes = NestList[RecurGCA, {1 - freqAa, freqAa, 0}, RecTime] // N;
   {γ, RecurRes[[RecTime]][[3]] + RecurRes[[RecTime]][[2]] / 2,
    (RecurRes[[RecTime]][[3]] * RecurRes[[RecTime]][[1]] - (RecurRes[[RecTime]][[2]] / 2)^2) /
     ((RecurRes[[RecTime]][[3]] + RecurRes[[RecTime]][[2]] / 2)
       (RecurRes[[RecTime]][[1]] + RecurRes[[RecTime]][[2]] / 2))}, {j, 0, 10}];
pA3recσ1 = Partition[Drop[Flatten[Vrin], {3, Length[Flatten[Vrin]], 3}], 2];
F3recσ1 = Partition[Drop[Flatten[Vrin], {2, Length[Flatten[Vrin]] - 1, 3}], 2];

Clear[Vrin];
σ = σit[[2]];
Vrin = Table[Clear[RecurRes, γ];
   γ = γit[[j + 1]];
   RecurRes = NestList[RecurGCA, {1 - freqAa, freqAa, 0}, RecTime] // N;
   {γ, RecurRes[[RecTime]][[3]] + RecurRes[[RecTime]][[2]] / 2,
    (RecurRes[[RecTime]][[3]] * RecurRes[[RecTime]][[1]] - (RecurRes[[RecTime]][[2]] / 2)^2) /
     ((RecurRes[[RecTime]][[3]] + RecurRes[[RecTime]][[2]] / 2)
       (RecurRes[[RecTime]][[1]] + RecurRes[[RecTime]][[2]] / 2))}, {j, 0, 10}];
pA3recσ05 = Partition[Drop[Flatten[Vrin], {3, Length[Flatten[Vrin]], 3}], 2];
F3recσ05 = Partition[Drop[Flatten[Vrin], {2, Length[Flatten[Vrin]] - 1, 3}], 2];

Clear[Vrin];
σ = σit[[3]];
Vrin = Table[Clear[RecurRes, γ];
   γ = γit[[j + 1]];
   RecurRes = NestList[RecurGCA, {1 - freqAa, freqAa, 0}, RecTime] // N;
   {γ, RecurRes[[RecTime]][[3]] + RecurRes[[RecTime]][[2]] / 2,
    (RecurRes[[RecTime]][[3]] * RecurRes[[RecTime]][[1]] - (RecurRes[[RecTime]][[2]] / 2)^2) /
     ((RecurRes[[RecTime]][[3]] + RecurRes[[RecTime]][[2]] / 2)
       (RecurRes[[RecTime]][[1]] + RecurRes[[RecTime]][[2]] / 2))}, {j, 0, 10}];
pA3recσ001 = Partition[Drop[Flatten[Vrin], {3, Length[Flatten[Vrin]], 3}], 2];
F3recσ001 = Partition[Drop[Flatten[Vrin], {2, Length[Flatten[Vrin]] - 1, 3}], 2];

In[ ]:= p5σ1 = ListPlot[pA3recσ1, PlotStyle → {Red, PointSize[Large]}];
p5σ05 = ListPlot[pA3recσ05, PlotStyle → {Orange, PointSize[Large]}];
p5σ001 = ListPlot[pA3recσ001, PlotStyle → {Blue, PointSize[Large]}];

p6σ1 = Plot[{pAnumAsex[μ, s, h, 1, γin]},
   {γin, 0, 0.1}, PlotRange → Full, PlotStyle → {{Red}}];
p6σ05 = Plot[{pAnumAsex[μ, s, h, 0.5, γin]},
   {γin, 0, 0.1}, PlotRange → Full, PlotStyle → {{Orange}}];
p6σ001 = Plot[{pAnumAsex[μ, s, h, 0.01, γin]},
   {γin, 0, 0.1}, PlotRange → Full, PlotStyle → {{Blue}}];
```

```
In[◦]:= plotpA3 = Labeled[Show[p6σ001, p5σ001, p5σ05, p6σ05, p5σ1, p6σ1, ImageSize → 550,
           LabelStyle → {FontFamily → "Arial", FontSize → 18, FontColor → Black}],
          {Text@TraditionalForm@Style["Frequency\nOf Deleterious\nAllele", 18], Text@
            TraditionalForm@Style["Mitotic Gene Conversion, γ", 18]}, {Left, Bottom}]
Out[◦]=
```

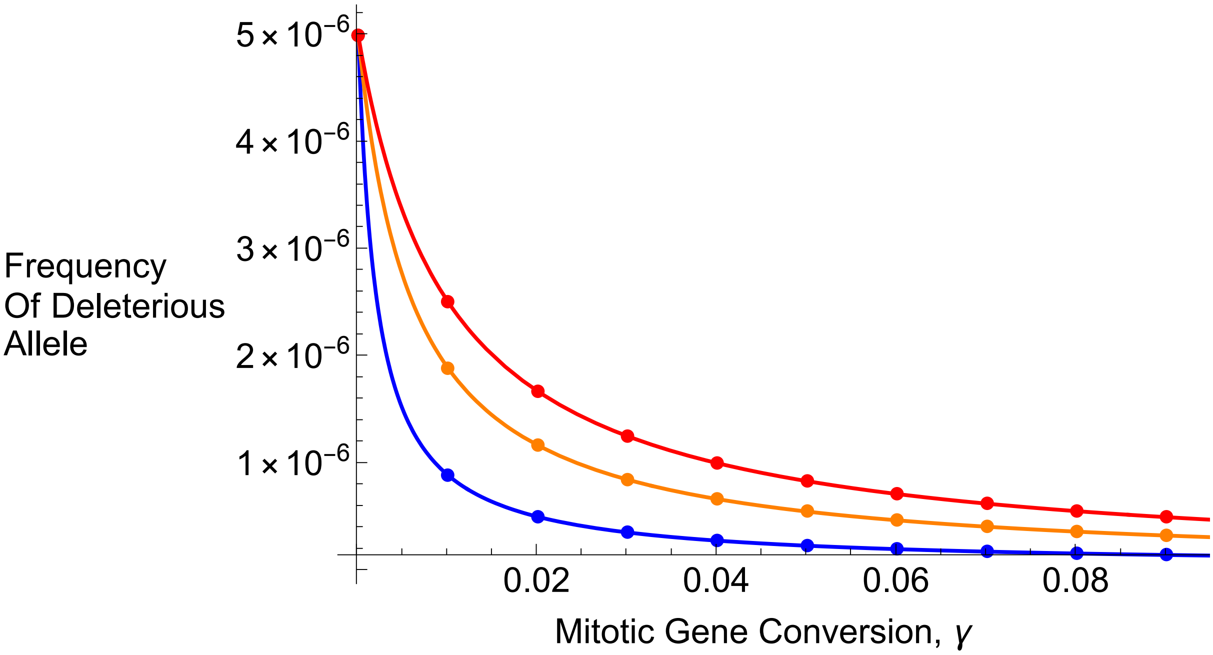


```
In[◦]:= p5Fσ1 = ListPlot[F3recσ1, PlotStyle → {Red, PointSize[Large]}];
        p5Fσ05 = ListPlot[F3recσ05, PlotStyle → {Orange, PointSize[Large]}];
        p5Fσ001 = ListPlot[F3recσ001, PlotStyle → {Blue, PointSize[Large]}];

        p6Fσ1 = Plot[{FnumAsex[s, 1, γin]},
            {γin, 0, 0.1}, PlotRange → Full, PlotStyle → {{Red}}];
        p6Fσ05 = Plot[{FnumAsex[s, 0.5, γin]},
            {γin, 0, 0.1}, PlotRange → Full, PlotStyle → {{Orange}}];
        p6Fσ001 = Plot[{FnumAsex[s, 0.01, γin]},
            {γin, 0, 0.1}, PlotRange → Full, PlotStyle → {{Blue}}];
```

In[ ]:=
```
plotF3 =
  Labeled[Show[p6Fσ001, p5Fσ001, p5Fσ1, p6Fσ1, p5Fσ05, p6Fσ05, ImageSize → 550,
    LabelStyle → {FontFamily → "Arial", FontSize → 18, FontColor → Black}],
   {Text@TraditionalForm@Style["Inbreeding\nCoefficient", 18], Text@
     TraditionalForm@Style["Mitotic Gene Conversion, γ", 18]}, {Left, Bottom}]
```

Out[ ]=

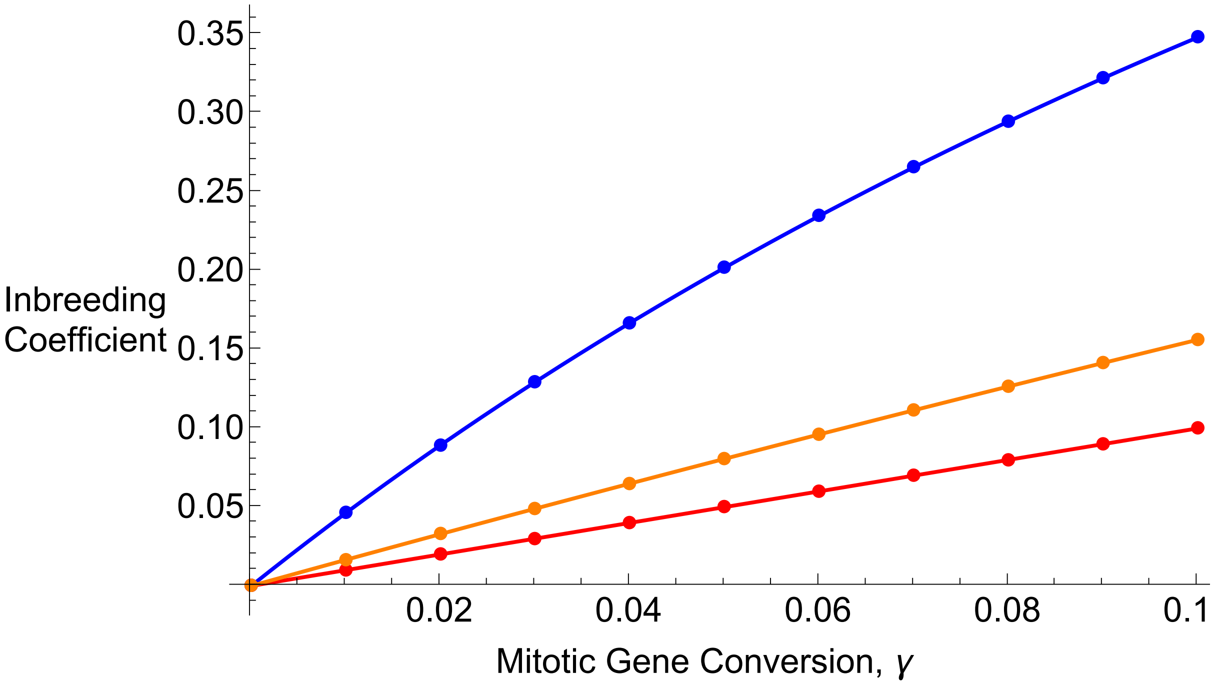


In[ ]:=
```
Export["Figures_Jan2025/RecursionsAsex/RecPlotpA_h001_s02.pdf", plotpA3];
Export["Figures_Jan2025/RecursionsAsex/RecPlotpA_h001_s02.jpg", plotpA3];

Export["Figures_Jan2025/RecursionsAsex/RecPlotF_h001_s02.pdf", plotF3];
Export["Figures_Jan2025/RecursionsAsex/RecPlotF_h001_s02.jpg", plotF3];
```

#### h = 0.5, s = 0.2 plots

In[ ]:=
```
Clear[freqAa, μ, s, h, RecTime]
freqAa = 0.001;
μ = 10^-8;
s = 0.2;
h = 0.5;
RecTime = 50 000;
```

```
In[•]:= Clear[Vrin];
σ = σit⟦1⟧;
Vrin = Table[Clear[RecurRes, γ];
   γ = γit⟦j + 1⟧;
   RecurRes = NestList[RecurGCA, {1 - freqAa, freqAa, 0}, RecTime] // N;
   {γ, RecurRes⟦RecTime⟧⟦3⟧ + RecurRes⟦RecTime⟧⟦2⟧ / 2,
    (RecurRes⟦RecTime⟧⟦3⟧ * RecurRes⟦RecTime⟧⟦1⟧ - (RecurRes⟦RecTime⟧⟦2⟧ / 2)^2) /
     ((RecurRes⟦RecTime⟧⟦3⟧ + RecurRes⟦RecTime⟧⟦2⟧ / 2)
       (RecurRes⟦RecTime⟧⟦1⟧ + RecurRes⟦RecTime⟧⟦2⟧ / 2))}, {j, 0, 10}];
pA4recσ1 = Partition[Drop[Flatten[Vrin], {3, Length[Flatten[Vrin]], 3}], 2];
F4recσ1 = Partition[Drop[Flatten[Vrin], {2, Length[Flatten[Vrin]] - 1, 3}], 2];

Clear[Vrin];
σ = σit⟦2⟧;
Vrin = Table[Clear[RecurRes, γ];
   γ = γit⟦j + 1⟧;
   RecurRes = NestList[RecurGCA, {1 - freqAa, freqAa, 0}, RecTime] // N;
   {γ, RecurRes⟦RecTime⟧⟦3⟧ + RecurRes⟦RecTime⟧⟦2⟧ / 2,
    (RecurRes⟦RecTime⟧⟦3⟧ * RecurRes⟦RecTime⟧⟦1⟧ - (RecurRes⟦RecTime⟧⟦2⟧ / 2)^2) /
     ((RecurRes⟦RecTime⟧⟦3⟧ + RecurRes⟦RecTime⟧⟦2⟧ / 2)
       (RecurRes⟦RecTime⟧⟦1⟧ + RecurRes⟦RecTime⟧⟦2⟧ / 2))}, {j, 0, 10}];
pA4recσ05 = Partition[Drop[Flatten[Vrin], {3, Length[Flatten[Vrin]], 3}], 2];
F4recσ05 = Partition[Drop[Flatten[Vrin], {2, Length[Flatten[Vrin]] - 1, 3}], 2];

Clear[Vrin];
σ = σit⟦3⟧;
Vrin = Table[Clear[RecurRes, γ];
   γ = γit⟦j + 1⟧;
   RecurRes = NestList[RecurGCA, {1 - freqAa, freqAa, 0}, RecTime] // N;
   {γ, RecurRes⟦RecTime⟧⟦3⟧ + RecurRes⟦RecTime⟧⟦2⟧ / 2,
    (RecurRes⟦RecTime⟧⟦3⟧ * RecurRes⟦RecTime⟧⟦1⟧ - (RecurRes⟦RecTime⟧⟦2⟧ / 2)^2) /
     ((RecurRes⟦RecTime⟧⟦3⟧ + RecurRes⟦RecTime⟧⟦2⟧ / 2)
       (RecurRes⟦RecTime⟧⟦1⟧ + RecurRes⟦RecTime⟧⟦2⟧ / 2))}, {j, 0, 10}];
pA4recσ001 = Partition[Drop[Flatten[Vrin], {3, Length[Flatten[Vrin]], 3}], 2];
F4recσ001 = Partition[Drop[Flatten[Vrin], {2, Length[Flatten[Vrin]] - 1, 3}], 2];
```

```
In[•]:= p7σ1 = ListPlot[pA4recσ1, PlotStyle → {Red, PointSize[Large]}];
p7σ05 = ListPlot[pA4recσ05, PlotStyle → {Orange, PointSize[Large]}];
p7σ001 = ListPlot[pA4recσ001, PlotStyle → {Blue, PointSize[Large]}];

p8σ1 = Plot[{pAnumAsex[μ, s, h, 1, γin]},
   {γin, 0, 0.1}, PlotRange → Full, PlotStyle → {{Red}}];
p8σ05 = Plot[{pAnumAsex[μ, s, h, 0.5, γin]},
   {γin, 0, 0.1}, PlotRange → Full, PlotStyle → {{Orange}}];
p8σ001 = Plot[{pAnumAsex[μ, s, h, 0.01, γin]},
   {γin, 0, 0.1}, PlotRange → Full, PlotStyle → {{Blue}}];
```

```
In[ ]:= plotpA4 = Labeled[Show[p8σ001, p7σ001, p7σ05, p8σ05, p7σ1, p8σ1, ImageSize → 550,
      LabelStyle → {FontFamily → "Arial", FontSize → 18, FontColor → Black}],
    {Text@TraditionalForm@Style["Frequency\nOf Deleterious\nAllele", 18], Text@
      TraditionalForm@Style["Mitotic Gene Conversion, γ", 18]}, {Left, Bottom}]
```

Out[ ]=

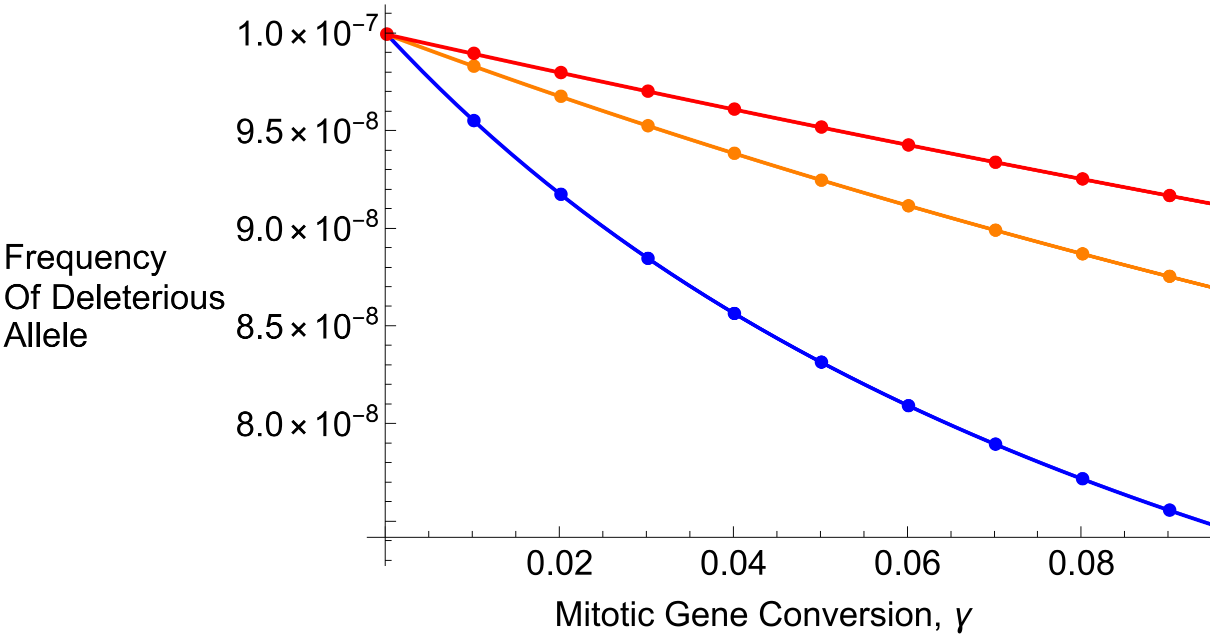


```
In[ ]:= p7Fσ1 = ListPlot[F4recσ1, PlotStyle → {Red, PointSize[Large]}];
p7Fσ05 = ListPlot[F4recσ05, PlotStyle → {Orange, PointSize[Large]}];
p7Fσ001 = ListPlot[F4recσ001, PlotStyle → {Blue, PointSize[Large]}];

p8Fσ1 = Plot[{FnumAsex[s, 1, γin]},
   {γin, 0, 0.1}, PlotRange → Full, PlotStyle → {{Red}}];
p8Fσ05 = Plot[{FnumAsex[s, 0.5, γin]},
   {γin, 0, 0.1}, PlotRange → Full, PlotStyle → {{Orange}}];
p8Fσ001 = Plot[{FnumAsex[s, 0.01, γin]},
   {γin, 0, 0.1}, PlotRange → Full, PlotStyle → {{Blue}}];
```

In[ ]:=
```
plotF4 =
  Labeled[Show[p7Fσ001, p8Fσ001, p7Fσ1, p8Fσ1, p7Fσ05, p8Fσ05, ImageSize → 550,
    LabelStyle → {FontFamily → "Arial", FontSize → 18, FontColor → Black}],
   {Text@TraditionalForm@Style["Inbreeding\nCoefficient", 18], Text@
     TraditionalForm@Style["Mitotic Gene Conversion, γ", 18]}, {Left, Bottom}]
```

Out[ ]=

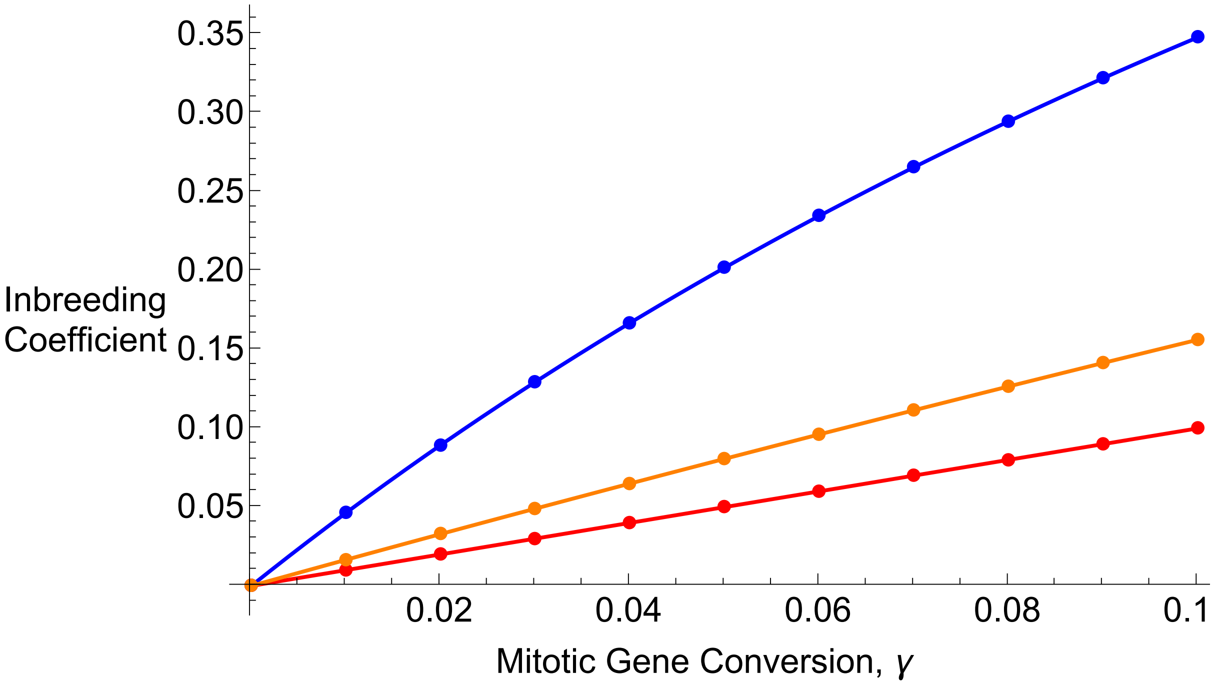


In[ ]:=
```
Export["Figures_Jan2025/RecursionsAsex/RecPlotpA_h05_s02.pdf", plotpA4];
Export["Figures_Jan2025/RecursionsAsex/RecPlotpA_h05_s02.jpg", plotpA4];

Export["Figures_Jan2025/RecursionsAsex/RecPlotF_h05_s02.pdf", plotF4];
Export["Figures_Jan2025/RecursionsAsex/RecPlotF_h05_s02.jpg", plotF4];
```

#### Numerical recursions, self-fertilisation

Below is the function for a change in genotype frequencies over a single generation:

In[ ]:=
```
RecurGCS[{gaa_, gAa_, gAA_}] :=
  {-(((-1 + μ) (4 gaa^2 (1 + (-1 + γ) μ) + gAa (-1 + h s) (gAA (-1 + s)
              (2 γ + S (-1 + γ) (-1 + μ)) + gAa (-1 + h s) (1 + γ - μ + γ μ)) +
          gaa (-4 gAA (-1 + s) (γ + S (-1 + γ) (-1 + μ)) - gAa (-1 + h s)
              (S (-1 + γ) (-1 + μ) + 2 (2 + γ - 2 μ + 2 γ μ))))) /
      (4 (gaa + gAa + gAA - gAA s - gAa h s)^2)), ((-1 + γ) (-1 + μ)
      (4 gaa^2 μ + gAa (-1 + h s) (gAA (-1 + s) (2 + S (-1 + μ)) + gAa (-1 + h s) (1 + μ)) +
        gaa (-4 gAA (-1 + s) (1 + S (-1 + μ)) - gAa (-1 + h s) (2 + S (-1 + μ) + 4 μ)))) /
    (2 (gaa + gAa + gAA - gAA s - gAa h s)^2),
   -((-4 gAA^2 (-1 + s)^2 - 4 gaa gAA (-1 + s) (γ (-1 + μ) + S (-1 + γ) (-1 + μ)^2 - 2 μ) +
        4 gaa^2 (γ (-1 + μ) - μ) μ + gAa^2 (-1 + h s)^2 (-1 + γ (-1 + μ) - μ) (1 + μ) +
        gAa (-1 + h s) (gAA (-1 + s) (2 γ (-1 + μ) + S (-1 + γ) (-1 + μ)^2 - 4 (1 + μ)) +
           gaa (-S (-1 + γ) (-1 + μ)^2 + 4 μ (1 + μ) + 2 γ (1 + μ - 2 μ^2)))) /
      (4 (gaa + gAa + gAA - gAA s - gAa h s)^2))};
```

Defining range of S to be evaluated over:

In[ ]:=
```
Sit = {0, 0.5, 0.99};
```

#### h = 0.01, s = 0.02 plots

In[ ]:=
```
Clear[freqAa, μ, s, h, RecTime]
freqAa = 0.001;
μ  =  10^-8;
s = 0.02;
h = 0.01;
RecTime = 50 000;
```

In[ ]:=
```
Clear[Vrin];
S = Sit[[1]];
Vrin = Table[Clear[RecurRes, γ];
   γ = γit[[j + 1]];
   RecurRes = NestList[RecurGCS, {1 - freqAa, freqAa, 0}, RecTime] // N;
   {γ, RecurRes[[RecTime]][[3]] + RecurRes[[RecTime]][[2]] / 2,
    (RecurRes[[RecTime]][[3]] * RecurRes[[RecTime]][[1]] - (RecurRes[[RecTime]][[2]] / 2)^2) /
     ((RecurRes[[RecTime]][[3]] + RecurRes[[RecTime]][[2]] / 2)
       (RecurRes[[RecTime]][[1]] + RecurRes[[RecTime]][[2]] / 2))}, {j, 0, 10}];
pArecS0 = Partition[Drop[Flatten[Vrin], {3, Length[Flatten[Vrin]], 3}], 2];
FrecS0 = Partition[Drop[Flatten[Vrin], {2, Length[Flatten[Vrin]] - 1, 3}], 2];

Clear[Vrin];
S = Sit[[2]];
Vrin = Table[Clear[RecurRes, γ];
   γ = γit[[j + 1]];
   RecurRes = NestList[RecurGCS, {1 - freqAa, freqAa, 0}, RecTime] // N;
   {γ, RecurRes[[RecTime]][[3]] + RecurRes[[RecTime]][[2]] / 2,
    (RecurRes[[RecTime]][[3]] * RecurRes[[RecTime]][[1]] - (RecurRes[[RecTime]][[2]] / 2)^2) /
     ((RecurRes[[RecTime]][[3]] + RecurRes[[RecTime]][[2]] / 2)
       (RecurRes[[RecTime]][[1]] + RecurRes[[RecTime]][[2]] / 2))}, {j, 0, 10}];
pArecS05 = Partition[Drop[Flatten[Vrin], {3, Length[Flatten[Vrin]], 3}], 2];
FrecS05 = Partition[Drop[Flatten[Vrin], {2, Length[Flatten[Vrin]] - 1, 3}], 2];

Clear[Vrin];
S = Sit[[3]];
Vrin = Table[Clear[RecurRes, γ];
   γ = γit[[j + 1]];
   RecurRes = NestList[RecurGCS, {1 - freqAa, freqAa, 0}, RecTime] // N;
   {γ, RecurRes[[RecTime]][[3]] + RecurRes[[RecTime]][[2]] / 2,
    (RecurRes[[RecTime]][[3]] * RecurRes[[RecTime]][[1]] - (RecurRes[[RecTime]][[2]] / 2)^2) /
     ((RecurRes[[RecTime]][[3]] + RecurRes[[RecTime]][[2]] / 2)
       (RecurRes[[RecTime]][[1]] + RecurRes[[RecTime]][[2]] / 2))}, {j, 0, 10}];
pArecS099 = Partition[Drop[Flatten[Vrin], {3, Length[Flatten[Vrin]], 3}], 2];
FrecS099 = Partition[Drop[Flatten[Vrin], {2, Length[Flatten[Vrin]] - 1, 3}], 2];
```

In[•]:=
```
p1S0 = ListPlot[pArecS0, PlotStyle → {Red, PointSize[Large]}, PlotLegends →
    Placed[PointLegend[{Directive[Red, AbsolutePointSize[8]], Directive[Orange,
        AbsolutePointSize[8]], Directive[Blue, AbsolutePointSize[8]]},
      {"S = 0", "S = 0.5", "S = 0.99"}], {0.4, 0.7}], LabelStyle → 18];
p1S05 = ListPlot[pArecS05, PlotStyle → {Orange, PointSize[Large]},
   PlotLegends → Placed[PointLegend[{Directive[Black, AbsolutePointSize[8]]},
      {"Numerical Recursions"}], {0.8, 0.8}], LabelStyle → 18];
p1S099 = ListPlot[pArecS099, PlotStyle → {Blue, PointSize[Large]}];

p2S0 = Plot[{pAnumSelf[μ, s, h, 0, γin]},
   {γin, 0, 0.1}, PlotRange → Full, PlotStyle → {{Red}}, PlotLegends →
    Placed[LineLegend[{Black}, {"Equation 1"}, LabelStyle → 18], {0.75, 0.65}]];
p2S05 = Plot[{pAnumSelf[μ, s, h, 0.5, γin]},
   {γin, 0, 0.1}, PlotRange → Full, PlotStyle → {{Orange}}];
p2S099 = Plot[{pAnumSelf[μ, s, h, 0.99, γin]},
   {γin, 0, 0.1}, PlotRange → Full, PlotStyle → {{Blue}}];
```

In[•]:=
```
plotpS1 = Labeled[Show[p1S0, p2S0, p1S05, p2S05, p1S099, p2S099, ImageSize → 550,
   LabelStyle → {FontFamily → "Arial", FontSize → 18, FontColor → Black}],
  {Text@TraditionalForm@Style["Frequency\nOf Deleterious\nAllele", 18], Text@
    TraditionalForm@Style["Mitotic Gene Conversion, γ", 18]}, {Left, Bottom}]
```

Out[•]=

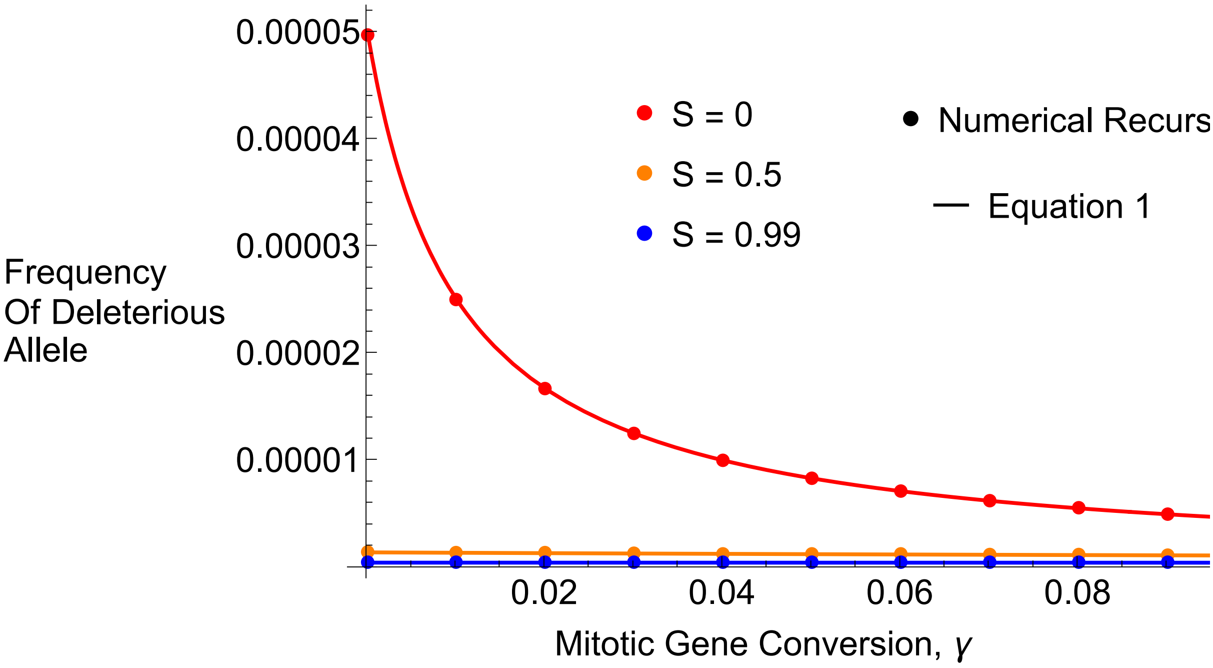


In[•]:=
```
Export["Figures_Jan2025/RecursionsSelf/RecPlotpA_h001_s002_Self.pdf", plotpS1];
Export["Figures_Jan2025/RecursionsSelf/RecPlotpA_h001_s002_Self.jpg", plotpS1];
```

In[∘]:=
```
p1FS0 = ListPlot[FrecS0, PlotStyle → {Red, PointSize[Large]}];
p1FS05 = ListPlot[FrecS05, PlotStyle → {Orange, PointSize[Large]}];
p1FS099 = ListPlot[FrecS099, PlotStyle → {Blue, PointSize[Large]}];

p2FS0 = Plot[{FnumSelf[s, 0, γin, 0.01]}, {γin, 0, 0.1},
   PlotRange → {Full, {-0.05, 1}}, PlotStyle → {{Red}}, PlotLegends →
    Placed[LineLegend[{Black}, {"Equation 3"}, LabelStyle → 18], {0.75, 0.65}]];
p2FS05 = Plot[{FnumSelf[s, 0.5, γin, 0.01]}, {γin, 0, 0.1},
   PlotRange → {Full, {-0.05, 1}}, PlotStyle → {{Orange}}];
p2FS099 = Plot[{FnumSelf[s, 0.99, γin, 0.01]},
   {γin, 0, 0.1}, PlotRange → {Full, {-0.05, 1}}, PlotStyle → {{Blue}}];
```

In[∘]:=
```
plotF1S =
 Labeled[Show[p2FS099, p1FS099, p1FS0, p2FS0, p1FS05, p2FS05, ImageSize → 550,
   LabelStyle → {FontFamily → "Arial", FontSize → 18, FontColor → Black}],
  {Text@TraditionalForm@Style["Inbreeding\nCoefficient", 18], Text@
    TraditionalForm@Style["Mitotic Gene Conversion, γ", 18]}, {Left, Bottom}]
```

Out[∘]=

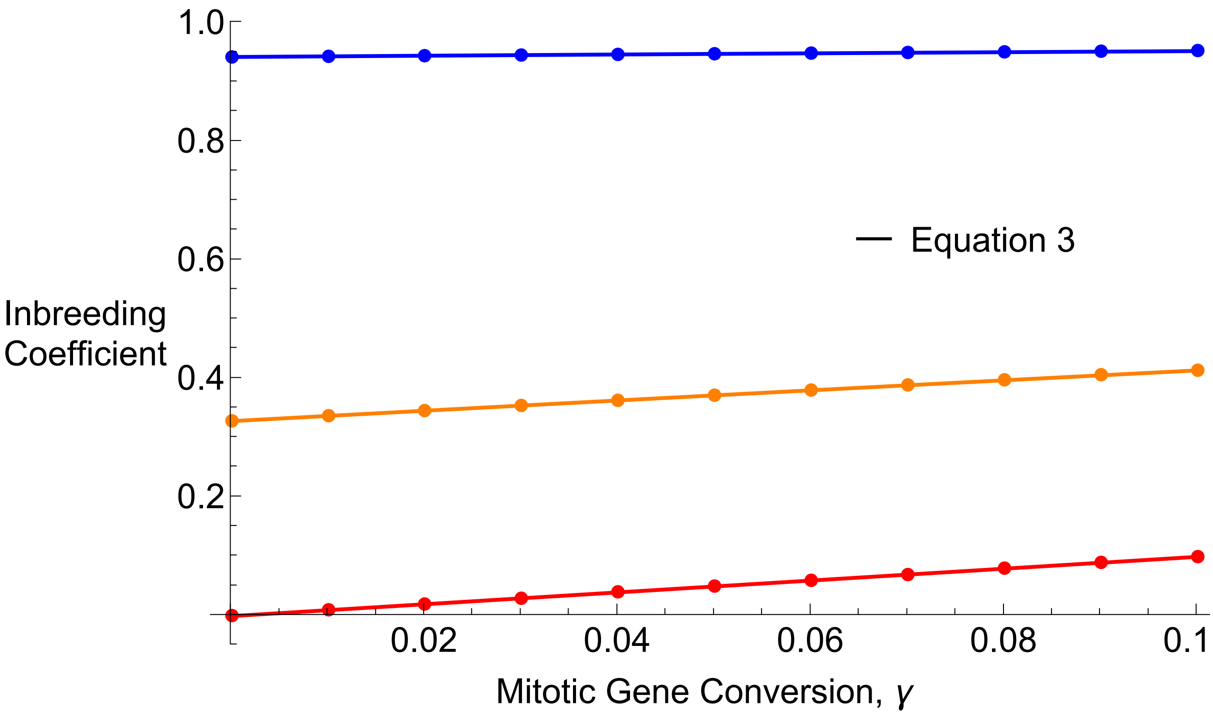


In[∘]:=
```
Export["Figures_Jan2025/RecursionsSelf/RecPlotF_h001_s002_Self.pdf", plotF1S];
Export["Figures_Jan2025/RecursionsSelf/RecPlotF_h001_s002_Self.jpg", plotF1S];
```

#### h = 0.5, s = 0.02 plots

In[∘]:=
```
Clear[freqAa, μ, s, h, RecTime]
freqAa = 0.001;
μ = 10^-8;
s = 0.02;
h = 0.5;
RecTime = 50 000;
```

```
In[•]:= Clear[Vrin];
S = Sit[[1]];
Vrin = Table[Clear[RecurRes, γ];
   γ = γit[[j + 1]];
   RecurRes = NestList[RecurGCS, {1 - freqAa, freqAa, 0}, RecTime] // N;
   {γ, RecurRes[[RecTime]][[3]] + RecurRes[[RecTime]][[2]] / 2,
    (RecurRes[[RecTime]][[3]] * RecurRes[[RecTime]][[1]] - (RecurRes[[RecTime]][[2]] / 2)^2) /
     ((RecurRes[[RecTime]][[3]] + RecurRes[[RecTime]][[2]] / 2)
       (RecurRes[[RecTime]][[1]] + RecurRes[[RecTime]][[2]] / 2))}, {j, 0, 10}];
pA2recS0 = Partition[Drop[Flatten[Vrin], {3, Length[Flatten[Vrin]], 3}], 2];
F2recS0 = Partition[Drop[Flatten[Vrin], {2, Length[Flatten[Vrin]] - 1, 3}], 2];

Clear[Vrin];
S = Sit[[2]];
Vrin = Table[Clear[RecurRes, γ];
   γ = γit[[j + 1]];
   RecurRes = NestList[RecurGCS, {1 - freqAa, freqAa, 0}, RecTime] // N;
   {γ, RecurRes[[RecTime]][[3]] + RecurRes[[RecTime]][[2]] / 2,
    (RecurRes[[RecTime]][[3]] * RecurRes[[RecTime]][[1]] - (RecurRes[[RecTime]][[2]] / 2)^2) /
     ((RecurRes[[RecTime]][[3]] + RecurRes[[RecTime]][[2]] / 2)
       (RecurRes[[RecTime]][[1]] + RecurRes[[RecTime]][[2]] / 2))}, {j, 0, 10}];
pA2recS05 = Partition[Drop[Flatten[Vrin], {3, Length[Flatten[Vrin]], 3}], 2];
F2recS05 = Partition[Drop[Flatten[Vrin], {2, Length[Flatten[Vrin]] - 1, 3}], 2];

Clear[Vrin];
S = Sit[[3]];
Vrin = Table[Clear[RecurRes, γ];
   γ = γit[[j + 1]];
   RecurRes = NestList[RecurGCS, {1 - freqAa, freqAa, 0}, RecTime] // N;
   {γ, RecurRes[[RecTime]][[3]] + RecurRes[[RecTime]][[2]] / 2,
    (RecurRes[[RecTime]][[3]] * RecurRes[[RecTime]][[1]] - (RecurRes[[RecTime]][[2]] / 2)^2) /
     ((RecurRes[[RecTime]][[3]] + RecurRes[[RecTime]][[2]] / 2)
       (RecurRes[[RecTime]][[1]] + RecurRes[[RecTime]][[2]] / 2))}, {j, 0, 10}];
pA2recS099 = Partition[Drop[Flatten[Vrin], {3, Length[Flatten[Vrin]], 3}], 2];
F2recS099 = Partition[Drop[Flatten[Vrin], {2, Length[Flatten[Vrin]] - 1, 3}], 2];
```

In[ ]:=
```
p3S0 = ListPlot[pA2recS0, PlotStyle → {Red, PointSize[Large]}];
p3S05 = ListPlot[pA2recS05, PlotStyle → {Orange, PointSize[Large]}];
p3S099 = ListPlot[pA2recS099, PlotStyle → {Blue, PointSize[Large]}];

p4S0 = Plot[{pAnumSelf[μ, s, h, 0, γin]}, {γin, 0, 0.1},
   PlotRange → {Full, {0, μ/(h s) (1.05)}}, PlotStyle → {{Red}}];
p4S05 = Plot[{pAnumSelf[μ, s, h, 0.5, γin]}, {γin, 0, 0.1},
   PlotRange → {Full, {0, μ/(h s) (1.05)}}, PlotStyle → {{Orange}}];
p4S099 = Plot[{pAnumSelf[μ, s, h, 0.99, γin]}, {γin, 0, 0.1},
   PlotRange → {Full, {0, μ/(h s) (1.05)}}, PlotStyle → {{Blue}}];
```

In[ ]:=
```
plotpS2 = Labeled[Show[p4S0, p3S0, p3S05, p4S05, p4S099, p3S099, ImageSize → 550,
   LabelStyle → {FontFamily → "Arial", FontSize → 18, FontColor → Black}],
  {Text@TraditionalForm@Style["Frequency\nOf Deleterious\nAllele", 18], Text@
    TraditionalForm@Style["Mitotic Gene Conversion, γ", 18]}, {Left, Bottom}]
```

Out[ ]=

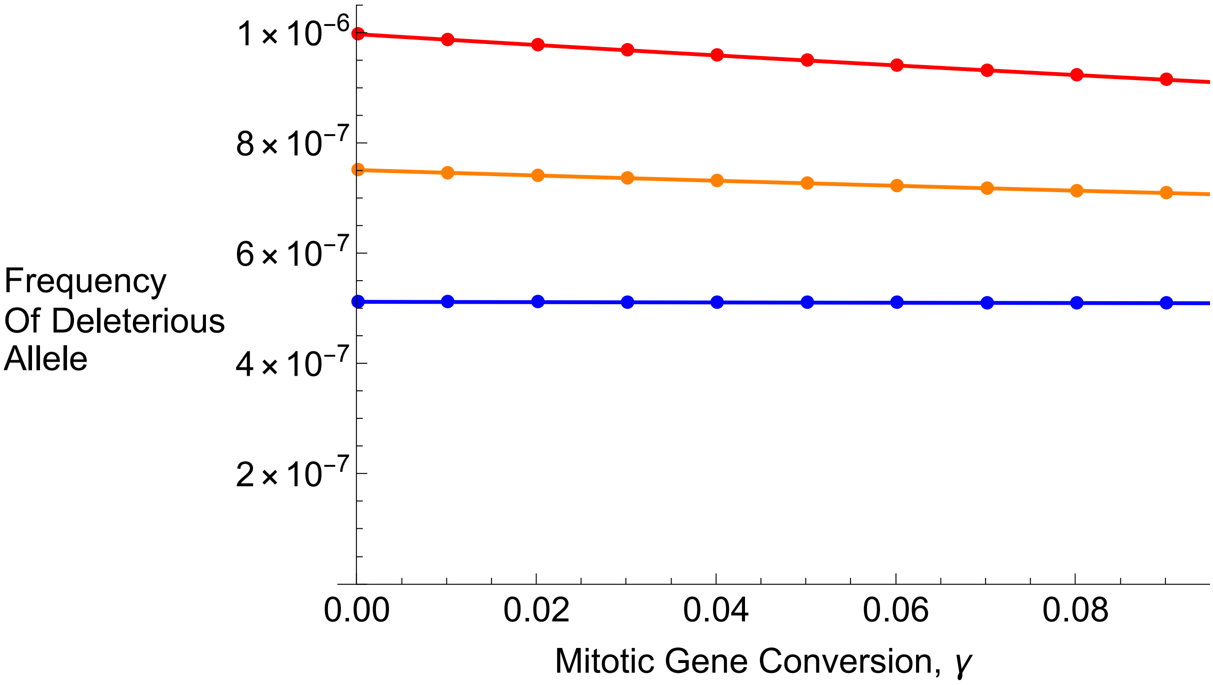


In[ ]:=
```
p3FS0 = ListPlot[F2recS0, PlotStyle → {Red, PointSize[Large]}];
p3FS05 = ListPlot[F2recS05, PlotStyle → {Orange, PointSize[Large]}];
p3FS099 = ListPlot[F2recS099, PlotStyle → {Blue, PointSize[Large]}];

p4FS0 = Plot[{FnumSelf[s, 0, γin, h]}, {γin, 0, 0.1},
   PlotRange → {Full, {-0.05, 1}}, PlotStyle → {{Red}}];
p4FS05 = Plot[{FnumSelf[s, 0.5, γin, h]}, {γin, 0, 0.1},
   PlotRange → {Full, {-0.05, 1}}, PlotStyle → {{Orange}}];
p4FS099 = Plot[{FnumSelf[s, 0.99, γin, h]}, {γin, 0, 0.1},
   PlotRange → {Full, {-0.05, 1}}, PlotStyle → {{Blue}}];
```

In[ ]:=
```
plotF2S =
  Labeled[Show[p4FS099, p3FS099, p3FS0, p4FS0, p3FS05, p4FS05, ImageSize → 550,
    LabelStyle → {FontFamily → "Arial", FontSize → 18, FontColor → Black}],
   {Text@TraditionalForm@Style["Inbreeding\nCoefficient", 18], Text@
     TraditionalForm@Style["Mitotic Gene Conversion, γ", 18]}, {Left, Bottom}]
```

Out[ ]=

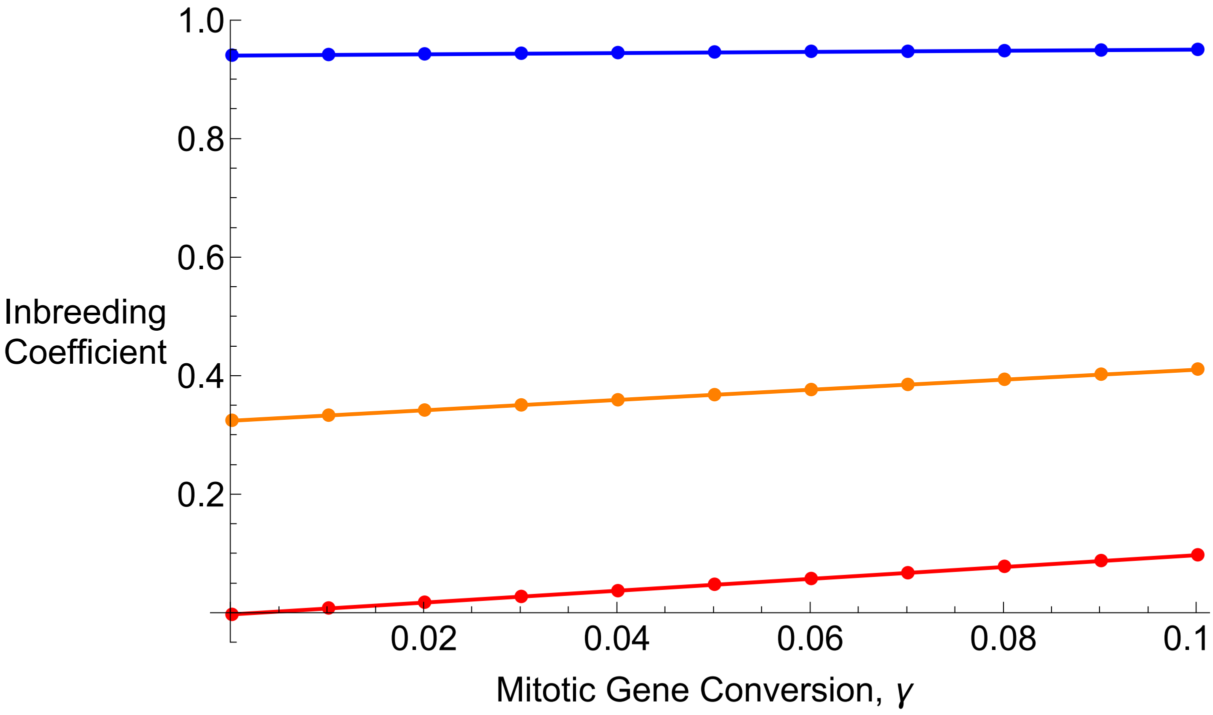


In[ ]:=
```
Export["Figures_Jan2025/RecursionsSelf/RecPlotpA_h05_s002_Self.pdf", plotpS2];
Export["Figures_Jan2025/RecursionsSelf/RecPlotpA_h05_s002_Self.jpg", plotpS2];

Export["Figures_Jan2025/RecursionsSelf/RecPlotF_h05_s002_Self.pdf", plotF2S];
Export["Figures_Jan2025/RecursionsSelf/RecPlotF_h05_s002_Self.jpg", plotF2S];
```

#### h = 0.01, s = 0.2 plots

In[ ]:=
```
Clear[freqAa, μ, s, h, RecTime]
freqAa = 0.001;
μ = 10^-8;
s = 0.2;
h = 0.01;
RecTime = 50 000;
```

In[ ]:=
```
Clear[Vrin];
S = Sit⟦1⟧;
Vrin = Table[Clear[RecurRes, γ];
   γ = γit⟦j + 1⟧;
   RecurRes = NestList[RecurGCS, {1 - freqAa, freqAa, 0}, RecTime] // N;
   {γ, RecurRes⟦RecTime⟧⟦3⟧ + RecurRes⟦RecTime⟧⟦2⟧ / 2,
    (RecurRes⟦RecTime⟧⟦3⟧ * RecurRes⟦RecTime⟧⟦1⟧ - (RecurRes⟦RecTime⟧⟦2⟧ / 2)^2) /
     ((RecurRes⟦RecTime⟧⟦3⟧ + RecurRes⟦RecTime⟧⟦2⟧ / 2)
       (RecurRes⟦RecTime⟧⟦1⟧ + RecurRes⟦RecTime⟧⟦2⟧ / 2))}, {j, 0, 10}];
pA3recS0 = Partition[Drop[Flatten[Vrin], {3, Length[Flatten[Vrin]], 3}], 2];
F3recS0 = Partition[Drop[Flatten[Vrin], {2, Length[Flatten[Vrin]] - 1, 3}], 2];

Clear[Vrin];
S = Sit⟦2⟧;
Vrin = Table[Clear[RecurRes, γ];
   γ = γit⟦j + 1⟧;
   RecurRes = NestList[RecurGCS, {1 - freqAa, freqAa, 0}, RecTime] // N;
   {γ, RecurRes⟦RecTime⟧⟦3⟧ + RecurRes⟦RecTime⟧⟦2⟧ / 2,
    (RecurRes⟦RecTime⟧⟦3⟧ * RecurRes⟦RecTime⟧⟦1⟧ - (RecurRes⟦RecTime⟧⟦2⟧ / 2)^2) /
     ((RecurRes⟦RecTime⟧⟦3⟧ + RecurRes⟦RecTime⟧⟦2⟧ / 2)
       (RecurRes⟦RecTime⟧⟦1⟧ + RecurRes⟦RecTime⟧⟦2⟧ / 2))}, {j, 0, 10}];
pA3recS05 = Partition[Drop[Flatten[Vrin], {3, Length[Flatten[Vrin]], 3}], 2];
F3recS05 = Partition[Drop[Flatten[Vrin], {2, Length[Flatten[Vrin]] - 1, 3}], 2];

Clear[Vrin];
S = Sit⟦3⟧;
Vrin = Table[Clear[RecurRes, γ];
   γ = γit⟦j + 1⟧;
   RecurRes = NestList[RecurGCS, {1 - freqAa, freqAa, 0}, RecTime] // N;
   {γ, RecurRes⟦RecTime⟧⟦3⟧ + RecurRes⟦RecTime⟧⟦2⟧ / 2,
    (RecurRes⟦RecTime⟧⟦3⟧ * RecurRes⟦RecTime⟧⟦1⟧ - (RecurRes⟦RecTime⟧⟦2⟧ / 2)^2) /
     ((RecurRes⟦RecTime⟧⟦3⟧ + RecurRes⟦RecTime⟧⟦2⟧ / 2)
       (RecurRes⟦RecTime⟧⟦1⟧ + RecurRes⟦RecTime⟧⟦2⟧ / 2))}, {j, 0, 10}];
pA3recS099 = Partition[Drop[Flatten[Vrin], {3, Length[Flatten[Vrin]], 3}], 2];
F3recS099 = Partition[Drop[Flatten[Vrin], {2, Length[Flatten[Vrin]] - 1, 3}], 2];
```

```
In[ ]:= p5S0 = ListPlot[pA3recS0, PlotStyle → {Red, PointSize[Large]}];
p5S05 = ListPlot[pA3recS05, PlotStyle → {Orange, PointSize[Large]}];
p5S099 = ListPlot[pA3recS099, PlotStyle → {Blue, PointSize[Large]}];

p6S0 = Plot[{pAnumSelf[μ, s, h, 0, γin]}, {γin, 0, 0.1},
   PlotRange → {Full, {0, μ/(h s) (1.05)}}, PlotStyle → {{Red}}];
p6S05 = Plot[{pAnumSelf[μ, s, h, 0.5, γin]}, {γin, 0, 0.1},
   PlotRange → {Full, {0, μ/(h s) (1.05)}}, PlotStyle → {{Orange}}];
p6S099 = Plot[{pAnumSelf[μ, s, h, 0.99, γin]}, {γin, 0, 0.1},
   PlotRange → {Full, {0, μ/(h s) (1.05)}}, PlotStyle → {{Blue}}];

In[ ]:= plotpS3 = Labeled[Show[p6S099, p5S099, p5S05, p6S05, p5S0, p6S0, ImageSize → 550,
   LabelStyle → {FontFamily → "Arial", FontSize → 18, FontColor → Black}],
  {Text@TraditionalForm@Style["Frequency\nOf Deleterious\nAllele", 18], Text@
    TraditionalForm@Style["Mitotic Gene Conversion, γ", 18]}, {Left, Bottom}]
```

Out[ ]=

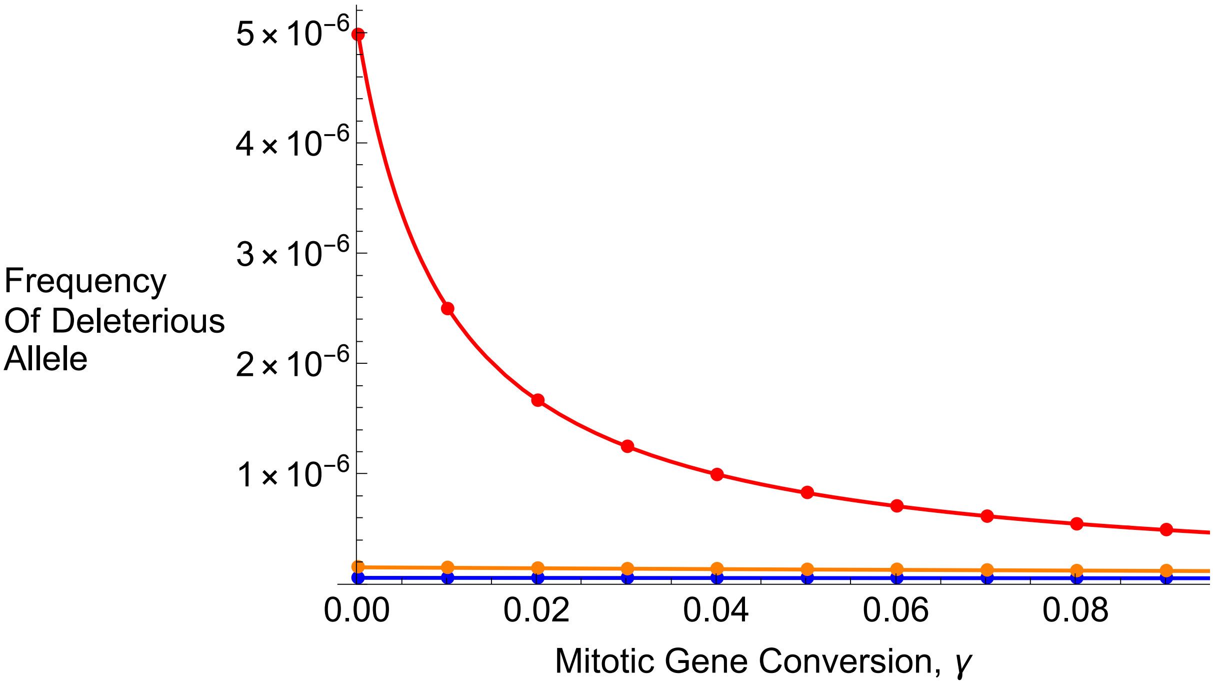


```
In[ ]:= p5FS0 = ListPlot[F3recS0, PlotStyle → {Red, PointSize[Large]}];
p5FS05 = ListPlot[F3recS05, PlotStyle → {Orange, PointSize[Large]}];
p5FS099 = ListPlot[F3recS099, PlotStyle → {Blue, PointSize[Large]}];

p6FS0 = Plot[{FnumSelf[s, 0, γin, h]}, {γin, 0, 0.1},
   PlotRange → {Full, {-0.05, 1}}, PlotStyle → {{Red}, {Red, Dashed}}];
p6FS05 = Plot[{FnumSelf[s, 0.5, γin, h]}, {γin, 0, 0.1},
   PlotRange → {Full, {-0.05, 1}}, PlotStyle → {{Orange}, {Orange, Dashed}}];
p6FS099 = Plot[{FnumSelf[s, 0.99, γin, h]}, {γin, 0, 0.1},
   PlotRange → {Full, {-0.05, 1}}, PlotStyle → {{Blue}, {Blue, Dashed}}];
```

In[∘]:=
```
plotF3S =
  Labeled[Show[p6FS099, p5FS099, p5FS0, p6FS0, p5FS05, p6FS05, ImageSize → 550,
    LabelStyle → {FontFamily → "Arial", FontSize → 18, FontColor → Black}],
   {Text@TraditionalForm@Style["Inbreeding\nCoefficient", 18], Text@
     TraditionalForm@Style["Mitotic Gene Conversion, γ", 18]}, {Left, Bottom}]
```

Out[∘]=

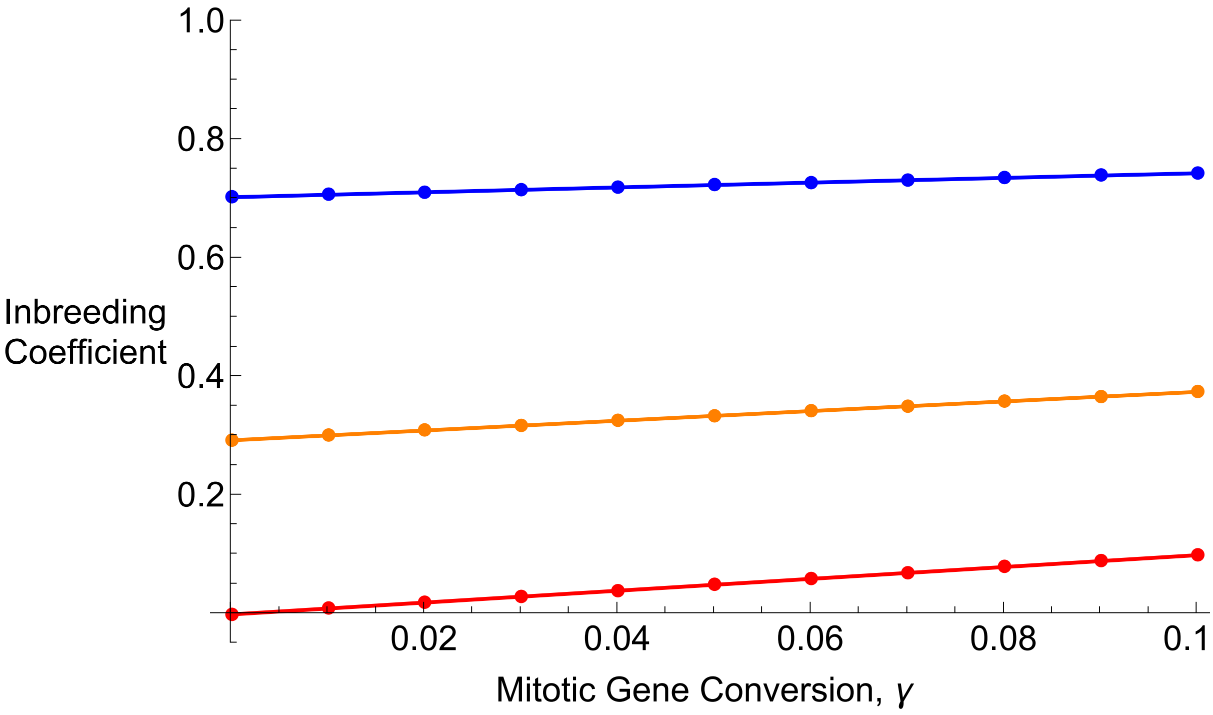


In[∘]:=
```
Export["Figures_Jan2025/RecursionsSelf/RecPlotpA_h001_s02_Self.pdf", plotpS3];
Export["Figures_Jan2025/RecursionsSelf/RecPlotpA_h001_s02_Self.jpg", plotpS3];

Export["Figures_Jan2025/RecursionsSelf/RecPlotF_h001_s02_Self.pdf", plotF3S];
Export["Figures_Jan2025/RecursionsSelf/RecPlotF_h001_s02_Self.jpg", plotF3S];
```

#### h = 0.5, s = 0.2 plots

In[∘]:=
```
Clear[freqAa, μ, s, h, RecTime]
freqAa = 0.001;
μ = 10^-8;
s = 0.2;
h = 0.5;
RecTime = 50 000;
```

```
In[•]:= Clear[Vrin];
S = Sit[[1]];
Vrin = Table[Clear[RecurRes, γ];
   γ = γit[[j + 1]];
   RecurRes = NestList[RecurGCS, {1 - freqAa, freqAa, 0}, RecTime] // N;
   {γ, RecurRes[[RecTime]][[3]] + RecurRes[[RecTime]][[2]] / 2,
    (RecurRes[[RecTime]][[3]] * RecurRes[[RecTime]][[1]] - (RecurRes[[RecTime]][[2]] / 2)^2) /
     ((RecurRes[[RecTime]][[3]] + RecurRes[[RecTime]][[2]] / 2)
       (RecurRes[[RecTime]][[1]] + RecurRes[[RecTime]][[2]] / 2))}, {j, 0, 10}];
pA4recS0 = Partition[Drop[Flatten[Vrin], {3, Length[Flatten[Vrin]], 3}], 2];
F4recS0 = Partition[Drop[Flatten[Vrin], {2, Length[Flatten[Vrin]] - 1, 3}], 2];

Clear[Vrin];
S = Sit[[2]];
Vrin = Table[Clear[RecurRes, γ];
   γ = γit[[j + 1]];
   RecurRes = NestList[RecurGCS, {1 - freqAa, freqAa, 0}, RecTime] // N;
   {γ, RecurRes[[RecTime]][[3]] + RecurRes[[RecTime]][[2]] / 2,
    (RecurRes[[RecTime]][[3]] * RecurRes[[RecTime]][[1]] - (RecurRes[[RecTime]][[2]] / 2)^2) /
     ((RecurRes[[RecTime]][[3]] + RecurRes[[RecTime]][[2]] / 2)
       (RecurRes[[RecTime]][[1]] + RecurRes[[RecTime]][[2]] / 2))}, {j, 0, 10}];
pA4recS05 = Partition[Drop[Flatten[Vrin], {3, Length[Flatten[Vrin]], 3}], 2];
F4recS05 = Partition[Drop[Flatten[Vrin], {2, Length[Flatten[Vrin]] - 1, 3}], 2];

Clear[Vrin];
S = Sit[[3]];
Vrin = Table[Clear[RecurRes, γ];
   γ = γit[[j + 1]];
   RecurRes = NestList[RecurGCS, {1 - freqAa, freqAa, 0}, RecTime] // N;
   {γ, RecurRes[[RecTime]][[3]] + RecurRes[[RecTime]][[2]] / 2,
    (RecurRes[[RecTime]][[3]] * RecurRes[[RecTime]][[1]] - (RecurRes[[RecTime]][[2]] / 2)^2) /
     ((RecurRes[[RecTime]][[3]] + RecurRes[[RecTime]][[2]] / 2)
       (RecurRes[[RecTime]][[1]] + RecurRes[[RecTime]][[2]] / 2))}, {j, 0, 10}];
pA4recS099 = Partition[Drop[Flatten[Vrin], {3, Length[Flatten[Vrin]], 3}], 2];
F4recS099 = Partition[Drop[Flatten[Vrin], {2, Length[Flatten[Vrin]] - 1, 3}], 2];
```

```
In[•]:= p7S0 = ListPlot[pA4recS0, PlotStyle → {Red, PointSize[Large]}];
p7S05 = ListPlot[pA4recS05, PlotStyle → {Orange, PointSize[Large]}];
p7S099 = ListPlot[pA4recS099, PlotStyle → {Blue, PointSize[Large]}];

p8S0 = Plot[{pAnumSelf[μ, s, h, 0, γin]}, {γin, 0, 0.1},
   PlotRange → {Full, {0, μ/(h s) (1.05)}}, PlotStyle → {{Red}, {Red, Dashed}}];
p8S05 = Plot[{pAnumSelf[μ, s, h, 0.5, γin]}, {γin, 0, 0.1},
   PlotRange → {Full, {0, μ/(h s) (1.05)}}, PlotStyle → {{Orange}, {Orange, Dashed}}];
p8S099 = Plot[{pAnumSelf[μ, s, h, 0.99, γin]}, {γin, 0, 0.1},
   PlotRange → {Full, {0, μ/(h s) (1.05)}}, PlotStyle → {{Blue}, {Blue, Dashed}}];
```

```
In[•]:= plotpS4 = Labeled[Show[p8S099, p7S099, p7S05, p8S05, p7S0, p8S0, ImageSize → 550,
    LabelStyle → {FontFamily → "Arial", FontSize → 18, FontColor → Black}],
  {Text@TraditionalForm@Style["Frequency\nOf Deleterious\nAllele", 18], Text@
    TraditionalForm@Style["Mitotic Gene Conversion, γ", 18]}, {Left, Bottom}]
```

Out[•]=

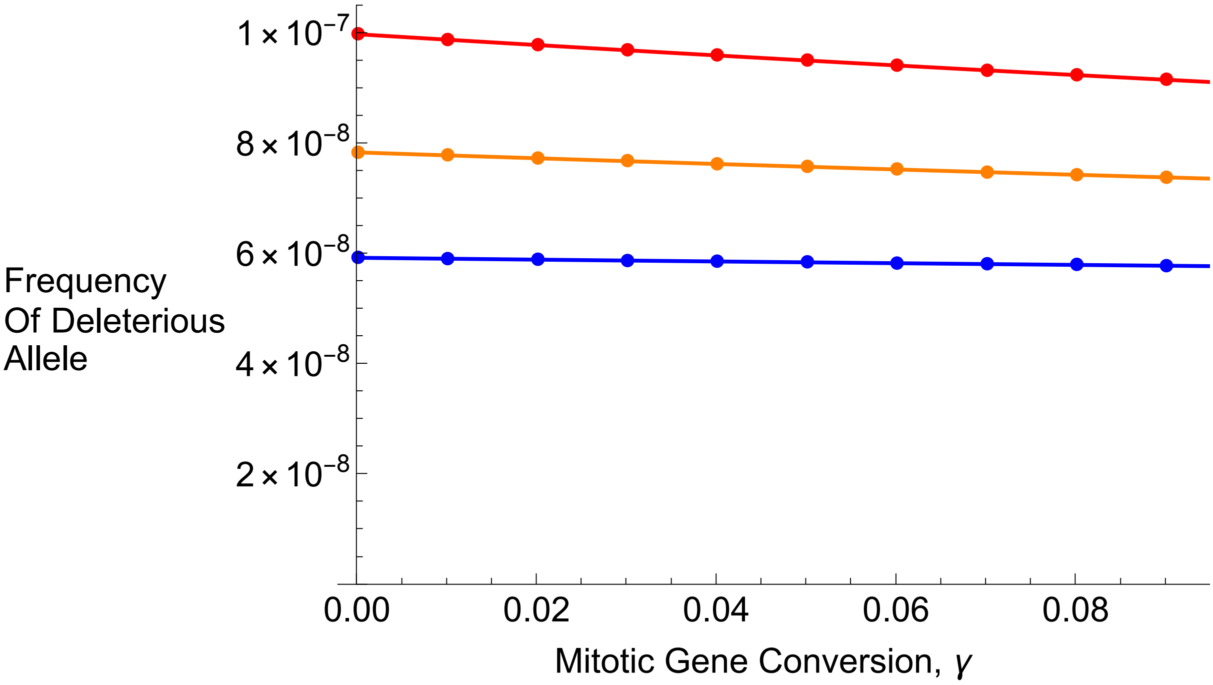


```
In[•]:= p7FS0 = ListPlot[F4recS0, PlotStyle → {Red, PointSize[Large]}];
p7FS05 = ListPlot[F4recS05, PlotStyle → {Orange, PointSize[Large]}];
p7FS099 = ListPlot[F4recS099, PlotStyle → {Blue, PointSize[Large]}];

p8FS0 = Plot[{FnumSelf[s, 0, γin, h]}, {γin, 0, 0.1},
   PlotRange → {Full, {-0.05, 1}}, PlotStyle → {{Red}, {Red, Dashed}}];
p8FS05 = Plot[{FnumSelf[s, 0.5, γin, h]}, {γin, 0, 0.1},
   PlotRange → {Full, {-0.05, 1}}, PlotStyle → {{Orange}, {Orange, Dashed}}];
p8FS099 = Plot[{FnumSelf[s, 0.99, γin, h]}, {γin, 0, 0.1},
   PlotRange → {Full, {-0.05, 1}}, PlotStyle → {{Blue}, {Blue, Dashed}}];
```

In[∘]:=
```
plotF4S =
  Labeled[Show[p8FS099, p7FS099, p7FS0, p8FS0, p7FS05, p8FS05, ImageSize → 550,
    LabelStyle → {FontFamily → "Arial", FontSize → 18, FontColor → Black}],
   {Text@TraditionalForm@Style["Inbreeding\nCoefficient", 18], Text@
     TraditionalForm@Style["Mitotic Gene Conversion, γ", 18]}, {Left, Bottom}]
```

Out[∘]=

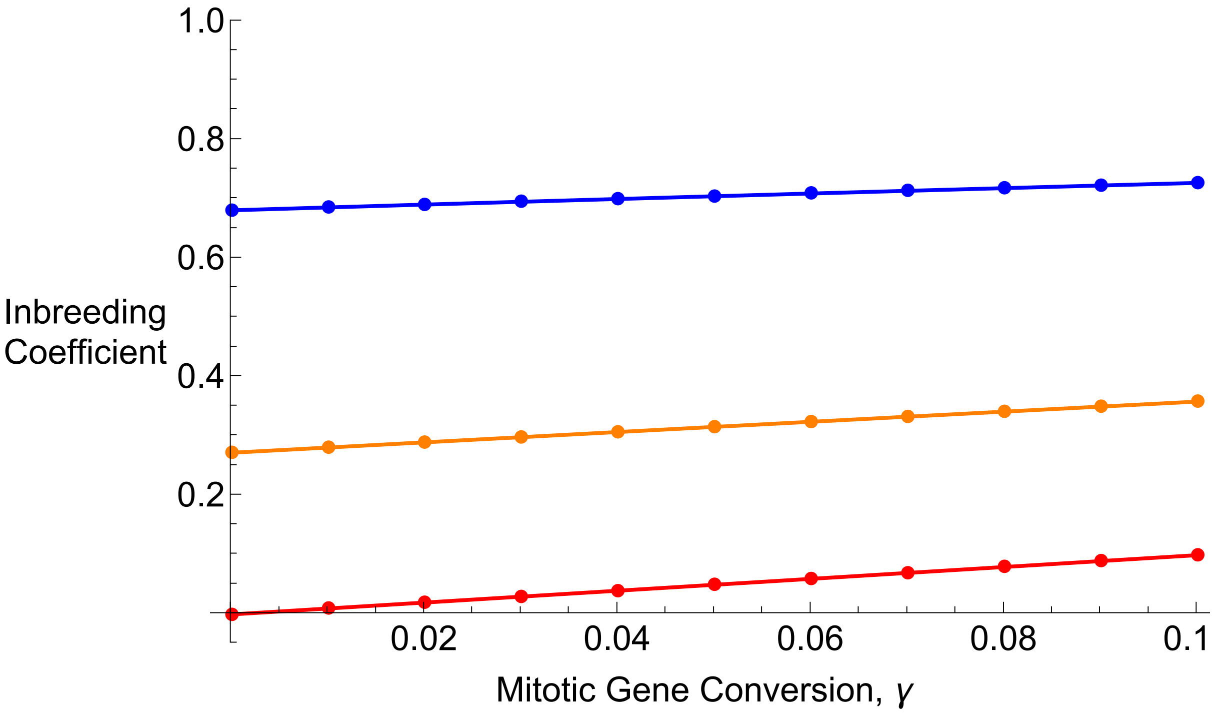


In[∘]:=
```
Export["Figures_Jan2025/RecursionsSelf/RecPlotpA_h05_s02_Self.pdf", plotpS4];
Export["Figures_Jan2025/RecursionsSelf/RecPlotpA_h05_s02_Self.jpg", plotpS4];

Export["Figures_Jan2025/RecursionsSelf/RecPlotF_h05_s02_Self.pdf", plotF4S];
Export["Figures_Jan2025/RecursionsSelf/RecPlotF_h05_s02_Self.jpg", plotF4S];
```

### Section B: Gene conversion lengths, conditioning on non-zero tracts

The gene conversion (GC) routine works as follows:
- A ‘centre’ is defined for the GC event
- Two GC tracts are drawn going left, right from this centre from a geometric distribution with mean $\lambda/2$
- the total sum is hence from a negative binomial distribution.

The PDF of a neg bin distribution(n, p) with n = 1 is the same as a Geometric distribution in *Mathematica*:

In[ ]:= `PDF[NegativeBinomialDistribution[1, p], x]`

Out[ ]=

$$\begin{cases} (1-p)^x p & x \geq 0 \\ 0 & \text{True} \end{cases}$$

In[ ]:= `PDF[GeometricDistribution[p], x]`

Out[ ]=

$$\begin{cases} (1-p)^x p & x \geq 0 \\ 0 & \text{True} \end{cases}$$

And the mean of the geometric distribution is $\frac{1}{p}-1$, so one is drawing for the number of *failures* before a success. This is the same definition that SLiM uses.

In[ ]:= `Mean[GeometricDistribution[p]]`

Out[ ]=

$$-1+\frac{1}{p}$$

So the total length is drawn from NegBin(2,p).

We want to know:
1) What is the distribution, *conditional on rejecting zero-length tracts?*
2) What is the *mean* of this conditional distribution?
3) How do we parameterize it in the simulations?

To answer (1), we use Bayes' theorem:

$$P(X=x|X!=0) = \frac{P(X\neq 0\,|X=x)\,P(X=x)}{P(X\neq 0)}$$

P(X!=0|X=x) = 1 by logic.
P(X=x) is the negative binomial distribution, NB(2,p):

In[ ]:= `PDF[NegativeBinomialDistribution[2, p], x]`

Out[ ]=

$$\begin{cases} (1-p)^x p^2 (1+x) & x \geq 0 \\ 0 & \text{True} \end{cases}$$

P(X!=0) can be calculated from the distribution as 1-P(X=0)

In[ ]:= `1 - PDF[NegativeBinomialDistribution[2, p], 0]`

Out[ ]=

$$1-p^2$$

So the conditional distribution is simply:

In[ ]:=

$$\frac{(1-p)^x p^2 (1+x)}{1-p^2}$$

Out[ ]=

$$\frac{(1-p)^x p^2 (1+x)}{1-p^2}$$

Checking that it is valid (i.e. it sums to 1)

In[ ]:= 
```
Sum[((1 - p)^x p^2 (1 + x))/(1 - p^2), {x, 1, ∞}] // Simplify
```
Out[ ]=

$$1$$

(2) What is the mean? It is simply this sum:

In[ ]:= 
```
Sum[x ((1 - p)^x p^2 (1 + x))/(1 - p^2), {x, 1, ∞}]
```

$$\frac{2}{p\ (1+p)}$$

(3) How do we parameterize this distribution?

Let y be the mean value we want to set our simulation to (e.g. 2000bp, denoted $\lambda$ in the manuscript). Then $y = \frac{2}{p\ (1+p)}$.

Solving for p:

In[ ]:= 
```
Solve[y == 2/(p (1 + p)), p]
```
Out[ ]=

$$\left\{\left\{p \to \frac{-y - \sqrt{y}\ \sqrt{8+y}}{2\ y}\right\}, \left\{p \to \frac{-y + \sqrt{y}\ \sqrt{8+y}}{2\ y}\right\}\right\}$$

Hence there are two possible solutions of p, but with some inspection it seems that only the latter yield valid values of p between 0 and 1. We can see this, for example, by taking the limit of both solutions between y = 1 (the lowest value) and infinity:

In[ ]:= 
```
Limit[{{(-y - Sqrt[y] Sqrt[8 + y])/(2 y)}, {(-y + Sqrt[y] Sqrt[8 + y])/(2 y)}}, y → 1]
Limit[{{(-y - Sqrt[y] Sqrt[8 + y])/(2 y)}, {(-y + Sqrt[y] Sqrt[8 + y])/(2 y)}}, y → ∞]
```
Out[ ]=

{{-2}, {1}}

Out[ ]=

{{-1}, {0}}

So when the geometric draws are defined in SLiM, if each is parameterized with $p = \frac{-y+\sqrt{y}\ \sqrt{8+y}}{2\ y}$ and zero-length tracts are discarded then the mean of the final distribution will equal y.

We can also calculate the CDF of this conditional distribution:

In[ ]:= 
```
Sum[((1 - p)^x p^2 (1 + x))/(1 - p^2), {x, 1, z}] // Simplify
```
Out[ ]=

$$-\frac{-1 + (1-p)^z + p\ (-1 + (1-p)^z + (1-p)^z\ z)}{1+p}$$

Then if we substitute $p = \frac{-y+\sqrt{y}\ \sqrt{8+y}}{2\ y}$ to define this CDF with overall mean y:

In[ ]:=
$$-\frac{-1+(1-p)^z+p\,(-1+(1-p)^z+(1-p)^z\,z)}{1+p}\ /.\ p\to\frac{-y+\sqrt{y}\,\sqrt{8+y}}{2\,y}\ //\ \text{Simplify}$$

Out[ ]=
$$\frac{\sqrt{y}\left(1-\left(\frac{3}{2}-\frac{1}{2\sqrt{\frac{y}{8+y}}}\right)^z+\left(\frac{3}{2}-\frac{1}{2\sqrt{\frac{y}{8+y}}}\right)^z z\right)-\sqrt{8+y}\left(-1+\left(\frac{3}{2}-\frac{1}{2\sqrt{\frac{y}{8+y}}}\right)^z+\left(\frac{3}{2}-\frac{1}{2\sqrt{\frac{y}{8+y}}}\right)^z z\right)}{\sqrt{y}+\sqrt{8+y}}$$

In[ ]:= CDFCon[y_, z_] :=
$$\frac{\sqrt{y}\left(1-\left(\frac{3}{2}-\frac{1}{2\sqrt{\frac{y}{8+y}}}\right)^z+\left(\frac{3}{2}-\frac{1}{2\sqrt{\frac{y}{8+y}}}\right)^z z\right)-\sqrt{8+y}\left(-1+\left(\frac{3}{2}-\frac{1}{2\sqrt{\frac{y}{8+y}}}\right)^z+\left(\frac{3}{2}-\frac{1}{2\sqrt{\frac{y}{8+y}}}\right)^z z\right)}{\sqrt{y}+\sqrt{8+y}}$$

This function gives the probability that a draw will have less then length z, given mean y.
We can numerically solve it to work out what the upper quantile is for a given probability and overall mean.
For example, if y = 2000 then 99% of draws will be less than the following length:

In[ ]:= `NSolve[CDFCon[2000, z] == 0.99, z]`

NSolve: Inverse functions are being used by NSolve, so some solutions may not be found; use Reduce for complete solution information.

Out[ ]= `{{z → -998.305}, {z → 6640.17}}`

Obviously only the positive value is valid here, i.e. 6640.
If you want more events of longer tract length then we can use a larger mean. Letting y = 3000 means that 99% of events are less than the following:

In[ ]:= `NSolve[CDFCon[3000, z] == 0.99, z]`

NSolve: Inverse functions are being used by NSolve, so some solutions may not be found; use Reduce for complete solution information.

Out[ ]= `{{z → -1496.46}, {z → 9959.34}}`

So 1% of events exceed 9959 or nearly 10,000, so this mean might be better to use in the simulations. In this case, p would equal:

In[ ]:=
$$\frac{-y+\sqrt{y}\,\sqrt{8+y}}{2\,y}\ /.\ y\to 3000\ //\ N$$

Out[ ]= `0.000666223`

In[ ]:= `NSolve[CDFCon[4000, z] == 0.95, z]`

NSolve: Inverse functions are being used by NSolve, so some solutions may not be found; use Reduce for complete solution information.

Out[ ]= `{{z → -1964.51}, {z → 9488.6}}`
